# Nup153 regulates neuronal responsiveness through HDAC1-mediated epigenetic modulation

**DOI:** 10.64898/2026.08.02.742238

**Authors:** Abhinav Soni, Stavroula Petridi, Maria Ludovica Sforza, Diana Klütsch, Koki Sakurai, Xin Hu, Nicole Rund, Anne Karasinsky, Julia Hernandez Pineda, Judith Houtman, Enzo Scifo, Mathias Lesche, Katrin Sameith, Andreas Dahl, Christian Rosenmund, Dan Ehninger, Anna R. Poetsch, Jelle van den Ameele, Hayder Amin, Tomohisa Toda

## Abstract

Neural activity-dependent gene regulation is central to the development of neural networks and neuronal plasticity. Induction of activity-dependent gene programs is equally important as repression of these programs, and both need to be balanced carefully. However, little is known about how repressive mechanisms modulate neuronal responsiveness across the genome before stimulation. Here, we identify nucleoporin-dependent regulation of neuronal responsiveness, in which Nup153 represses neuronal genes including activity-regulated genes (ARGs), in the basal state. By characterizing the genome-wide landscape of chromatin accessibility, histone modifications and Nup153 chromatin binding, we show that Nup153 bidirectionally regulates chromatin states through both basal activity-dependent and -independent mechanisms and influences associated genes. Mechanistically, Nup153 associates with HDAC1 to modulate histone acetylation and chromatin accessibility at target regulatory regions. Our data suggests Nup153 organizes chromatin states that regulate neuronal gene programs involved in maintaining neuronal responsiveness.

**Teaser:** Nup153 primes neuronal responsiveness through multi-layered epigenetic regulation.

## INTRODUCTION

Neurons respond to synaptic inputs elicited by sensory input or neural network activity, which drive the activation of intrinsic genetic programs (*1*). This neuronal activity-dependent gene expression is an essential program for nervous system maturation and for learning and memory in the adult brain (*2–4*). To adapt to environmental changes, neurons must maintain neuronal responsiveness, i.e., the ability to drive this program in response to inputs. In order to do so, a delicate balance between activation and repression of the program is crucial. Tremendous efforts have already been made to understand the mechanisms underlying activation, and several signaling pathways and key transcription factors have been identified. The activated programs induce the transcription of activity-regulated genes (ARGs), such as *Fos*, *Egr-1, Npas4*, *Arc,* and *Bdnf* (*5–8*). To induce ARGs, key transcription factors activated by synaptic inputs bind to regulatory genetic elements such as ARG promoters or enhancers, modulate chromatin states and consequently promote ARG transcription (*6, 9–11*). The transient activation of early ARGs further induces downstream gene programs that are tailored to activate neural cell type-specific genetic pathways to regulate cell type-specific neuroplastic functions (*12–15*).

However, in contrast to the mechanisms underlying ARG activation, the mechanisms underlying the repression of neuronal activity-dependent gene programs still remain largely elusive. For neuronal activity-dependent gene programs, it is crucial that ARGs are activated within a few minutes of neuronal activation. At the same time, it is equally important that ARG expression levels are kept low prior to neuronal activation in the basal state. Robust repression of ARGs in the basal state reduces the leaky induction of ARGs, ensuring that neurons can accurately encode the spatiotemporal information conveyed by neuronal activity into ARG expression and downstream genetic programs. Previous studies have shown that histone deacetylases (HDACs) bind to the regulatory elements of ARGs and likely contribute to their repression (*16–18*). However, most studies have used artificially silenced neurons and focused on ARG regulation following stimulation, or investigated the long-term effects of genetic perturbations (*16–19*). Furthermore, although it is known that neuronal activation triggers changes in histone modifications that can increase chromatin accessibility and control the permissiveness of chromatin states for ARG induction (*20*), the mechanisms that enforce a repressive neuronal state prior to neuronal stimulation in the basal state, and those that coordinate repressive factors, including HDACs, on the regulatory elements of ARGs genome-wide are still unclear.

In the present study, we addressed how neuronal responsiveness is maintained in the basal state through the balanced regulation of neuronal genes including key ARGs by focusing on the gene regulatory mechanisms mediated by the nuclear pore protein Nup153. Nup153, a nuclear basket protein, interacts with chromatin and regulates developmental genetic programs in cooperation with transcription factors and epigenetic factors in stem cells (*21–24*). In neural progenitor cells (NeuPCs), Nup153 acts as a structural platform to recruit the key transcription factor Sox2 and maintain the neural progenitor state, and Nup153 levels change dynamically upon neural differentiation. Furthermore, Nup153 binds to chromatin to regulate gene expression in a bimodal manner depending on its binding location (*21*). In embryonic stem cells, Nup153 is essential for maintaining the pluripotent state through gene repression in cooperation with the Polycomb repressive complex 1 (PRC1) (*22*). These results indicate that Nup153-directed nuclear architecture can act as a gatekeeper for maintaining cellular identity by contributing to the organization of cell type-specific nuclear architecture and epigenetic programs. Because nuclear pores are the interface between the cytoplasm and the nucleus, and several critical signals originating from synapses have to go through nuclear pores to find their targets on chromatin in neurons, we hypothesized that Nup153 may also organize neuronal responsiveness through cell type-specific epigenetic regulation in post-mitotic functional neurons.

Here, we investigated the role of Nup153 in neuronal responsiveness through basal activity-dependent and -independent gene regulation in post-mitotic functional neurons. We show that adequate levels of Nup153 are essential for maintaining ARG expression at low levels in the basal state, and loss of Nup153 dysregulated a wide range of neuronal genes, including a subset of key ARGs. Mechanistically, Nup153 binds directly to key regulatory regions of the neuronal genome and bidirectionally regulates chromatin accessibility and histone modification in both an activity-dependent and an activity-independent manner. Furthermore, Nup153 associates with HDAC1 to repress a subset of targets through histone modification, thus controlling the epigenetic state of target genes with its partner. Loss of Nup153 dysregulated neuronal gene expression in the basal state and impaired the balance of neural network activity. Taken together, our results indicate that Nup153 not only is important for lineage-specific gene regulation but also organizes neuronal responsiveness by regulating the epigenetic states of target genes in post-mitotic functional neurons. Our findings highlight that Nup153-directed nuclear architecture contributes to the organization of neuronal states through dynamic epigenetic regulation.

## RESULTS

### Dynamic changes in Nup153 levels in mature cortical neurons

To probe the potential role of Nup153 in post-mitotic neurons, Nup153 levels were assessed in mature primary cortical neurons. A previous study showed that Nup153 is selectively downregulated upon neural differentiation from neural progenitor cells (NeuPCs) (*21*). Primary cortical neurons (PCNs) were cultured and maturated until days *in vitro* (DIV)13. To our surprise, Nup153 levels in Map2-positive neurons were markedly higher than in Sox2-positive NeuPCs and in GFAP-positive astrocytes (NeuPCs vs Map2-positive neurons, Astrocytes vs Map2-positive neurons, p < 0.0001, Mann-Whitney test) (Fig. 1A, 1B), indicating that Nup153 is downregulated upon neural differentiation, but it is upregulated during neuronal maturation. This was confirmed by comparing Nup153 levels in maturing PCNs from DIV4 until DIV10 (Fig. 1C,1D). Furthermore, immunohistochemical analyses in the primary somatosensory cortex (S1) of adult mouse brains revealed that levels of Nup153 were significantly higher in NeuN-positive neurons than in NeuN-negative cells (p < 0.0001, Mann-Whitney test) (Fig. 1E, 1F). These data suggest that Nup153 levels are dynamically upregulated during neuronal maturation. Since Nup153 plays critical roles in both the activation and repression of developmental genes (*21, 25*), our observation prompted us to investigate the role of Nup153 in post-mitotic mature neurons, with a particular focus on post-mitotic neuron-specific genetic programs.

**Fig. 1.**
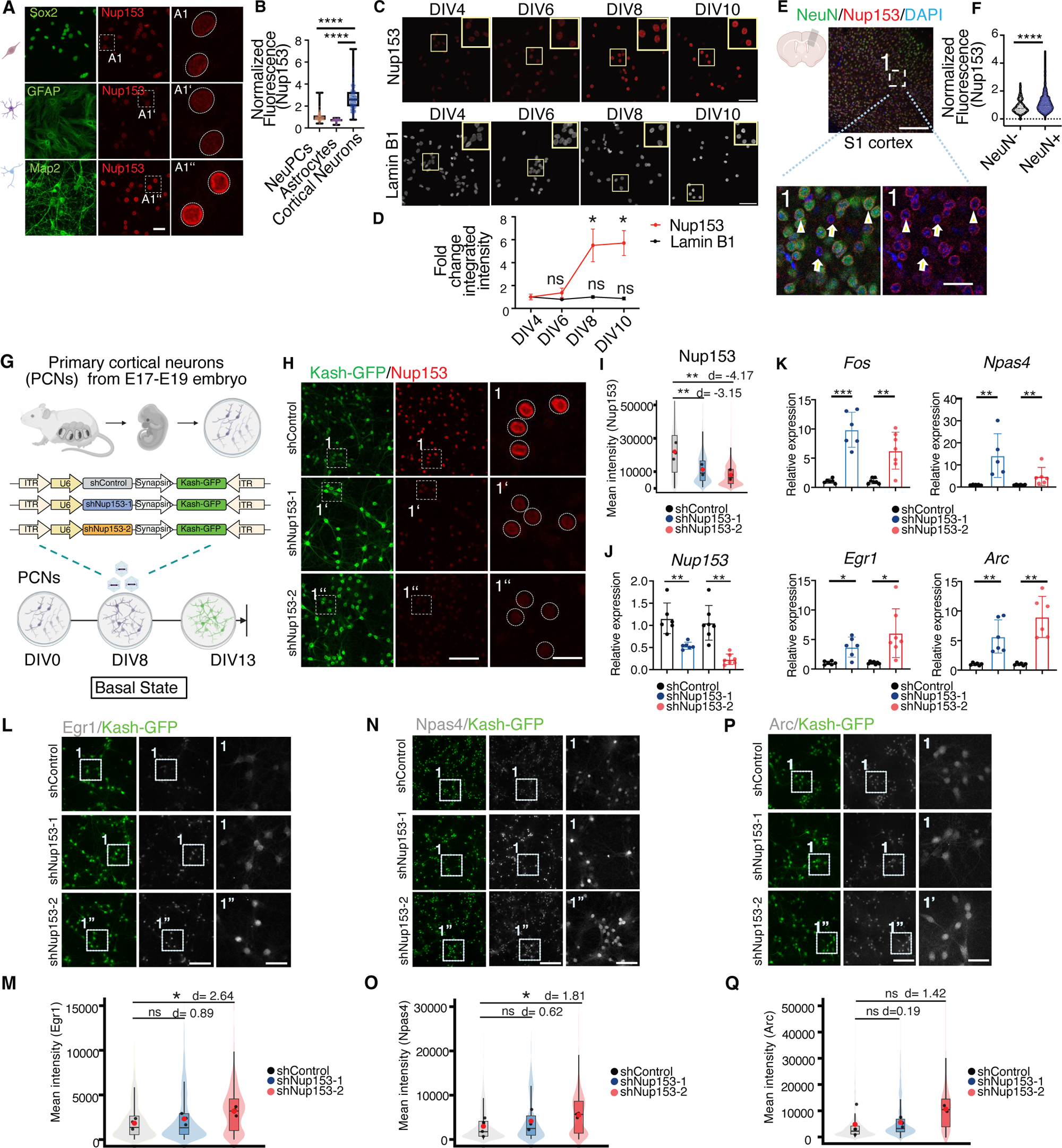
The essential role of Nup153 in balancing the levels of ARGs in basal neurons. (A) Different neural cell types showed distinct levels of Nup153 (red). Scale bars = 50 μm in the middle, 10 μm in the inset. (B) Quantification of Nup153 levels in (A), which depicts the normalized fluorescence intensity of Nup153 in MAP2^+^ neurons and GFAP^+^ astrocytes relative to Sox2^+^ NeuPCs. (****p < 0.0001 for NeuPCs compared to neurons and for astrocytes compared to neurons, Mann-Whitney test; n = 145 NeuPCs, n = 38 astrocytes, n = 220 neurons. Three independent experiments were conducted. The data are presented as a min to max whisker plot showing all points). (C) Representative immunostaining of Nup153 (top) and Lamin B1 (bottom) in PNCs across days in vitro (DIV4, 6, 8, and 10) (scale bar = 50 µm) (D) Quantification of integrated intensity fold changes (relative to DIV4 values) for Nup153 and Lamin B1 levels across days *in vitro*. Statistical significance was independently assessed for each marker and determined by comparing values against DIV4 via ANOVA followed by Dunńs or Dunnett’s correction accordingly. All p values are indicated as *p < 0.05 and “ns” when p > 0.05. (E) Confocal images of immunostaining against NeuN (green) and Nup153 (red) in the primary somatosensory cortex (S1) of the adult brain. The inset in the first image is magnified in the images to the right. Arrowheads indicate NeuN^+^ neurons, arrows indicate NeuN^-^ cells. Scale bar in the left panel = 200 μm. Scale bar in the images taken from the inset = 25 μm. (F) Quantification of Nup153 levels in NeuN^+^ mature neurons and NeuN^-^ cells (****p < 0.0001, Mann-Whitney test; n = 125 for NeuN^-^ cells and n = 1188 for NeuN^+^ neurons taken from the S1of three mice). (G) A schema for the Nup153 knockdown strategy in mouse PCNs at the basal state. (H) Confocal images of PCNs for KASH-GFP^+^ (green) and Nup153 (red). Scale bar = 100 μm (left) and Scale bar = 20 μm (right) (I) Quantification of Nup153 (red) levels in KASH-GFP^+^ (green) neurons. The integrated fluorescence intensity of Nup153 in KASH-GFP^+^ PCNs was quantified (**p < 0.001; shControl, 22038 ± 4098; shNup153-1, 11390 ± 2531; shNup153-2, 7993 ± 2435; Welch’s two sample t-test; n = 3 biological replicate; Cohen’s d = −4.17, shControl vs shNup13-2; Cohen’s d = −3.15, shControl vs shNup153-1). (J) The expression level of Nup153 mRNA in PCNs after Nup153 knockdown (**p = 0.0063, for 1.16 ± 0.35 in shControl, 0.54 ± 0.082 in shNup153-1, **p = 0.0011, 0.23 ± 0.12 in shNup153-2, Welch’s t-test; 5-7 independent experiments were conducted). (K) *Fos, Npas4, Egr1* and *Arc* mRNA expression levels after Nup153 knockdown (***p < 0.001, **p < 0.01, *p < 0.05. A Welch’s t-test was used for *Fos, Egr1*, and *Arc* and a Mann-Whitney test was used for *Npas4*. 5 to 7 independent experiments were conducted). (L) Upregulation of Egr1 protein levels after Nup153 knockdown in PCNs. Representative confocal images of KASH-GFP^+^ (green) and Egr1 (gray) in PCNs. Scale bar = 125 μm (middle) and Scale bar = 30 μm (right). (M) Quantification of Egr1 (grey) levels in KASH-GFP^+^ (green) neurons in (L). The integrated fluorescence intensity of Egr1 in KASH-GFP^+^ PCNs was quantified (*p < 0.05; ns = not significant; shControl, 1832 ± 606; shNup153-1, 2327 ± 520; shNup153-2, 3185 ± 440; Welch’s two sample t-test; n = 3 biological replicate; Cohen’s d=2.64 shControl vs shNup13-2; cohen’s d=0.89, shControl vs shNup153-1). (N) Representative confocal images of KASH-GFP^+^ (green) and Npas4 (gray) in PCNs. Scale bar = 125 μm (middle) and Scale bar = 30 μm (right) (O) Quantification of Npas4 (grey) levels in KASH-GFP^+^ (green) neurons in (M). The integrated fluorescence intensity of Egr1 in KASH-GFP^+^ PCNs was quantified (*p < 0.05; ns = not significant; shControl, 1832 ± 606; shNup153-1, 2327 ± 520; shNup153-2, 3185 ± 440; Welch’s two sample t-test; n = 3-4 biological replicates; Cohen’s d = 1.81 shControl vs shNup13-2; Cohen’s d = 0.62, shControl vs shNup153-1). (P) Representative confocal images of KASH-GFP^+^ (green) and Arc (gray) in PCNs. Scale bar = 125 μm (middle) and Scale bar = 30 μm (right). (Q) Quantification of Arc (grey) levels in KASH-GFP^+^ (green) neurons in (P). The integrated fluorescence intensity of Arc in KASH-GFP^+^ PCNs was quantified (ns = not significant; shControl, 4829 ± 5172; shNup153-1, 5558 ± 1497; shNup153-2, 10622 ± 1222; Welch’s two sample t-test; n = 4 biological replicates; Cohen’s d = 1.42, shControl vs shNup13-2, Cohen’s d = 0.19 shControl vs shNup153-1). The data for RNA expression level (Fig. 1K & 1J) are presented as mean ± standard deviation (s.d). The data for fluorescence intensity (Fig. 1M, 1O & 1Q) are presented as boxplot, with lines representing mean, red dot representing median and violin plot for individual cells. See also Fig. S1-S3 for extended data for Fig. 1.

### The essential role of Nup153 in balancing ARG expression levels in basal state neurons

To test whether Nup153 plays a critical role in the regulation of ARGs in basal state neurons (basal neurons), we first performed loss-of-function experiments. To mildly reduce levels of Nup153, we used previously validated shRNAs delivered by adeno-associated virus (AAV) vectors (*21*). To knock down Nup153 after neuronal maturation, AAVs harboring shRNAs against *Nup153* as well as Kash-EGFP were applied at DIV8 (Fig. 1G) (*26*). Five days after the application of AAVs, knockdown of Nup153 at the transcript and protein levels was confirmed by quantitative reverse transcription polymerase chain reaction (qRT-PCR) and immunocytochemistry for each of the two shRNAs (Fig. 1H-1J). shNup153-1 knocked down approximately 49 % of Nup153 expression, whereas shNup153-2 knocked down approximately 64 % of Nup153 expression, indicating a stronger knockdown using shNup153-2 (Fig. 1I). We did not observe any apparent changes in the density of GFP-positive cells or in the percentage of NeuN^+^ cells among GFP-positive cells (GFP; p > 0.05, Welch’s t-test; NeuN fraction, p > 0.1 Mann-Whitney test) (Fig. S1A-S1C), suggesting that knockdown of Nup153 did not exhibit apparent adverse effects in mature neurons. Similarly, knockdown of Nup153 did not increase the levels of active caspase-3 (Fig. S1D, S1E) (p = 0.59 for shControl vs. shNup153-1, p = 0.002 for shControl vs. shNup153-2, Mann-Whitney test). Moreover, the integrity of the nuclear lamina revealed by immunostaining for lamin B2 and MAB414, a marker for FG-repeat-containing nucleoporins, showed no apparent differences (Fig. S2A, S2B).

Next, we examined the transcript levels of several ARGs (*Fos, Egr1, Naps4,* and *Arc*) in the basal state without any stimulation. Strikingly, the levels of all tested ARG transcripts were significantly and consistently upregulated in Nup153-depleted neurons (Fig. 1K). Upregulation of Egr1, Npas4, Arc and Fos at the protein level in response to Nup153 depletion by each of the two shRNAs was also observed using immunocytochemistry, albeit to a lesser extent with shNup153-1, presumably owing to the lower level of Nup153 knockdown (Fig. 1L-1Q, S3A-S3B). Overall, these results suggest that Nup153 represses the expression of ARGs in mature neurons in the basal state. The upregulation of several ARGs indicated a potential increase in global transcription, and to test this possibility, we assessed nascent transcription levels by incubating cells with ethynyl-uridine (EU) for 6 hours. EU-labeled transcripts were visualized by click chemistry, and there were no obvious changes in the levels of EU-labeled transcripts in Nup153-depleted neurons (p > 0.05, Mann-Whitney test) (Fig. S3C, S3D). Our data supports the idea that, in mature neurons, Nup153 knockdown does not alter transcription levels in general.

### Global de-repression of neuronal genes including key ARGs in Nup153-depleted basal neurons

Our data indicated that Nup153 represses the expression of ARGs in mature neurons and thus may regulate neuronal responsiveness by balancing the expression of ARGs in the basal state. To investigate the extent to which Nup153 regulates genetic programs on a genome-wide scale in mature neurons, we performed RNA-seq analyses after Nup153 knockdown (Fig. 2A-2C, Fig. S4A-G, Table S1). The two shRNAs had similar effects (Fig. 2C, r = 0.46), with 82.8% of genes significantly dysregulated by one or both of the two shRNAs, showing changes in gene expression in the same direction (Fig. S4D, 1751/1973 in Q1:+/+ or Q3:-/-). Although the majority of DEGs showed a similar direction of dysregulation in response to each of the two shRNAs, there were some degrees of inconsistency between the two shRNAs (Fig.2C, Fig.S4D, Q2:+/- or Q4:-/+). Therefore, we used the genes commonly dysregulated by the two shRNAs, and we used shNup153-2 for the depletion of Nup153 in subsequent experiments due to its stronger effects. We defined genes that were commonly affected by the two shRNAs with high confidence (padj < 0.01 for both shRNAs, black) (Fig. 2B, 2C, Table S1) as “differentially expressed genes (DEGs)” and identified a total of 1070 DEGs (Q1-Q4), including 599 commonly upregulated DEGs (Nup153-repressed genes, genes in Q1:+/+) and 436 commonly downregulated DEGs (Nup153-activated genes, genes in Q3:-/-) (Fig. 2C, Fig. S4D, Table S1, see methods). Other than Nup153, other nucleoporins were not classified as DEGs (Fig. S4E).

**Fig. 2.**
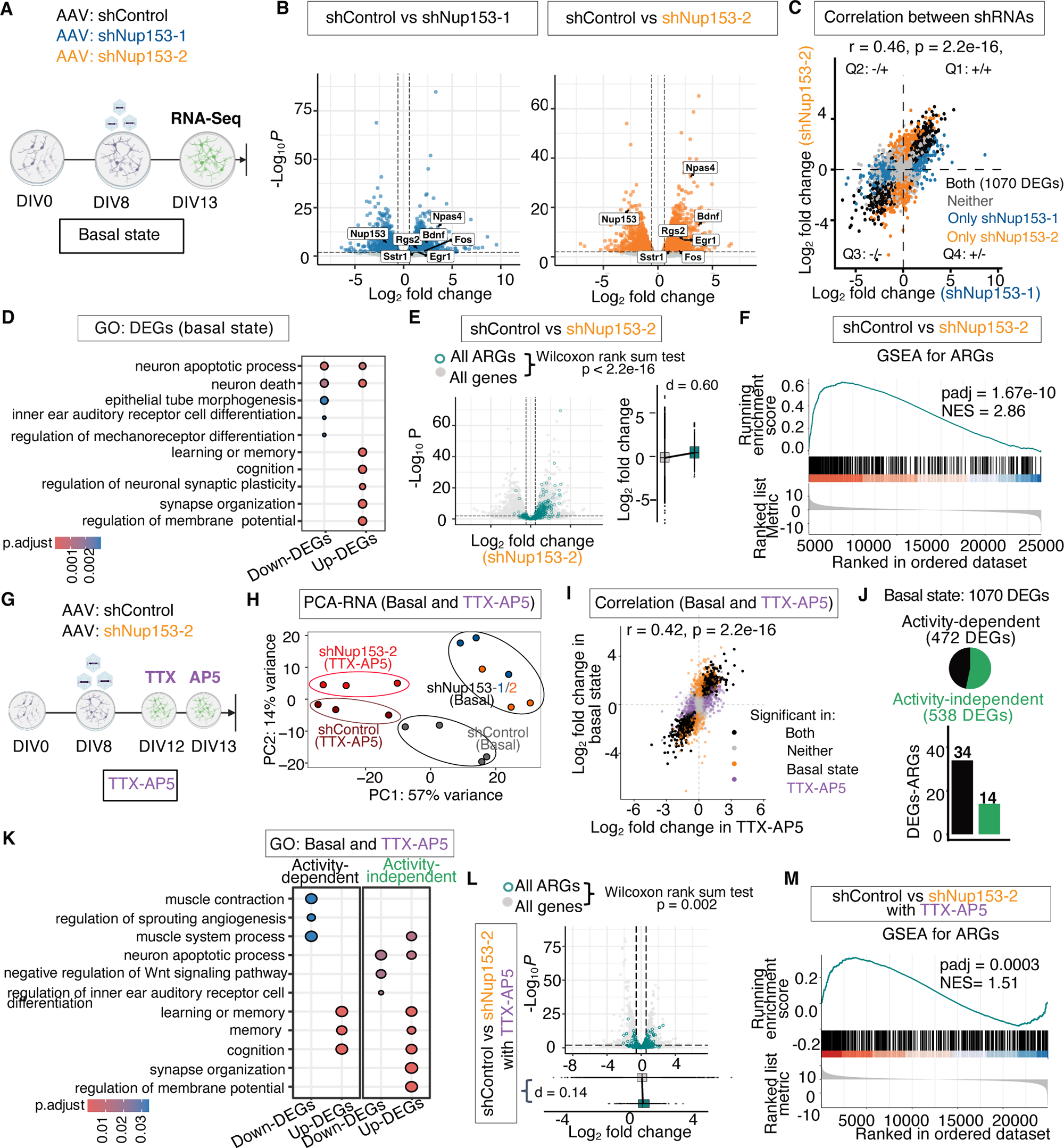
Global de-repression of ARGs in Nup153-depleted basal neurons. (A) A schema for the Nup153 knockdown strategy in basal neurons. (B) Volcano plots for DEGs after Nup153 knockdown by the two shRNAs. DEGs (blue represents knockdown by shNup153-1 and orange represents knockdown by shNup153-2) are defined as log_2_ fold change greater than 0.58 with a 1% FDR cut-off. (C) Correlation of DEGs between the two shRNAs. DEGs induced by the two shRNAs were similar (r = 0.46, p = 2.2e-16, Pearson’s correlation). DEGs common between the two shRNAs are shown as black dots. Different quadrants are shown to indicate the consistency of the effects, in terms of the direction of regulation, between the two shRNAs. (D) GO analysis for Nup153-commonly dysregulated genes, subdivided as either upregulated or downregulated upon Nup153 depletion. (E) A volcano plot highlighting the differential expression of previously discovered ARGs by Kim et.al(*8*). The boxplot depicts the log_2_ fold change of ARGs in comparison to all other genes upon Nup153 depletion (p < 2.2e-16, Wilcoxon Rank sum test; Cohen’s d = 0.6, 95% CI [0.51, 0.70]). (F) Gene set enrichment analysis (GSEA) for ARGs(*8*) within the transcriptome of Nup153-depleted neurons. (G) A schema for the Nup153 knockdown strategy in TTX-AP5 treated neurons. (H) A PCA plot for the transcriptomes of Nup153-depleted neurons in the basal state and after treatment with TTX-AP5. (I) Correlation between DEGs in in basal neurons and TTX-AP5 treated neurons (r = 0.42, p = 2.2e-16, Pearson’s correlation). Common DEGs between the two conditions are highlighted as black dots. (J) (Top) The fraction of basal activity-dependent DEGs and basal activity-independent DEGs. (Bottom) The bar graph shows the number of activity-dependent DEG-ARGs and activity-independent DEG-ARGs. (K) GO analysis for Nup153-dysregulated genes subcategorized as either “basal activity-dependent” or “basal activity-independent”. Within these subcategories, the genes were further subcategorized as either “upregulated” or “downregulated” upon Nup153 depletion, and then enriched for gene ontologies in each category. (L) A volcano plot highlighting differential expression of the previously discovered ARGs in Nup153-depleted neurons treated with TTX-AP5. The boxplot depicts quantification of log_2_ fold changes in ARGs in comparison to those of all other genes upon Nup153 depletion (p = 0.002, Wilcoxon Rank sum test, Cohen’s d = 0.14, 95% CI [0.05, 0.24]). (M) GSEA for the previously discovered ARGs in the presence of TTX-AP5 in Nup153-depleted neurons. See also Fig. S4-S5 as extended data for Fig. 2A-2F and Fig. S6 as extended data for Fig. 2G-2M. See also Table S1 as supplementary tables for Fig. 2

Gene Ontology (GO) analysis revealed that Nup153-repressed genes were enriched in learning/memory (padj = 2.09e-09, Benjamini-Hochberg), cognition (padj = 1.75e-08, Benjamini-Hochberg), and the regulation of membrane potential (padj = 7.36e-08, Benjamini-Hochberg), indicating that Nup153 is required for the regulation of neuronal functions (Fig. 2D, Table S1). To investigate whether the ARGs were globally upregulated when Nup153 was knocked down, changes in the previously discovered 568 ARGs were assessed (*8*). We found significant upregulation of ARGs upon Nup153 depletion compared to the rest of the expressed genes in PCNs (Fig. 2E, S4F) (log_2_ fold change of all 568 ARGs vs. log_2_ fold change in all other genes for shControl vs. shNup153-1 condition: p = 0.004, Cohen’s d = 0.14, 96/568 significantly upregulated ARGs; log_2_ fold change in ARGs vs log_2_ fold change in all other genes for shControl vs. shNup153-2 condition: p = 2.2e-16, Cohen’s d = 0.60, 141/568 significantly upregulated ARGs). Additionally, the global enrichment of ARGs was assessed using gene set enrichment analysis (GSEA)(*8, 27*). ARGs were significantly enriched upon depletion with both the shRNAs (Fig. 2F, Fig. S4G) (shNup153-1: padj = 4.29e-4, NES = 1.43; shNup153-2: padj = 1.67e-10, NES = 2.86). In summary, a total of 187 ARGs were dysregulated by at least one shRNA and 81.3% (152/187) of these dysregulated ARGs exhibited changes in their expression in the same direction in response to both of the two shRNAs (Fig. S5A, S5B, r = 0.62), indicating consistency in the ARGs dysregulation caused by both shRNAs (Fig. S5A-5D). To ensure consistency in our analysis, we evaluated ARGs commonly dysregulated by the two shRNAs (DEG-ARGs) and identified 47 upregulated DEG-ARGs and 2 downregulated DEG-ARGs. We also investigated whether specific ARG subtypes were differentially affected by Nup153 depletion. ARGs were categorized as rapid primary responsive genes (rPRGs), delayed PRGs (dPRGs) or secondary responsive genes (SRGs) based on the previous work (*14*). All ARG subtypes were affected, rPRGs being impacted the most (Fig. S5E, S5F). These results clearly indicate that Nup153 represses ARG expression in mature post-mitotic neurons, and the loss of Nup153 de-represses ARG expression without requiring additional stimulation.

Since Nup153 is deeply involved in epigenetic regulation in stem cells (*21, 22*), it is plausible that Nup153 regulates ARGs through epigenetic mechanisms in post-mitotic neurons. However, it is also possible that Nup153-depletion initially leads to an increase in levels of basal neuronal activity or an increase in synaptic activity through intermediate effectors, resulting in the induction of ARGs (Fig. S6A). To experimentally test these possibility, basal neuronal activity were blocked by tetrodotoxin (TTX) and DL-2-amino-5-phosphonopentanoic acid (AP5) (Fig. 2G, S6A, S6B), and genome-wide gene expression changes in Nup153-depleted neurons were examined by RNA-seq. PCA of the transcriptomes revealed that TTX-AP5 treatment significantly shifted the treated samples away from the vehicle samples (Fig. 2H). However, Nup153-depleted samples were still clearly differentiated from control samples in the presence of TTX-AP5, indicating a contribution of Nup153-dependent but neuronal basal activity-independent gene dysregulation (Fig. 2H). Indeed, in the presence of TTX-AP5, we identified 1402 differentially expressed genes (Fig. 2I, S6C, Table S1). 719 genes were upregulated and 683 genes were downregulated by Nup153 knockdown in the presence of TTX-AP5 (Fig. S6C, Table S1). There was a positive correlation between genes dysregulated upon Nup153 depletion in the basal state and in the presence of TTX-AP5 (Fig. 2I, r = 0.42, p = 2.2e-16, Pearson correlation). To identify the effects of Nup153 depletion in combination with basal neuronal activity, we classified the intersection of the set of 1070 DEGs identified in the basal state with the set of genes dysregulated in the presence of TTX-AP5 and shNup153-2 as basal activity-independent genes (538 genes), and the non-intersecting set of DEGs in the basal state was classified as basal activity-dependent genes (472 genes) (Fig. 2J, S6D, see methods). GO analysis revealed enrichment of basal activity-independent Nup153-upregulated genes in synapse organization (padj = 1.74e-05, Benjamini-Hochberg), memory (padj = 0.003, Benjamini-Hochberg), and cognition (padj = 0.0003, Benjamini-Hochberg) (Fig. 2K, Table S1). On the other hand, apoptotic processes and the Wnt signaling pathway were enriched in downregulated genes, suggesting that these are also involved in the processes of basal activity-independent mechanisms. Furthermore, the expression levels of 568 ARGs showed higher fold changes than all other genes (p = 0.002 Wilcoxon rank sum test, Cohen’s d = 0.14, 54/568 significantly upregulated ARGs) (Fig. 2L), and GSEA analysis revealed significant enrichment of ARGs within Nup153-repressed genes in the presence of TTX-AP5 (Fig. 2M, padj = 0.0003, NES =1.51). These data suggest some several neuronal genes including ARGs are regulated by Nup153 in combination with basal neuronal activity, whereas others are regulated by Nup153 independent of neuronal activity.

To specifically determine which ARGs belong to two categories, we identified ARGs among the DEGs that were either basal activity-dependent or basal activity-independent. Our analysis revealed 14 DEG-ARGs as basal activity-independent and 34 DEG-ARGs as basal activity-dependent (Fig. 2J, S6E, S6F). As expected, the upregulation of activity-dependent DEG-ARGs induced by Nup153 depletion is attenuated in the presence of TTX-AP5 (Fig. S6E). However, there was still a moderate but consistent, increase in their expression even in the presence of TTX-AP5 as observed in the z-score/normalized count boxplot (Fig. S6F, Cohen’s d = 0.12 for shControl vs shNup153-2 ARGs). Since *Egr1* is not part of the previous defined list of 568 ARGs (*8*), we also assessed it separately. Notably, we found *Egr1* to be significantly upregulated upon Nup153 depletion in both a basal activity-dependent and a basal activity-independent manner (Fig. S6F). Similar trends were also observed using qPCR (Fig. S6B,6F) (qPCR mean levels, *Fos*, shControl 0.52 ± 0.31 vs shNup153-2 0.90 ± 0.67; *Egr1*, shControl 0.52 ± 0.31 vs shNup153-2 0.90 ± 0.67), indicating that *Fos/Egr1* RNA expression levels are mildly increased after Nup153 depletion in the presence of TTX-AP5. These data support the idea that a combination of Nup153 depletion and basal neuronal activity is essential for de-repressing these ARGs. Altogether, our results suggest that some Nup153-regulated ARGs are more influenced by basal activity but are still mildly influenced by Nup153 depletion, while other Nup153-regulated ARGs are less sensitive to the level of basal neuronal activity though it sill exerts a partial influence.

### Dose-dependent repression of ARGs by Nup153

Our data showed that mature neurons express higher levels of Nup153 than other neural cell types and that Nup153 is essential for the regulation of neuronal genes including ARGs in basal neurons. These observations raised the hypothesis that higher Nup153 levels in mature neurons are essential for keeping those genes at low levels in the basal state, thus maintaining neuronal responsiveness through a high signal-to-noise ratio for ARG expression upon neuronal stimulation. To test whether higher levels of Nup153 in neurons can further suppress ARGs, endogenous Nup153 was increased using a CRISPR activation system (*28*). Two lentiviral vectors, one harboring dCas9-VP64 and another harboring either a scrambled sequence or an sgRNA targeting the Nup153 promoter, were simultaneously introduced. Five days later, a high co-infection efficiency of the two lentiviruses was confirmed (Fig. 3A, GFP/RFP double-positive cells, 67.82 ± 25.49% cells infected with the virus carrying scrambled sgRNA; 68.33 ± 13.60% cells infected with the virus carrying Nup153-sgRNA). Upregulation of Nup153 was confirmed by qRT-PCR and immunocytochemistry (Fig. 3C-3E) (immunocytochemistry, p < 0.001, Welch’s *t*-test; qRT-PCR, p < 0.05). Importantly, the upregulation of Nup153 in mature neurons reduced the levels of *Fos* and *Egr1* expression in basal neurons (Fig. 3E). Combining the results of loss-of-function and gain-of-function experiments indicates that the expression levels of ARGs in mature neurons are affected by the levels of Nup153 in a dose-dependent manner.

**Fig. 3.**
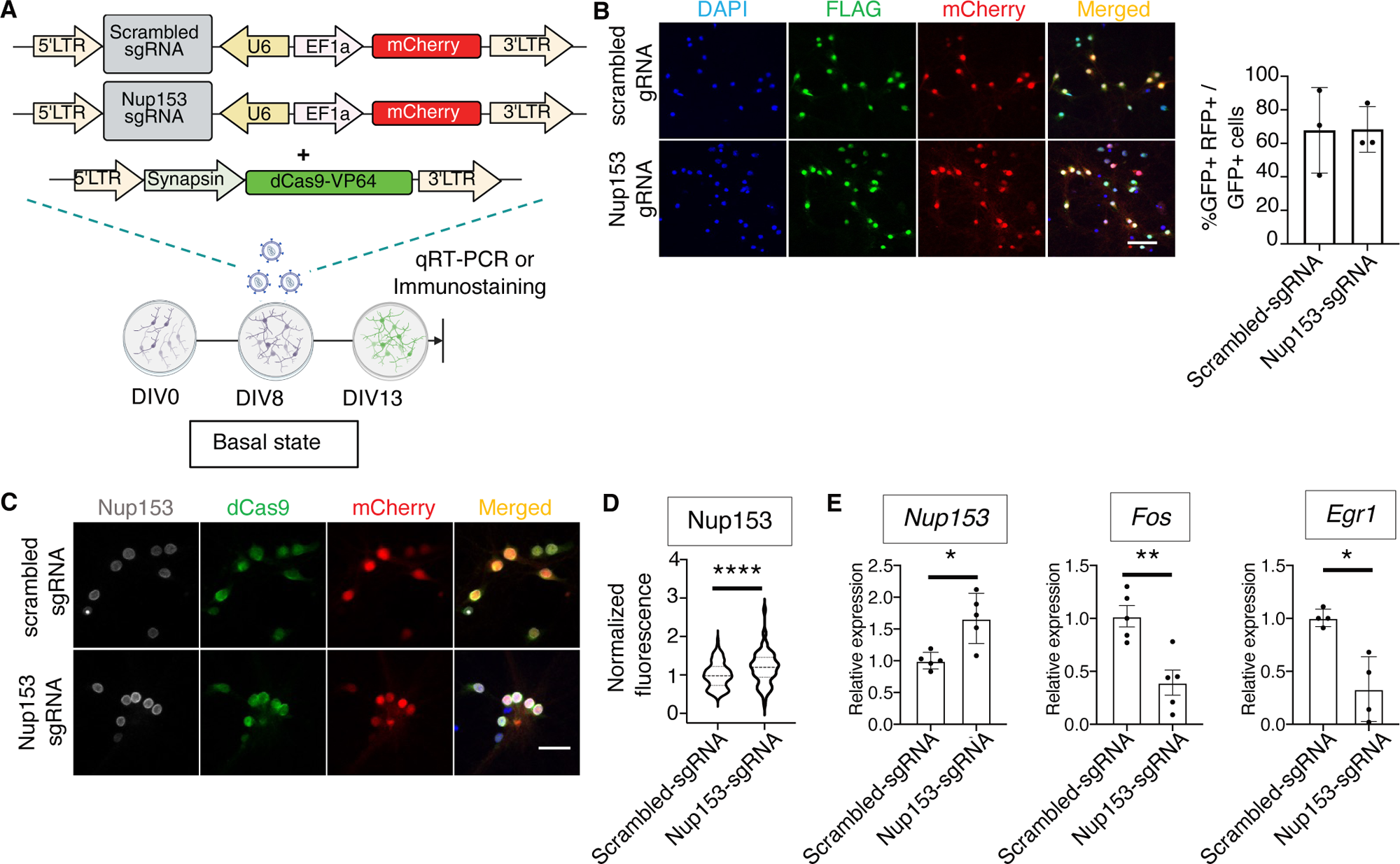
Dose-dependent repression of ARGs by Nup153. (A) A schematic illustration of the CRISPR-based overexpression strategy. PCNs were treated at DIV8 with a combination of two lentiviruses (LV) to mediate the neuron-specific expression of dCas9 fused with VP64 (Syn-dCas9-VP64) along with a sgRNA targeting the promoter of Nup153 (sgRNA-Nup153; mCherry) or a scrambled sequence (scrambled sgRNA; mCherry). (B) Representative confocal images of PCNs at DIV13 transduced with a combination of Syn-dCas9-VP64 and sgRNA-Nup153 or scrambled sgRNA. PCNs were stained for dCas9 (green) and DAPI (blue). The percentage of DAPI^+^ cells co-transduced with dCas9-VP64 (green) and sgRNA-Nup153/scrambled sgRNA (red), from three independent experiments. Scale bar = 50 μm. (C) Representative confocal images of PCNs transduced with two LVs and stained for nucleoporin Nup153 (grey) and dCas9 (green). Scale bar = 25 μm. (D) Quantification of Nup153 levels in (C). The graph depicts the normalized fluorescence intensity of Nup153 (grey) in DAPI^+^/GFP^+^/mCherry^+^ PCNs (*p < 0.05, **p < 0.01, Welch’s t-test, n = 131 cells for scrambled sgRNA, n = 124 cells for Nup153-sgRNA. The data are presented as violin plot). (E) Nup153, Fos, and Egr1 mRNA expression levels after the overexpression of Nup153 in basal neurons (*p < 0.05, **p < 0.01, Welch’s t-test, 4-5 independent experiments were conducted. The data are presented as mean ± standard deviation (s.d)).

### Nup153-directed, basal activity-dependent and -independent regulation of chromatin organization

Accumulating evidence suggests that open chromatin states dictate gene expression, and that changes in chromatin accessibility of functional genomic elements directly and/or indirectly instruct ARG’s responsiveness in functional neurons (*13, 20*). Because Nup153 has been shown to regulate chromatin accessibility in NeuPCs (*21*), we next investigated whether Nup153 regulates chromatin accessibility in mature neurons and how it relates to gene regulation. To assess changes in chromatin accessibility, an assay for transposase-accessible chromatin with sequencing (ATAC-Seq) was used (Fig. 4A) (*29*). In the basal state, 29658 and 42605 open chromatin regions were identified using the Genrich pipeline in control and Nup153-depleted neurons, respectively. In the presence of TTX-AP5, 36863 and 34913 open chromatin regions were identified in control and Nup153-depleted neurons, respectively (Table S2). Curiously, a PCA plot of the ATAC-seq data clearly separated Nup153-depleted samples with and without TTX-AP5, but all control samples clustered together even with TTX-AP5 treatment (Fig. 4B). This result indicates that basal neuronal activity has little effect on global chromatin accessibility and only affects chromatin accessibility when Nup153 is depleted. Furthermore, our finding that TTX-AP5-treated Nup153-depleted samples were clearly separated from Nup153-depleted basal neurons indicates that significant changes in chromatin accessibility occur after Nup153 knockdown, independent of neuronal activity. To gain an overview of genome-wide changes in open chromatin status, we identified 6255 differential accessible peaks (DAPs) in the basal state and 6156 DAPs in the TTX-AP5-treated state (Fig. S7A). We found that the major fraction of DAPs were gained-open peaks in both states (4209 DAPs, 67.6% in the basal state; 5372 DAPs, 87.3% in the TTX-AP5-treated state)(Fig. S7A, Table S2), indicating a critical role of Nup153 in restricting accessibility, independent of neuronal activity. Next, we analyzed the overlap between DAPs in the basal and TTX-AP5-treated states. We observed that 29.8% (1866/6255) of DAPs in the basal state overlapped with those in the TTX-AP5-treated state (overlapping DAPs), whereas 70.2% (4387/6255) of DAPs were non-overlapping (Fig. 4C, Table S2). Based on our observations at the RNA level, we assessed whether overlapping DAPs were independent of basal activity and if Nup153 could regulate chromatin accessibility at two different levels, in combination with basal neuronal activity. To this end, we created a heat map showing the direction of change in chromatin accessibility upon Nup153 depletion for both overlapping (activity-independent) and non-overlapping (activity-dependent) DAPs (Fig. 4D, 4E). Chromatin accessible regions were classified by hierarchical clustering based on changes in accessibility in the basal state. Notably, the overlapping DAPs exhibited consistent increases or decreases in accessibility upon Nup153 depletion in both the basal and TTX-AP5-treated states. These data suggest that the accessibility of these chromatin regions is regulated by Nup153, independent of basal neuronal activity. Moreover, changes in non-overlapping DAPs also showed a similar trend in the TTX-AP5-treated condition compared to the basal state, albeit to a lesser extent (Fig. 4E). These data indicate the following important points: (i) Nup153 depletion leads to significant changes in chromatin accessibility in neurons. (ii) Similar to transcriptomic changes, Nup153 regulates chromatin accessibility at two levels: one fraction is independent of basal activity, while the other partially depends on it. In a Nup153-dependent manner, both basal activity-dependent and basal activity-independent changes in chromatin accessibility tend to be affected in the same direction, regardless of neuronal activity.To establish a direct relationship between transcriptome-level changes resulting from Nup153 depletion and changes in chromatin accessibility, we first annotated all DAPs to nearby genes using ChIPSeeker (*30*). We found that 315 of 1070 DEGs were located near DAPs in the basal state (Fig. 4F, Table S2). Of these 315 DAP-associated DEGs, 173 were basal activity-independent and 132 were basal activity-dependent DEGs. In the basal state, we identified 266 DAPs (205 accessibility-gained DAPs, 61 accessibility-lost DAPs) and 206 DAPs (152 accessibility-gained DAPs, 53 accessibility-lost DAPs) associated with basal activity-independent and activity-dependent DEGs, respectively. Gene Ontology analysis of the basal activity-dependent or -independent DEGs associated with accessibility-gained DAPs indicated an enrichment of neuronal function terms, such as dendrite development and regulation of membrane potential (Fig. 4G, Table S2). These results are consistent with the involvement of Nup153-directed nuclear architecture in neuronal responsiveness. Subsequently, we visualized the aggregated changes in chromatin accessibility associated with basal activity-dependent and basal activity-independent DEGs (Fig. 4H,4I). Our analysis revealed a pattern similar to that observed for changes in global accessibility (Figure 4D, 4E). Specifically, we observed consistent directional changes in chromatin accessibility associated with both basal activity-dependent and activity-independent DEGs. To further understand the association between open chromatin status and changes in gene expression, we correlated changes in chromatin accessibility with changes in gene expression. Consistent with previous reports (*20, 31*), changes in chromatin accessibility were mildly correlated with changes in gene expression in both conditions (r = 0.29, p = 2.2e-16, Pearson’s correlation for the basal state; r = 0.15, p = 3.02e-07, Pearson’s correlation for the TTX-AP5-treated state) (Fig. S7B). Overall, these data suggest that Nup153 directs chromatin accessibility in the basal state. Nup153 depletion leads to changes in chromatin accessibility, and the extent of the changes can be influenced in both a basal activity-dependent and -independent manner.

**Fig. 4.**
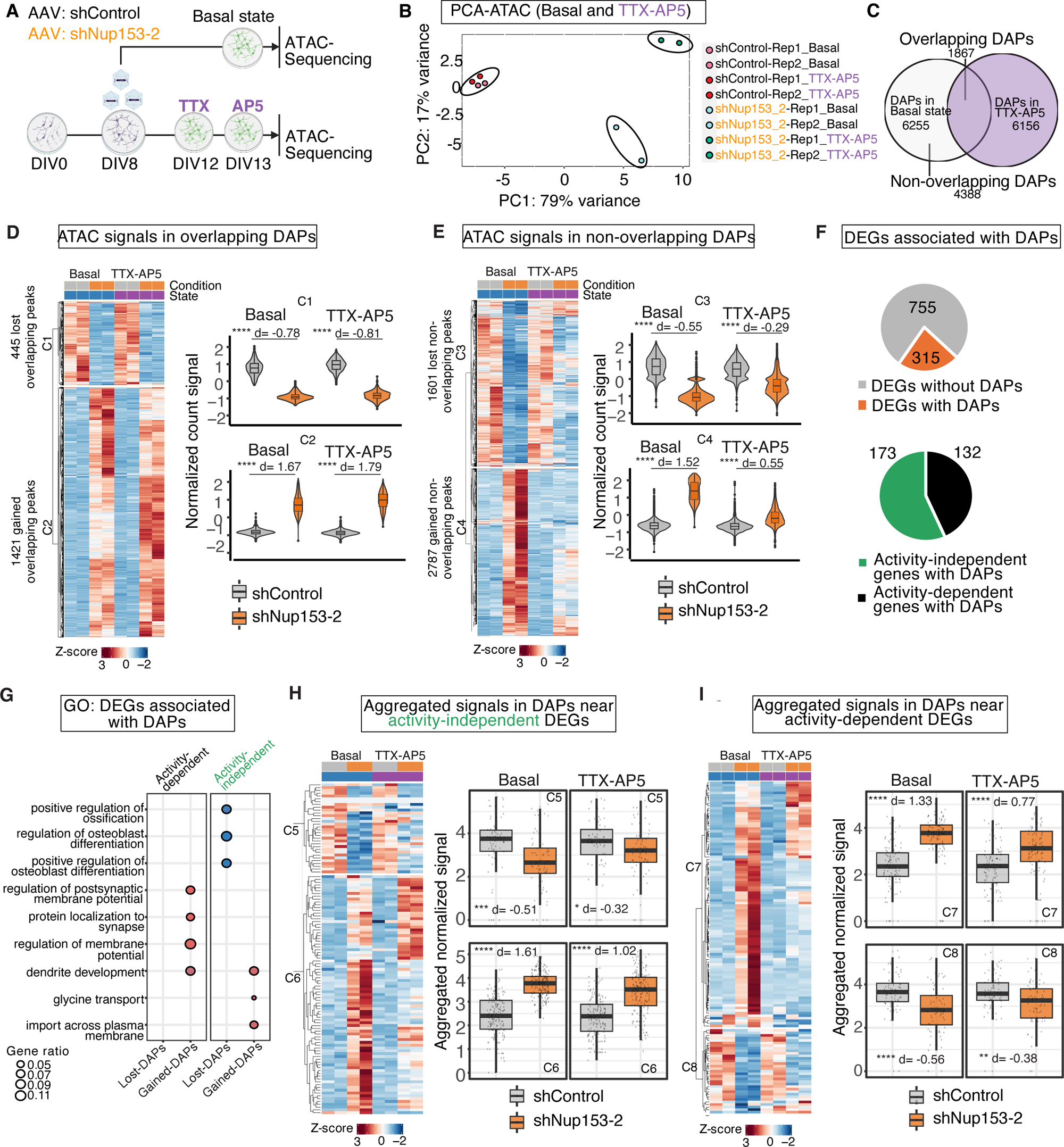
Nup153-directed, basal activity-dependent and -independent regulation of chromatin organization. (A) A schematic illustration of the experimental plan for ATAC-Seq in basal and TTX-AP5 treated neurons. (B) A PCA of ATAC-Seq data from control and Nup153-depleted neurons in the basal and TTX-AP5-treated conditions. (C) A venn diagram illustrating overlap between DAPs in basal and TTX-AP5-treated neurons. (D) Left) A comparison of chromatin accessibility in overlapping DAPs. Count values are visualized in the basal or TTX-AP5-treated conditions upon Nup153 depletion. The heatmap is k-means clustered into the subclusters C1 and C2, which indicate the presence of DAPs with gained accessibility (C2) and lost accessibility (C1) upon Nup153 depletion. (Right) Normalized count values for the subclusters C1 and C2 are also visualized as boxplots integrated with violin plots (****p < 0.0001, Wilcox-test; C2: Cohen’s d = 1.67 (Basal) and 1.79 (TTX-AP5); C1: Cohen’s d = −0.78 (Basal), and −0.81 (TTX-AP5). (E) (Left) A comparison of chromatin accessibility in non-overlapping DAPs. The heatmap is k-means clustered into the subclusters showing groups with consistently gained or lost accessibility in both the basal and TTX-AP5-treated conditions upon Nup153 depletion. (Right) Normalized count values are visualized as combined boxplot/violin plots (****p < 0.0001, Wilcox-test; C4: Cohen’s d = 1.52 (Basal), and 0.55 (TTX-AP5); C3: Cohen’s d = −0.55 (Basal), and −0.29 (TTX-AP5)). (F) (Top) A pie chart showing the fraction of Nup153-regulated DEGs associated with DAPs. (Bottom) A pie chart showing the fraction of the DEGs with DAPs (orange from Top) sub-categorized as basal activity-dependent genes) or basal activity-independent genes (black). (G) GO analysis for activity-dependent or activity-independent DEGs associated with DAPs, which are further categorized into genes with gained or lost accessibility. (H) (Left) A comparison of chromatin accessibility at DAPs associated with activity-independent DEGs. Count values were visualized and compared in the basal or TTX-AP5-treated conditions upon Nup153 depletion. Theheatmap is k means-clustered into the subclusters C5 and C6. (Right) Normalized count values for subclusters C5 and C6 are visualized as boxplots. (****p < 0.00001,***p < 0.0001,*p < 0.05, Wilcox-test; C5: Cohen’s d = −0.51 (Basal), and −0.32 (TTX-AP5); C6: Cohen’s d = 1.61 (Basal state), and 1.02 (TTX-AP5)). (I) (Left) A comparison of chromatin accessibility within DAPs associated with activity-dependent DEGs. Count values were visualized in the basal or TTX-AP5-treated conditions upon Nup153 depletion. The heatmap is k-means-clustered into the subclusters C7 and C8. C7/C8 indicate the presence of DAPs with gained accessibility (C7) and lost accessibility (C8). (Right) Normalized count values for the subcluster C7 and C8 are also visualized as boxplots. (****p < 0.00001, ***p < 0.0001, *p < 0.05, Wilcox-test; C7: Cohen’s d = 1.33 (Basal state), and 0.77 (TTX-AP5); C8: Cohen’s d = −0.56 (Basal state), and −0.38 (TTX-AP5)). See also Fig. S7-S9 as extended data Fig. 4. For supplementary tables corresponding to Fig.4, see Table S3.

To understand the effect of Nup153 depletion on chromatin accessibility near ARGs, we compared changes in accessibility for all ARGs (*8*), as well as for the previously defined DEG-ARGs (Fig. 2J). In the basal condition, 143 ARGs were associated with Nup153-dependent DAPs (Fig. S8A). Of these, 10 DEG-ARGs were associated with DAPs and were also affected in a basal activity-independent manner at the RNA level. Meanwhile, 18 basal activity-dependent DEG-ARGs were associated with DAPs (Fig. S8A, lower graph). We assessed the aggregated changes in chromatin accessibility associated with basal activity-dependent and activity-independent DEG-ARGs within the DAPs associated with these ARGs (Fig. S8B, S8C). For DAPs associated with activity-independent DEG-ARGs, we observed overall gained accessibility in both the basal and TTX-AP5-treated states for the majority of ARGs in this category (Fig. S8B, S8D). Conversely for DAPs associated with activity-dependent DEG-ARGs, we observed a sharp increase in accessibility in the basal state for ARGs such as *Fos, Bdnf, Npas4, Egr1,* while most of the accessibility was largely attenuated in the TTX-AP5-treated state (Fig. S8C, S8F). However, the accessibility of some DAPs associated with ARGs, such as *Grin1* and *Stk40*, was still increased in the TTX-AP5-treated state (Fig. S8C, S8E). Interestingly, when we summed signals across all consensus peaks (see methods), we observed an overall increase in accessibility near some of basal activity-dependent DEG-ARGs such as *Egr1* and *Npas4* in both the basal and TTX-AP5 conditions (Fig. S8G). These data indicate similar findings for basal activity-dependent ARGs in the transcript levels (Fig. S6E,6F). That is to say, even when the de-repression of ARGs triggered by Nup153-depletion was attenuated in the presence of TTX-AP5, we still observed an increase in accessibility for some basal activity-dependent ARGs. This observation also supports the idea that Nup153 primarily regulates chromatin architecture, while the extent of its effect is influenced by the dependence of target genes on neuronal activity.

To probe the distinct mechanisms underlying the changes in chromatin accessibility between basal activity-dependent and basal activity-independent regulation, we analyzed differences in transcription factor occupancy between non-overlapping DAPs and overlapping DAPs using TOBIAS (Fig. 4C, Fig. S9)(*32*). Our analysis revealed a significant enrichment in AP-1/bZIP family transcription factor footprints within the non-overlapping DAPs (S9A). The top 25 motif clusters whose occupancy changed in the basal state upon Nup153 depletion in the non-overlapping DAPs included the C_FOSL1 cluster, along with several motifs associated with neuronal activation, including the Jun-1, JUNB, and FOS::JUND motifs (Table S2). Upon TTX-AP5 treatment, we observed a clear attenuation in transcription factor occupancy for these motif clusters in non-overlapping DAPs (Fig. S9A). For instance, the increased occupancy of C_FOSL1 was substantially reduced in the TTX-AP5-treated state (Fig. S9A, Table S2, binding score basal shControl vs shNup153-2 = −1.06, binding score TTX-AP5 shControl vs shNup153-2 = −0.36), although the enrichment of these motifs was not completely abolished. A similar pattern was observed across other motif clusters as well (Fig. S9A, Table S2). These findings suggest that, in the non-overlapping DAPs, Nup153 restrains AP-1 binding, and that its depletion could alter the occupancy of several key transcription factors such as Fos and Fosl1 in a manner that depends on basal neuronal activity.

By contrast, the top 25 motif clusters in the overlapping DAPs exhibited different motif enrichment patterns compared to the non-overlapping DAPs (Fig. S9B). This suggests that the two DAP classes are likely bound and regulated by distinct transcription factors. In the overlapping DAP, changes in motif enrichment patters were predominantly characterized by a loss of transcription factor occupancy upon Nup153 depletion. However, we also observed significant gains in several neuronal activity-relevant motif clusters, such as C_Npas4 and C_Fos::JUN (Table S2, C_Npas4 binding score TTX-AP5 shControl vs shNup153-2 = −0.44, C_Fos::JUN binding score TTX-AP5 shControl vs shNup153-2 = −0.37). Notably, the occupancy changes in these clusters were maintained between the basal and TTX-AP5-treated states (Table S2). Together, these results suggest that chromatin accessibility at the overlapping and non-overlapping DAPs are regulated by distinct combinations of transcription factors and Nup153.

In order to promote neuronal gene expression, neuronal promoters and enhancers undergo a coordinated change in chromatin states from a basal to an active state (*8*). The changes we observed in the occupancy of several key transcription factors, including AP-1 in Nup153-regulated chromatin areas suggests that Nup153 control specific neuronal chromatin states associated with these elements. To gain a better understanding of how Nup153 contributes to regulating specific chromatin states and consequently neuronal gene expression, we employed ChromHMM (*33*), which uses multivariate chromatin information to annotate chromatin states. Firstly, we used the published ChIP-Seq datasets from basal and active cortical neurons to train a ChromHMM model (*8, 33*). The ChromHMM model predicts the existence of hidden chromatin states in basal and active neurons (Fig. S9C, S9D). To train the model to define the chromatin states specific to basal and active neurons, the distinct binding of transcription factors such as Fos, Jun, CREB and RNA polymerase 2 (Pol2), as well as the presence of histone modifications were used (*8, 34*). For instance, state 2 was defined by the co-occurrence of Fos/Jun/CBP binding and the histone modifications H3K27ac/H3K4Me3, and defined as enhancers in the active state (Fig. S9C). Subsequently, the enrichment of chromatin states in DAPs from Nup153-depleted neurons was projected onto the defined chromatin states by ChromHMM, revealing a high degree of overlap between DAPs and active enhancers (state 2) in the state of active neurons (overlap score = 88.77, −log_10_ (p) > 300) in contrast to that of basal neurons (overlap score = 37.45, −log_10_ (p) = 48.2) (Fig. S9D, see methods). Furthermore, we found that DAPs overlapped more frequently with poised enhancers (states 4 and 6) in basal state neurons than in active neurons (Fig. S9D). These results support a model in which Nup153 is involved in the regulation of chromatin states, balancing the permissiveness of functional elements in the neuronal genome.

### Direct chromatin regulation by Nup153 in basal neurons

Our findings support the function of Nup153 in the regulation of chromatin organization and the regulation of neuronal states in neurons. To investigate its direct function on chromatin organization, we employed Targeted DamID followed by next-generation sequencing (TaDa-seq) (Fig. 5A, S10)(*35*)(*36*). TaDa relies on ectopic expression of Dam or Dam-Nup153 fusion constructs but incorporates an upstream primary open reading frame that substantially reduces translation efficiency and was previously shown to prevent overexpression phenotypes (*35*). Lentivirus particles carrying DAM control or DAM-Nup153 were applied at DIV8, and samples were collected on DIV13. Using four biological replicates, we identified 32290 common peaks (Fig. 5A-C, S10A-C, Table S3). As expected, Nup153 peaks were enriched in promoters or distal intergenic regions including for several ARGs such as *Fos, Egr1, BDNF*, and *Npas4* (Fig. 5B-D, S10C-D), and genes near Nup153 peaks were highly enriched for GO terms related to neuronal functions (Fig. 5D, Table S3). We then analyzed the genes directly regulated by Nup153. For those analyses, we defined direct regulation as occurring when Nup153 was found binding near to DEGs. We found that 59.7% of the Nup153-repressed genes were directly bound by Nup153 (odds ratio = 1.65, p = 2.08e-09, Fisher exact test), while 47.2% of the Nup153-activated genes were directly bound by Nup153 (odds ratio = 0.99, p = 0.88, fisher exact test) (Table S3), indicating a tendency for Nup153-binding to be associated with gene repression (p = 4.25e-05, chi-squared test) (Fig. 5E). 61.4% of all ARGs (349 genes, odds ratio = 1.77, p=2.62e-11, Fisher exact test) were bound by Nup153 (Fig. 5F), of which 12.3 % of Nup153-bound ARGs were DEGs upon Nup153 depletion (43 genes), whereas only 0.01% of Nup153-nonbound ARGs (3 genes) were Nup153 DEGs (p = 1.47e-09, chi-squared test) (Fig. 5F, Table S3). Conversely, 89.5% of Nup153-regulated ARGs were directly bound by Nup153 (43/48 genes) (Fig.2J), supporting the idea that changes in gene expression upon Nup153 depletion are directly regulated by Nup153-binding to the gene locus. Consistently, Nup153-bound ARGs exhibited stronger gene de-repression than other genes (Fig. 5G, other genes vs. Nup153-bound ARGs, p < 2.2e-16, Wilcoxon Rank sum test; Cohen’s d = 0.65, 95% CI [0.51, 0.70]; Nup153-nonbond ARGs vs. Nup153 bound ARGs, p = 0.0017, Wilcoxon rank sum test; Cohen’s d = −0.26, 95% CI [-0.49, −0.02]). Direct bidirectional regulation of chromatin accessibility was also observed for Nup153-bound DAPs associated with DEGs (Fig. 5H, 5I, Table S3, p = 0.0006, chi-squared test). Consistent with the changes in gene expression, chromatin accessibility was more open at Nup153-bound DEGs in Nup153-depleted cells compared to other Nup153-nonbound ATAC peaks (Fig. 5J, other ATAC peaks vs. Nup153 bound peaks, p < 2.2e-16, Wilcoxon rank sum test; Cohen’s d = 0.19, 95% CI [0.14, 0.23]; Nup153 non-bound peaks vs. Nup153 bound peaks, p = 0.0012, Wilcoxon rank sum test; Cohen’s d = −0.07, 95% CI [-0.02,-0.14]), indicating that Nup153 regulates chromatin accessibility around its target genes.

**Fig. 5.**
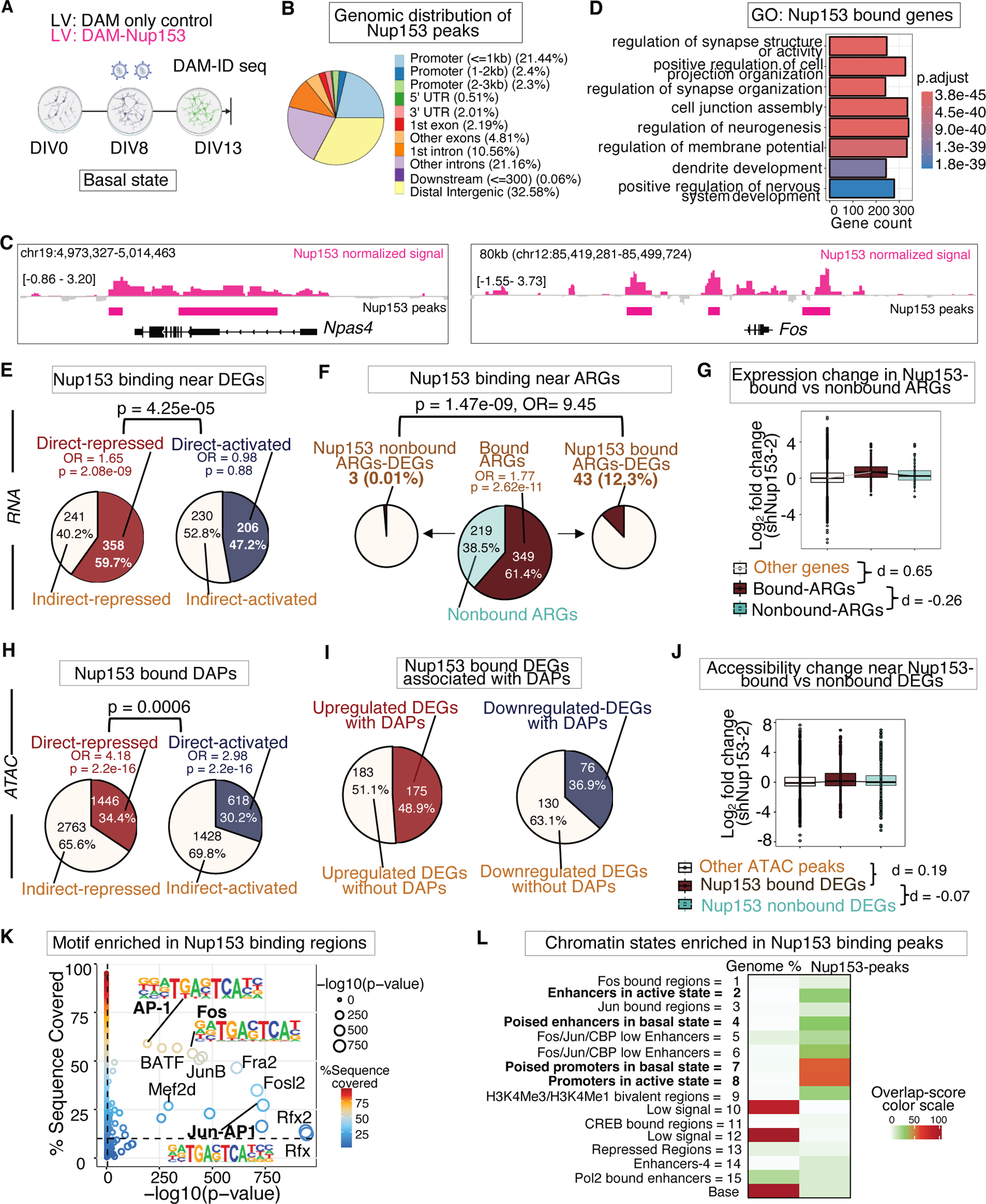
Direct chromatin regulation by Nup153 in basal neurons. (A) A schematic illustration of the experimental plan for DAM-ID in basal neurons. (B) Distribution of normalized DAM-Nup153 peaks across the genome. (C) Visualization of normalized DAM-Nup153 signals (pink) around the Fos in basal neurons alongside ATAC-sequencing (orange) and RNA-sequencing (red) tracks in shControl- and shNup153-2 treated samples, via the IGV genome browser. Reproducible DAM-Nup153 peaks are highlighted with pink boxes underneath the IGV track. (D) GO analysis for all Nup153-bound genes. (E) The proportion of Nup153-DEGs directly repressed or activated by Nup153 (p = 0.425e-05, chi-squared test). (F) The proportions of ARGs bound by Nup153 (middle) and the proportion of ARGs-DEGs within the Nup153-bound ARGs (p = 1.13e-12, chi-squared test). (G) Comparison of the fold change in the expression of Nup153-bound and Nup153-nonbound ARGs, and of all other genes, upon Nup153 depletion. (H) The proportion of DAPs directly repressed or activated by Nup153 (p = 0.0006, chi-squared test). (I) The proportion of upregulated or downregulated Nup153-DEGs associated with DAPs. (J) Comparison of the fold change in chromatin accessibility near Nup153-bound and Nup153-nonbound DEGs, and all other genes upon Nup153 depletion. (K) HOMER-based motif analysis of Nup153 bound regions. (L) The heatmap shows the overlap enrichment of Nup153-bound peaks with ChromHMM states in basal neurons. The scale reflects the proportion of Nup153-bound peaks annotated as the corresponding ChromHMM states in basal neurons. See Fig. S10 and Table S4 for the extended data and supplementary table respectively for Fig. 5. Data for Fig. 5G & 5J are presented as boxlplots, with the connecting lines to the median of each group.

To probe the potential factors involved in Nup153-binding regions, we performed a motif analysis on Nup153-bound regions, which revealed an enrichment of AP-1 motifs at these regions as expected (Fig. 5K, S9). This suggests that Nup153 could directly influence the activity of AP-1 factors in the neuronal genome. To infer the direct role of Nup153 in chromatin states, especially in the basal state, the chromatin states defined by ChromHMM analysis were used to categorize Nup153-binding regions (Fig. S9C), and it revealed that Nup153-binding regions were enriched at poised promoters in the basal state (state 7, Overlap-score = 14.25, −log_10_ (p) > 300) and at promoters in the active state (state 8, Overlap-score = 14.35, −log_10_ (p) > 300), in addition to state 2 (Overlap-score = 9.16, −log_10_ (p) = 72.5), state 4 (Overlap-score = 10.03, −log_10_ (p) = 280.75), and state 6 (Overlap-score = 10.51, −log_10_ (p) > 300) (Fig. 5L). These observations indicate that Nup153 preferentially binds active and poised promoters as well as active enhancers.

### Molecular mechanisms underlying epigenetic regulation by Nup153

Our data so far indicated that Nup153 directly regulates chromatin accessibility and neuronal genes including ARGs. Since Nup153 is known to act as a structural platform to recruit transcriptional or epigenetic modulators to regulate cell type-specific gene expression in stem cells (*21, 22, 37, 38*), we hypothesized that Nup153 is associated with epigenetic modulators to cooperatively regulate neuronal genes in neurons. To address this point, we investigated the interaction of Nup153 with known and potential epigenetic regulators of ARGs, including HDAC1, Ncor1, and Brg1 (*18, 39*). A proximity ligation assay (PLA) showed strong interaction signals between Nup153 and HDAC1 compared to the others, indicating that Nup153 preferentially interacts with HDAC1 in primary cortical neurons (Fig. 6A-B, Fig. S11A). Furthermore, high resolution microscopy analyses revealed that Nup153 and HDAC1 signals are overlapped in PCNs (Mander’s coefficient; fraction of Nup153 overlapping with HDAC1 = 0.52 ± 0.03) (Fig. S11B). A co-immunoprecipitation experiment also revealed an interaction between Nup153 and HDAC1, both in HEK293T cells and brain lysate (Fig. 6C, S12A, S12B), indicating that they are part of the same complex. These observations suggest that Nup153 interacts with HDAC1 and may recruit HDAC1 to regulate target genes. Since HDAC1 is known to suppress the expression of *Fos* by binding its functional elements (*18*), we examined whether Nup153 and HDAC1 bind similar regions, and cooperatively regulate target genes. We used chromatin immunoprecipitation sequencing (ChIP-seq) to determine whether HDAC1 binds to the Nup153-binding regions and found that the two proteins bind in spatially overlapping regions (Fig. 6D, S11C, Table S4), and they preferentially co-bind promoters (Fig. S11D). Of all Nup153-DEGs, 45.4% (486 genes) were co-bound by Nup153 and HDAC1 (Fig. 6D), and 61.7% (300 genes) of these were repressed by Nup153 (Table S4). These data indicate that Nup153 recruits HDAC1 to roughly half of its target genes, and that they cooperatively regulate these target genes (Fig. 6D). We also found that HDAC1 bound to the promoters or enhancers of 54.6% (316/568) of ARGs including *Fos*, *Arc, Npas4,* and *Egr1* (Fig. S11C, Table S4), and 69.3% of Nup153-repressed genes (415/599) were bound by HDAC1 (Fig. S11E, Table S4), indicating a strong association between Nup153-repressed genes and HDAC1-bound genes. We then tested whether HDAC1 binding on the functional elements of the *Fos* gene is affected by Nup153 depletion. First, using ChIP-qPCR assay, we validated that the binding of Nup153 to enhancer 4 (e4) and to the promoter of *Fos* was reduced after Nup153 knockdown (Fig. S11F, S11G). Consequently, the depletion of Nup153 reduced the binding of HDAC1 to the *Fos* gene promoter and e4 (Fig. 6E), indicating that Nup153 is indeed required to recruit HDAC1 to the *Fos l*ocus. HDAC1 reduces the level of histone acetylation and thus acts as a modulator of chromatin accessibility and gene repression (*40*). Therefore, one would expect the depletion of Nup153 should increase histone acetylation levels and chromatin accessibility at Nup153-HDAC1 co-bound regions. To test this, we performed ChIP-Seq to analyze the levels of histone H3 acetylation at lysine 27 (H3K27ac) in the basal and TTX-AP5-treated conditions with or without Nup153 depletion. PCA analysis showed consistent differences in H3K27ac between shControl- and shNup153-2-treated samples in both the basal and TTX-AP5-treated conditions (Fig. 6F), indicating that Nup153-depletion changes the pattern of histone acetylation in both a basal activity-dependent and -independent manner. Next, we tested whether changes in accessibility were accompanied by changes in histone acetylation in DAPs associated with activity-dependent and activity-independent DEGs (Fig. 6G, Table S5). Notably, we observed a consistent increase in histone acetylation in DAPs associated with both activity-dependent and activity-independent DEGs in both the basal and TTX-AP5-treated conditions, albeit to a lesser extent (Fig. 6G). This finding corroborates our previous observation that Nup153 depletion leads to epigenetic dysregulation in both basal activity-dependent and activity-independent manners. Furthermore, we observed increased levels of H2K27ac in Nup153-HDAC1 co-bound regions (Fig. S11I, other H3K27ac peaks vs. Nup153-HDAC1 bound peaks, p < 4.31e-07, Wilcoxon rank sum test; Cohen’s d = 0.08, 95% CI [0.04, 0.12]). This supports the idea that Nup153-dependent HDAC1 recruitment is essential for restricting chromatin accessibility in target regions and maintaining the repression of target genes.

**Fig. 6.**
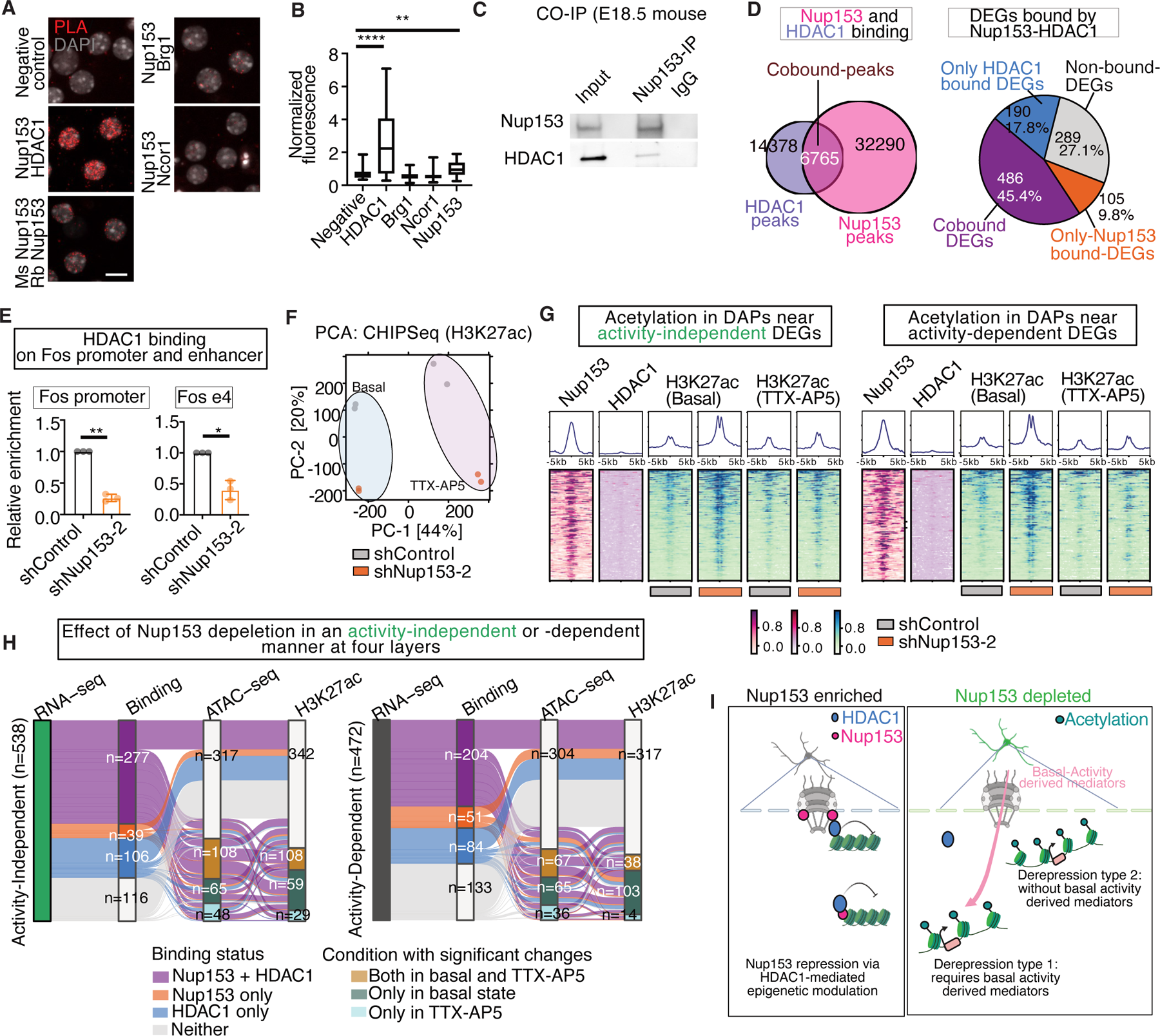
Nup153 cooperates with HDAC1 to regulate ARGs in basal neurons. (A) Max projected confocal images of PLA signals at DIV13 to test the interactions between Nup153 and several epigenetic repressors. Scale bar = 10 μm. (B) Quantification of PLA signals in (A). Normalized fluorescence intensity of the total PLA signals was measured (**p < 0.01, ****p < 0.0001, Kruskal-Wallis test followed by Dunn’s multiple comparison, n = 30 to 150 cells from three independent experiments). (C) Co-immunoprecipitation of endogenous Nup153 and HDAC1 from brain lysate of embryonic cortices (E18.5). (D) (Left) A Venn diagram showing the overlap of DAM-Nup153 peaks with HDAC1 peaks in the basal state. (Right) A pie chart showing the proportion of Nup153-DEGs that are bound by both Nup153 and HDAC1 (brown), only HDAC1 (purple), only Nup153 (pink), or neither HDAC1 nor Nup153 (orange). (E) Knockdown of Nup153 reduced the binding of HDAC1 on the *Fos* promoter and enhancer (e4) regions revealed by ChIP-qPCR (*p < 0.05, **p < 0.01, one-sample t-test. Three independent experiments were conducted). (F) A PCA of H3K27ac-ChIP-seq data from control and Nup153-depleted neurons in the basal and TTX-AP5-treated conditions. (G) Enrichment of DAM-Nup153 and HDAC1 peaks around DAPs near basal activity-independent or - dependent DEGs in the basal state, alongside changes in the enrichment of H3K27ac peaks after Nup153 depletion. The graphs at the top depict the summarized profile plots of the aggregated signals for each corresponding condition. (H) (Right) An alluvial plot showing the connection of basal activity-dependent DEGs with different epigenomic signatures. (Left) An alluvial plot showing the connection of basal activity-independent DEGs with different epigenomic signature. (I) A schematic illustration depicting the Nup153-dependent HDAC1-mediated epigenetic modulation underlying the repression of ARGs. Nup153 interacts and recruits HDAC1 to the regulatory regions of ARGs and cooperatively represses their expression in a basal-activity dependent and -independent manner in post-mitotic neurons. See also Fig. S11, S12, S13 and S14, and Table S4 and S5 for the extended and supplementary data for Fig. 6. Data for Fig. 6B are presented as boxplots with the connecting lines to the median of each data group. The data (Fig. 6E) are presented as mean ± standard deviation (s.d).

Finally, we analyzed the transcriptomic and epigenomic layers together, determining how many basal activity-dependent DEGs and basal activity-independent DEGs are bound by Nup153/HDAC1, as well as how many of them are associated with changes in ATAC and H3K27ac levels in the basal state, the TTX-AP5-treated state, or both (Fig. 6H, Table S6). In summary, out of the 472 activity-dependent DEGs, 21 were bound by both Nup153 and HDAC1, and were associated with changes in the ATAC and H3K27ac layers in both the basal and TTX-AP5-treated states. In contrast, of the 538 activity-independent DEGs, 33 bound by Nup153/HDAC1 showed changes in ATAC and H3K27ac layers in both conditions (Fig. 6H, Table S6). These data suggest that Nup153 can regulate its binding targets via epigenetic mechanisms in both basal activity-dependent and -independent manners, and some of them are regulated by Nup153 in combination with HDAC1.

To analyze all the interactions and determine how Nup153-dependent ARGs are affected at each layer, we analyzed them together (Fig. S13A, S13B, Table S6). Our analysis identified *Nr4a3* as an activity-independent DEG-ARG and *Maml3* and *Map3k5* as activity-dependent DEG-ARGs, bound by Nup153-HDAC1 and affected at the ATAC and H3K27ac layers in both the basal and TTX-AP5-treated states (Fig. S13A-S13C, Table S6). In addition, similar to changes in accessibility, ARGs such as *Fos, Bdnf* and *Npas4*, consistently exhibited an increasing trend in H3K27ac levels in the presence of TTX-AP5 (Fig. S8G). These observations support the idea that Nup153 regulates their target genes with HDAC1 at three layers. Nup153 depletion leads to changes in HDAC1-mediated histone acetylation and/or in chromatin accessibility, and ultimately in RNA expression. However, these three layers are further influenced by basal neuronal activity, with the extent of the influence varying depending on target genes. We summarized the Nup153-dependent HDAC1-mediated epigenetic modulation in mature neurons in Fig. 6I.

To more comprehensively evaluate regulators of both basal activity-dependent and -independent ARGs we also performed DecoupleR (*88*) based analysis of transcription factor activity (Fig.S14, see methods). Our analysis identified distinct transcription factors associated with basal activity-dependent and basal activity-independent ARGs, supporting our hypothesis that Nup153 depletion leads to ARG dysregulation through two separate mechanisms. Intriguingly, several regulators enriched among basal activity-dependent ARGs, including REST and E2F1, have been reported to form complexes with HDAC1 in both neuronal and non-neuronal contexts (Fig. S14A, S14B) {Qiu, 2008 #1469;Huang, 1999 #1861;Greene, 2004 #1862;Telles, 2012 #1863}. These findings further support the idea that Nup153 depletion affect ARGs that are directly repressed by HDAC1-containing complexes under basal conditions In contrast, for basal activity-independent ARGs, the transcriptional factors were mainly positive transcriptional regulators following Nup153 depletion, including CTNNB1, PITX2, CREB1 (Fig. S14C, S14D). This pattern suggests that Nup153 depletion allows positive regulators to upregulate their target genes independent of basal activity. Overall, our analysis suggests that Nup153 coordinates the repression of key ARGs under basal conditions, likely through cooperation with other repressive complexes or by directly restricting positive regulators. In this context, Nup153, together with HDAC1 and other molecular factors, may play an important role in preventing imprecise gene induction and thereby preserving neuronal responsiveness.

### Impact of Nup153-depletion on neuronal network development

Our data demonstrated that Nup153 regulates neuronal responsiveness through epigenetic modulation. Since Nup153-regulated DEGs, including ARGs, play a central role in neural network plasticity, we next examined whether Nup153 depletion leads to changes in neural network activity using a multi-well microelectrode array (MEA) platform as described previously (Fig. 7A)(*41*). We observed increased bursting events in Nup153-depleted neurons (Fig. 7B-D), and the number of spikes was also higher in Nup153-depleted neurons (Fig. 7E-F). These data indicate that Nup153 depletion leads to increased neuronal excitability. As hyperexcitability is often associated with neurodevelopmental disorders, we analyzed the association of Nup153-DEGs with genes related to human neuropsychiatric disorders and behavioral traits using the MAGMA gene-set analysis (*42*). Nup153-DEG related gene-sets were significantly associated with bipolar disorder (p = 0.0008 for all Nup153-repressed genes and p = 0.0378 for direct Nup153-repressed genes), obsessive compulsive disorder (OCD) (p = 0.0259 for direct Nup153-repressed genes) and insomnia (p = 0.0026 for all Nup153-repressed genes and p = 0.0112 for direct Nup153-repressed genes) indicating the potential involvement of Nup153-dependent epigenetic mechanisms in psychiatric disorders (Fig. 7G, Table S9). Moreover, GSEA indicated positive enrichment for depression- and schizophrenia-associated genes in Nup153 depleted neurons (enrichment for genes related to depression: p = 0.02, NES = 1.52; enrichment for genes related to schizophrenia: p = 0.03, NES = 1.24) (Fig. 7H, Table S9), which highlights the disease relevance of Nup153-regulated neuronal gene programs.

**Fig. 7.**
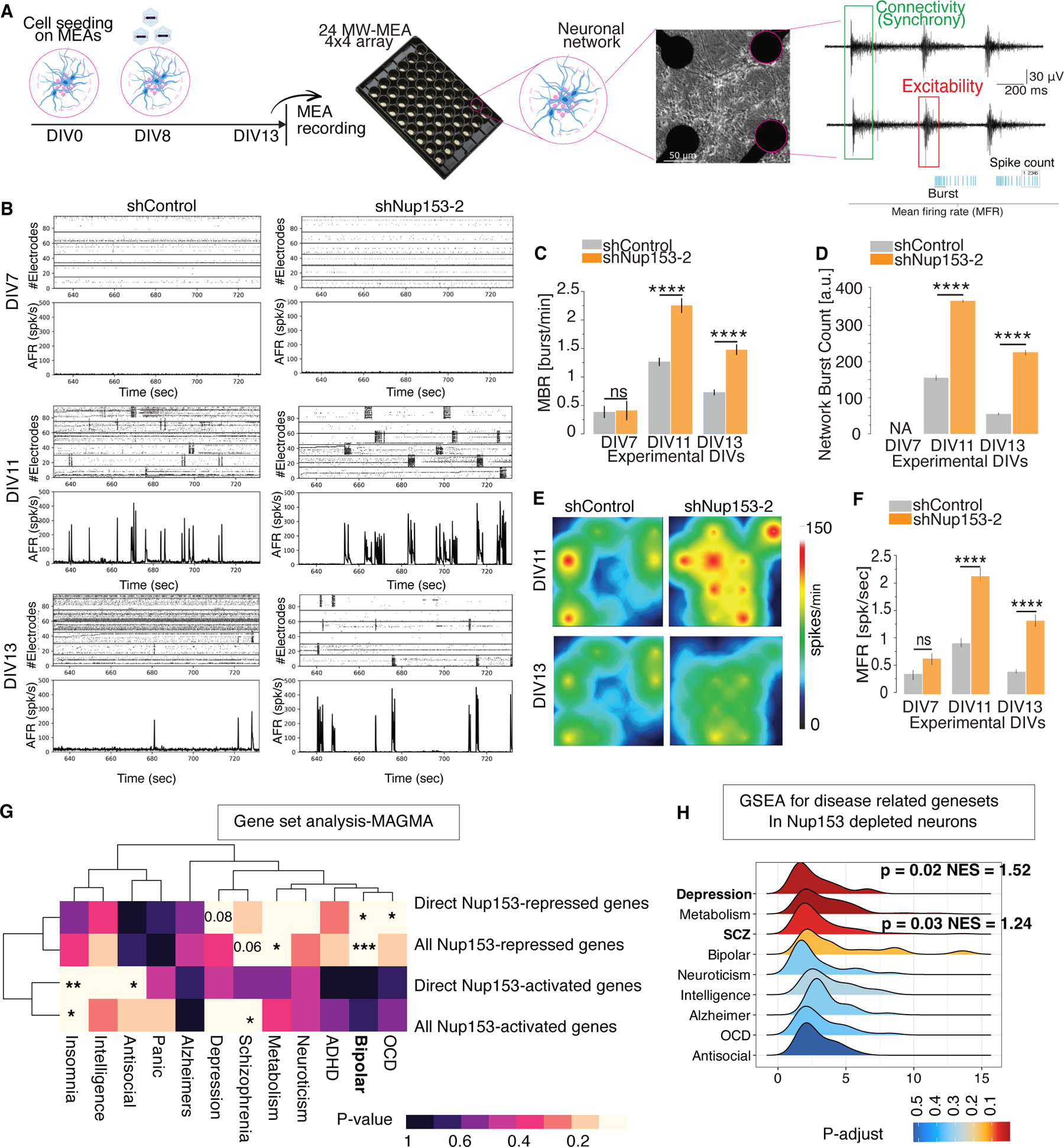
Impact of Nup153-depletion on neuronal networks and its relevance to disease. (A) An overview of the 24-well CytoView MEA plate setup and the experimental workflow. The Fig. includes exemplary spiking activity and annotations of extracted electrophysiological features, including the detection of synchronous firing patterns essential for identifying alterations in network dynamics. (B) Raster plot representations exhibit the development of network burst synchrony from DIV7 to DIV13 in shControl- and shNup153-2-treated neurons. (C) Quantification of mean burst rate per minute (MBR) during neuronal maturation (****p < 0.0001 for shControl vs shNup153-2, one-way ANOVA). (D) Quantification of Network burst count from control and shNup153-2-treated neurons (****p < 0.0001, one-way ANOVA). (E) Topographical pseudo-color maps displaying firing patterns at DIV11 and DIV13, showing changes in active neuronal network activity dynamics. (F) Quantification of mean spike rate from control and shNup153-2-treated neurons (****p < 0.0001, one-way ANOVA). (G) A heatmap showing the p-value obtained from competitive gene-set analysis for Nup153-DEGs related gene-sets for different neuropsychiatric disorders and cognitive traits. *p < 0.01, **p < 0.001 & ***p < 0.000,1 according to MAGMA based null hypothesis testing. (H) A ridgeplot showing the enrichment of gene-sets associated with different diseases in Nup153-depleted neurons in the basal state. The color scale visualizes the GSEA analysis-based p-values for the respective disease related gene-set. The data (Fig. 7C, 7D & 7F) are presented as mean ± standard error of the mean (s.e.m).

## DISCUSSION

### Neuronal responsiveness directed by nucleoporin-mediated epigenetic regulation

Our findings revealed an unexpected role of Nup153 in balancing neuronal responsiveness by regulating the chromatin state of neuronal genetic elements via basal activity-dependent and -independent mechanisms. The induction of neuronal genes including ARGs is a crucial process for neuronal plasticity, as it allows synaptic inputs to be translated into gene expression. Because this process is essential for circuit reorganization, learning, and memory, understanding the mechanisms underlying the induction of neuronal genes in response to neuronal activity has been a central question in neuroscience for the last few decades (*1, 43*). Thus far, several transcriptional and epigenetic mechanisms have been identified (*44*), and impairments in these mechanisms have been strongly linked to neurodevelopmental disorders as well as cognitive deficits in neurological disorders (*45–47*). However, compared to the mechanisms underlying the activation of neuronal gene programs, the mechanisms underlying the repression of them in basal neurons and their coordinated induction upon neuronal activation have remained largely elusive. These mechanisms likely ensure a high signal-to-noise ratio of neuronal gene expression including ARGs allowing neurons to encode spatiotemporal information with greater precision. Our results revealed that Nup153 directly binds to target neuronal genes including several ARGs and facilitates the organization of chromatin to prime their epigenetic states and expression in post-mitotic neurons, and depletion of Nup153 bidirectionally dysregulates these genes in both a basal activity-dependent and activity-independent manner. We have shown that Nup153 regulates chromatin accessibility at DEG-associated regulatory elements and recruits HDAC1 to these regions, acting as a structural platform to organize nuclear architecture including the repressive complexes and the chromosomes themselves (Fig. 6I). Taken together, our findings highlighted that Nup153 directly regulates chromatin states and neuronal responsiveness through chromatin accessibility and histone modifications. In addition to chromatin regulation, our data also suggest that Nup153-dependent DEGs are involved in the regulation of network excitability (Fig. 7). It will be intriguing to investigate how Nup153 levels are regulated in neurons to balance neuronal responsiveness at different epigenetic levels, and whether Nup153-regulated neuronal states dictate neuronal functions such as cellular memory or memory engrams.

Recent research has revealed the diverse functions of nucleoporins in genomic processes. Nucleoporins have been found to bind to promoters and super-enhancers, thereby regulating cell type-specific epigenomic programs in a bimodal manner (*21, 25, 37, 38*). Additionally, the nuclear pore complex plays a critical role in coordinating 3D genome interactions, which are important for cellular memory and responses to growth stimulation (*48–50*). Interestingly, although nucleoporins have been shown to be essential for the activation of immediate early genes in embryonic stem cells and T cells (*48, 50*), in this study, we observed that ARGs are repressed by Nup153 in post-mitotic neurons. These findings suggest that nucleoporins may have distinct roles in different cellular contexts and recruit distinct co-factors to regulate cellular responses in a cell type-specific and cellular state-dependent manner. Moreover, Nup153 binds both promoters and distal elements in the neuronal genome (Fig. 5B), and chromatin states are regulated in a basal activity-dependent and -independent manner (Fig. 4). It would be of great value to identify the common and cell type-specific co-factors of nucleoporins to understand their function as a structural platform for epigenetic regulation and to examine the different types of Nup153-dependent regulation of chromatin states in promoters and enhancers in the future.

### The nuclear pore complex as an organizer for the dynamic regulation of neuronal genes in post-mitotic neurons

Since Blobel proposed the gene gating hypothesis (*51*), the nuclear pore complex has been studied as a site for active transcription, chromosome positioning, chromatin interaction, and DNA repair (*23*). Recent studies further extended this notion, and the nuclear pore complex has been shown to coordinate developmental processes by regulating developmentally poised genes or by stabilizing the expression of lineage-specific genes (*21*). Although these studies uncovered critical roles of nucleoporin-directed nuclear architecture in gene regulation, most of them focused on developmental decisions such as proliferation, differentiation, and lineage specification. Therefore, although the nuclear pore complex is considered to be one of the most stable protein complexes in post-mitotic cells, its function in genomic regulation in long-lived post-mitotic cells, such as neurons, has not yet been explored. Our current study shows that Nup153 works as an interface to integrate cellular signaling into neuronal gene regulation. Importantly, in post-mitotic functional neurons, most cellular signaling, including Ca2^+^ and MAPK signaling, has to pass through nuclear pores and must activate target genes within a few minutes to precisely encode synaptic information into gene expression. To robustly organize this dynamic process, it is plausible that neuronal nuclei possess a structural basis for the induction of neuronal genes including ARGs upon neuronal activation. Our results suggest that nucleoporins could work as an epigenetic regulator to maintain the chromatin states of neuronal genes including a subset of ARGs in a poised state. This allows Nup153-associated chromatin regions to be easily regulated by incoming signals. Indeed, it has recently been shown that several ARG loci are located at the nuclear periphery, and there is a biochemical compartment beneath the nuclear periphery that induces ARG expression and neuronal activity-dependent DNA damage to the genomes (*52*). The nuclear pore is also known as a DNA repair site (*23*), and HDAC1 is involved in DNA damage repair (*53, 54*). Therefore, it would be intriguing to investigate how various molecular mechanisms, including transcriptional induction of ARGs, neuronal activity-induced DNA damage, and DNA repair, are coordinated by nucleoporins. Investigating the dynamic spatiotemporal interaction between nucleoporins and chromatin would be of interest to unravel the contribution of nucleoporins in the regulation of activity-dependent gene expression. Furthermore, understanding these processes could be instrumental for gaining a clear understanding of neuronal plasticity and relevant neuronal pathology. In fact, nuclear pores have been identified as a key target of neurological pathology including frontotemporal dementia, amyotrophic lateral sclerosis, and Alzheimer’s disease (*55–60*). Class I HDACs, including HDAC1, are also indicated as a potential regulator of learning, memory, and age-associated gene dysregulation (*53, 61*). Therefore, investigating how the disorganization of nucleoporin-directed nuclear architecture and their complexes impair the regulation of the neuronal genome in pathological conditions would be crucial.

### Limitations of the study

Although our data showed that Nup153 plays a critical role in regulating the neuronal epigenome and neuronal responsiveness, not all DEGs are co-bound by Nup153 and HDAC1. In order to fully understand the mechanisms of Nup153-dependent epigenetic modulation, it is crucial to study the Nup153 interactome in neurons. Furthermore, the localization of Nup153, its cofactors, and their target genome loci are of great importance. Nup153 could be localized in the nuclear interior, and the spatiotemporal dynamics of Nup153-dependent epigenetic regulation must be investigated to understand its role in time-sensitive gene regulation.

## MATERIALS AND METHODS

### Primary cortical neurons (PCNs)

All procedures related to the care and treatment of the mice were approved by the Government of Saxony and performed according to their guidelines. Embryonic cortices (embryonic day 17.5-19.5) (C57/BL6 mice) from one litter were dissected and pooled together to be processed as a single biological replicate. A Neural Tissue Dissociation Kit (Miltenyl Biotec) was used to dissociate tissue according to the manufacturer’s instructions. Cell culture dishes were pre-coated with Poly-Dl-Ornithine Hydrobromide (50 µg ml^−1^) (Sigma Aldrich) overnight at 37°C, washed three times with dH20, and allowed to dry at room temperature (RT) for an hour as described previously (*41*). PCNs were then plated at an appropriate cell density in Neurobasal medium (Life Technologies) supplemented with 2% B-27 supplement (Life Technologies), Penicillin/ Streptomycin (10U ml^−1^) (Life Technologies), and 1% FBS serum (Sigma Aldrich). PCNs were grown in incubators maintained at 37°C with a CO_2_ concentration of 5%. 30% medium was replaced at DIV4 and DIV8 with Brain Phys Neuronal Culture Medium (STEMCELL Tec), supplemented with 2% SM1-NC (STEMCELL Tec).

For most experiments, PCNs were harvested at DIV13 unless otherwise noted. For RNA-Seq/qPCR experiments, PCNs were plated at a cell density of 1000-1100 cells/mm^2^ in a 12-well plate. For each replicate of a ChIP-qPCR experiment, the PCNs were plated at a higher cell density of 1200-1300 cells/mm2 (7-8 million cells per plate) on a 10 cm plate (Corning). For the proximity ligation assay (PLA), PCNs were plated at a cell density of 1500-2500 cells/mm2 (50,000-80,000 cells per well) in a 10-well chamber (Greiner). For fluorescence *in situ* hybridization (FISH), PCNs were plated at a cell density of 1500-1600 cells/mm2 (1.5 million cells per well) in a 6-well plate (Costar) together with an 18 × 18 mm glass coverslip (Marienfeld). For immunocytochemistry, PCNs were plated at a density of 600-700 cells/mm2 (100,000-125,000 cells per well) in a 24-well plate (Costar) with a 13 × 13 mm glass coverslip (Marienfeld). In most experiments, PCNs were treated with adeno-associated viruses (AAVs) expressing shNup153-1/2 or shControl at DIV8 and collected for experiments at DIV13. For experiments using lentiviruses (LVs) carrying CRISPRa constructs, LVs were applied at DIV8, the media was completely replaced with BrainPhys medium supplemented with 2% SM1-NC (STEMCELL Tec) at DIV10, and the treated PCNs were collected at DIV13 for further experiments.

### Neural progenitors and astrocytes

#### NeuPC culture

The NeuPC line from the E15.5 embryonic cortex (C57/BL6) was isolated and cultured as described previously with minor modifications (*41*). The cell culture dish was pre-coated with Poly D-lysine (10 μg/ml, Sigma Aldrich) at room temperature (RT) overnight and then incubated with DMEM-F12 (Life Technologies) containing laminin (0.5 μg/ml) (Roche) at room temperature overnight. NeuPCs were cultured in pre-coated plates with DMEM/F-12 (Life Technologies) supplemented with 1% N2 and 1% B27 (Life Technologies) in the presence of FGF2 (10 ng/ml, PeproTech), EGF (10 ng/ml, PeproTech), and heparin (5 μg/ml, Th.Geyer). 30% of the cell medium was replaced with fresh cell medium every second day. NeuPCs were passaged to new wells if cell confluency exceeded 80%.

#### Astrocyte culture

The mouse NeuPCs were plated on a pre-coated dish with Poly D-lysine (10 μg/ml, Sigma Aldrich) and laminin (0.5μg/ml, Roche). The cells were cultured in DMEM/F-12 supplemented with 1% N2 and 1% B27 (Life Technologies) in the presence of 0.5% fetal bovine serum (FBS, Sigma) for 5-7 days. Astrocyte identity was verified using antibody staining for the known astrocyte marker GFAP.

### Knocking down Nup153 with AAVs

To knockdown Nup153, shRNAs targeting mouse Nup153 (TRCN0000102395 and TRCN0000102398) were cloned in a PX552 AAV vector (Addgene). AAVs with the AAV2/8 serotype were prepared by the viral core facilities at the Salk institute or Charité-Universitätsmedizin Berlin. The final titer of AAVs used for shControl, shNup153-1, and shNup153-2 was 1.2e+10 GC/ml.

### Pharmacological treatments

PCNs were either pre-silenced using tetrodotoxin (TTX,1-2 µM, hellobio) on DIV12 for 24 hours, and then D-AP5 (100 µM, Tocris) was applied for six hours before collection.

### Immunohistochemistry

Tissue sections for immunostaining were prepared as described previously (*62*). Briefly, animals were euthanized by sodium pentobarbital and then transcardially perfused with PBS followed by 4% PFA in PBS. For Nup153 staining, brains were postfixed at 4°C for 6-8 hr and then transferred to 30% sucrose-PBS. Forty-µm thick coronal sections were cut on a sliding microtome (Leica) and subjected to immunohistochemistry as described previously (*62*). Briefly, brain sections were blocked with 0.3% Triton X-100 in PBS containing 3% horse serum for 60 minutes at RT. After blocking, tissues were incubated with primary antibodies in blocking buffer for two nights at 4°C. Sections were washed three times with 0.3% Triton X-100 (Sigma) in PBS (PBS-T) and incubated with fluorophore-conjugated secondary antibodies (Jackson Immunoresearch) with DAPI for two hours at the room temperature (RT). Sections were washed three times with PBS-T and mounted with Moviol (Sigma-Aldrich).

### Immunocytochemistry

Cells attached to 13 mm x 13 mm glass slides (Marienfeld) were fixed with either 2% or 4% PFA (EMS) in PBS for 10 mins at RT depending on the antibody used. To quench residual PFA and to reduce background, cells were washed with PBS containing 0.1M glycine (Carl Roth) for 5 mins at RT, and two times with normal PBS for five minutes at RT. Subsequently, cells were blocked with 3% horse serum and 0.1% Triton X-100. After blocking, cells were incubated with primary antibodies for 2hrs at RT and washed three times with PBS-T. Cells were then incubated with fluorophore-conjugated secondary antibodies (Jackson Immunoresearch) for 60 minutes at RT, washed three times with PBS-T and mounted with Prolong Diamond Antifade mounting medium (Life Technologies).

### Ethyl Uridine (EU) labelling

5-Ethynyl-uridine (EU; Jena Bioscience) was applied to PCNs at DIV13 at a final concentration of 250 µM for six hours. Then, PCNs were fixed with 4% PFA/PBS for 10 mins at RT, followed by three washes with PBS. EU was visualized using a Click-iT™ Plus Alexa Fluor™ 555 Picolyl Azide Toolkit (Life Technologies) according to the manufacturer’s instructions, and PCNs were mounted with Prolong Gold anti-fade reagent (Invitrogen).

### Imaging and quantification for immunostainings

Immunostained samples were imaged using a Zeiss LSM 980 with a 20x objective (Plan, Apochromat, NA = 0.8) or a 63x objective (C-Plan, Apochromat, NA = 1.40). An identical protocol was used to quantify expression levels of nuclear staining. To remove background, the mean grey intensities of five different regions without cells + 3 × standard deviation were subtracted from all images of the Z-stack. To quantify nuclear signals, DAPI-positive nuclei were segmented using the Stardist program followed by creating a mask and thresholding using Otsu’s method in ImageJ (*63*) or CellProfiler (v4.2.1) (*64*). To identify AAV-treated cells, AAV-derived KASH-GFP signals were also segmented using Stardist. To identify KASH-GFP/DAPI positive nuclei, the image calculator function was used to subtract the inverted DAPI mask from the KASH-GFP mask. In CellProfiler a combination of DAPI and neuronal markers such as NeuN/Map2 was employed for nuclei segmentation using Otsu’s method or Cross-entropy method with additional shape- and size-related filters to remove non-nuclear signals. In the Fiji pipeline, the Analyze Particle function was used to identify circular objects in the mask of KASH-GFP/DAPI positive nuclei. The mean grey intensity or the integrated fluorescence intensity (IFI) of GFP/DAPI double-positive nuclei was measured. To correct for batch variation, the fluorescence intensity of all nuclei in control and shNup153-1/2 cells was divided by the average fluorescence intensity of all nuclei in the control condition. In CellProfiler, the mean integrated intensity of background-corrected channels within each segmented object was calculated. CellProfiler based analysis was used for Fig.1C-D, but for the rest of the nuclear staining quantifications, an Image-J based protocol was applied.

For high resolution imaging in Fig. S11B, immunostained PNCs were imaged with Zeiss LSM 880 Airyscan microscope with a 63x objective (C-Plan, Apochromat, NA = 1.40). To assess overlap between Nup153 and HDAC1 signals, Mander’s coefficient was quantified using JACoP plugin in ImageJ(*65*) (*63*).

### Proximity Ligation Assay (PLA)

Cells were fixed with 2% PFA (EMS) in PBS for 10 min at RT. To quench residual PFA and reduce background, cells were washed with PBS containing 0.1M glycine (Carl Roth) for 5min at RT, and twice with normal PBS for 5 min each. Then, cells were incubated in blocking solution consisting of 5% horse serum (VWR International) and 0.1% Triton X-100 (Sigma) in PBS. Cells were incubated with primary antibodies at RT for one hour, washed with PBS-T, and incubated with corresponding Duolink *in situ* PLA probes (Sigma) diluted in blocking solution for one hour at 37° in a humid chamber. They were washed twice for 5 min with Duolink wash solution (Sigma). The ligation and rolling circle amplification reactions (Sigma) were performed according to the manufacturer’s protocol. Cells were mounted using Duolink *in situ* mounting medium with DAPI (Sigma) and imaged using a Zeiss LSM 980 with a 40x objective.

### Co-immunoprecipitation (CO-IP)

HEK293T cells were transfected with plasmids P181 pK7-HDAC(GFP) (*66*) and/or pLVXEP-FLAG-Nup153-mCherry using Polyethylenimine (PEI, Polyscience Europa). Transfected cells were lysed using RIPA buffer (10mM Tris/Cl ph7.5, 150mM NaCl, 0.5mM EDTA, 0.1% SDS, 1% Triton^TM^ X-100, 1% deoxycholate, 0.09% sodium azide) for 30min at 4 °. Lysed cells were centrifuged at 16000G for 10 minutes at 4 ° and supernatant was collected. 5% of lysate was separated as input material. Remaining lysate was incubated with 20µl GFP-trap magnetic agarose (ChromoTek, gmtak-20) or RFP-trap magnetic agarose (ChromoTek, rtma) at 4 ° overnight. Following this the beads were processed as per manufacturer’s instruction. In brief, beads were washed with wash buffer twice and eluted via 2X laemmeli sample buffer (BioRad). The eluted samples were denatured at 95 ° for 10 minutes and subjected to western blot. Briefly, Denatured samples were separated in a 7.5% Mini-PROTEAN TGX^TM^ precast protein gel (BioRad) and transferred to a 0.2µm PVDF membrane (BioRad). The membrane was blocked using 5% skim milk and incubated with primary antibodies overnight at 4 °. Following this membrane was washed, incubated with HRP-conjugated secondary antibodies and signal was developed using chemiluminescent detection kit SuperSignal West Dura or Pico PLUS (Thermo Fischer).

#### Endogenous HDAC1 and Nup153 interaction

Adult (single 14-week-old) or embryonic (six pooled hemispheres) mouse brain tissue was harvested and dissociated on ice in Lysis Buffer (50 mM Tris-HCl pH 7.5, 30 mM NaCl, 120 mM KCl, 2 mM MgCl₂, 1.125 mM ZnCl₂, 1 mM DTT, 1 mM EDTA, 10% glycerol, 0.05% CHAPS, 0.15% Emphigen BB) supplemented with cOmplete EDTA-free protease and PhosSTOP phosphatase inhibitors (Roche). The tissues were then Dounce-homogenized (adult: 15 strokes with pestle A, 20 strokes with pestle B, then 70 µm filtration; embryonic: 10 strokes with each pestle), incubated on ice (embryonic, 15 min; adult, 40 min), treated with 150 U DNaseI (NEB) for 20 min at room temperature with rotation, and cleared at 20,000 ×g for 10 min at 4 °C. For co-immunoprecipitation, supernatants were split into equal experimental and control aliquots (embryonic samples first diluted 1:1 in lysis buffer) and incubated overnight at 4 °C with 5 µg anti-Nup153 (QE5, Abcam ab24700) or mouse IgG control (CST 5415S). 50 µl of pre-washedDynabeads Protein G (Thermo Fisher) were then added and rotated for 2.5 h at 4 °C, after which the beads were washed three times with 300 µl chilled lysis buffer, and the bound proteins were eluted in 40 µl elution buffer (Thermo Fisher 14321D). For Western blotting, input and IP eluates were combined with NuPAGE LDS sample buffer and reducing agent, heated at 70 °C for 10 min, resolved on NuPAGE 3–8% Tris-Acetate gels (Invitrogen), transferred to PVDF (Trans-Blot Turbo, Bio-Rad), blocked in StartingBlock (Thermo Fisher) for 35 min at room temperature, and probed overnight at 4 °C with primary antibodies in Can Get Signal Solution 1 (Toyobo). Following this, the membrane was washed and incubated with HRP-conjugated secondaries and developed with SuperSignal West Femto substrate (Thermo Fisher).

### Overexpression of Nup153 by CRISPRa/dCas9

To perform overexpression of Nup153, a CRISPR/dCas9 based strategy optimized for neurons was adopted (*28*). sgRNA specific for the Nup153 promoter was designed using the online tool CHOPCHOP. The sequence for the sgRNA targeting a scrambled sequence was derived from a previously published report (*67*).

#### Following oligos were used

Nup153-sgRNA: 5’ CACCGACGACTGCAGGGACGACGA 3‘ (sense), 5‘ AAACTCGTCGTCCCTGCAGTCGTC 3’ (antisense)

Scrambled-sgRNA: 5’ CACCAACCCCTGATTGTATCCGCA 3‘ (sense), 5’ AAACTGCGGATACAATCAGGGGTT 3’ (antisense)

The sgRNAs we designed were cloned into Addgene plasmid #11419 according to the strategy described previously (*68*). Briefly, sgRNA oligos were phosphorylated and annealed with T4 PNK(NEB) according to the manufacturer’s protocol. A simultaneous digestion of Addgene plasmid #11419 and a ligation reaction with sgRNA oligos was set up. Digestion was performed using Bbs1 enzyme (NEB), and ligation was performed using T7 DNA ligase (NEB). Successful cloning of sgRNA results in mutation of the Bbs1 site at the site of sgRNA insertion, so transformed colonies were screened by restriction digestion with Bbs1 and Xho1 to verify successful sgRNA insertion.

#### Lentivirus production for the CRISPRa and DAM-ID system

HEK293T cells were used to generate lentiviruses according to a protocol previously described. Briefly 6 × 10 cm plates with 50% confluent HEK293T cells were treated with specific CRISPR and packing plasmids using Polyethylenimine (PEI, Polyscience Europa). The media was then completely replaced with DMEM supplemented with 10% fetal bovine serum (FBS) (Sigma Aldrich) for virus production. Media containing viral particles were collected after three days of incubation. Viral particles were enriched by centrifugation or by LentiX^TM^ (Takara Bio), and stored at −80°C. Infection of PCNs with DAM-ID based lentivirus was performed at DIV8. Samples were collected at DIV13 for further processing. To improve efficiency of lentivirus transduction, PCNs were centrifuged at 300G for 5 minutes at room temperature. Further, entire media was changed next day to reduced lentivirus-based toxicity.

### ChIP-qPCR and ChIP-Sequencing

Chromatin immunoprecipitation (ChIP) was performed as previously described with minor modifications (*21, 49*). Briefly, PCNs were collected in the basal state at DIV13. PCNs were fixed by adding 16% formaldehyde (VWR International) directly to the media to a final concentration of 1% for 10 minutes at RT. Residual formaldehyde was quenched by adding 1M glycine (Carl Roth) directly to the media to a final concentration of 0.1M for 15 minutes at RT. After quenching, PCNs were washed twice with cold PBS solution containing protease inhibitor (Sigma Aldrich) (PBS-PI) for five minutes each. Cells were scraped from the plates and collected in a 1.5ml tube, pelleted by centrifugation at 300 G, and washed with PBS-PI. Samples were either stored at −80°C or processed directly for chromatin immunoprecipitation.

For ChIP-Seq of HDAC1 and H3K27ac, 1.5mM EGS (ethylene glycol bis succinimidyl succinate) was applied for 15 minutes at RT prior to formaldehyde fixation, followed by two 5-minute washes with PBS. A total of two biological replicates were used for ChIP-Seq for each condition. Samples were thawed and lysed in 1 ml of ChIP lysis buffer for 10 min at 4°C on a rotator. Samples were washed in ChIP Wash buffer for 10 min at 4°C and resuspended in ChIP buffer for sonication. Samples were sonicated using a focused-ultrasonicator (Covaris).

After confirming the size of the fragments, the samples were pre-cleaned with 10 µl of Dynabeads Protein G (Life Technologies) for 1 hour at 4°C, and then the samples were incubated with 2 ug of primary antibodies at 4°C overnight on a rotator. The next day, Dynabeads were added into the samples for 2 hours, and the tubes were rotated at 4°C. Dynabeads were separated from the samples using a magnetic stand DynaMag^TM^ (Thermo Fisher). The beads were sequentially washed with low-salt buffer, high-salt buffer, lithium chloride buffer and twice with TE 1X buffer (IDTE). The bound chromatin was eluted from the Dynabeads by incubating it in 100 ul of elution buffer. Following RNase A and proteinase K treatments, DNA was purified using a standard phenol chloroform extraction method. For ChIP-qPCR, the list of qPCR primers is shown in the table S10. Each pair of primers was tested, and only pairs of primers with an efficiency of 90% to 105% was selected. ChIP-qPCR was performed using iTAQ SYBER supergreen mix (BioRad).

### ChIP-seq library preparation

ChIP samples were subjected to Illumina fragment library preparation using the NEBnext Ultra II DNA library preparation chemistry (New England Biolabs, E7645L). Briefly, DNA fragments were end-repaired, A-tailed and ligated to unique-dual indexed Illumina Truseq adapters (IDT, prehybridized UDI-UMI Adapters for Illumina, 0.15 µM adapter per reaction). The resulting libraries were PCR-amplified for 15 cycles using universal primers (primer 1: CAAGCAGAAGACGGCATACGAGAT and Primer 2: AATGATACGGCGACCACCGA*G; *: Phosphothioate bond) and purified using XP beads (Beckman Coulter). The libraries were size-selected with XP beads. The resulting libraries were checked for quality and quantified using a Fragment Analyzer (Agilent) NGS Fragment Kit (1-6000bp). The final libraries were subjected to 100bp paired-end sequencing on the Illumina NovaSeq6000 platform at a depth of 35-55 million fragments per library.

### DAM-ID

#### Cloning

The pCDH-mCherry-i4Dam plasmid was constructed by Gibson assembly into pCDH-EF1, using mCherry as an upstream open reading frame, and Dam from Addgene plasmid #59217 with a C-terminal Myc-tag (*69*). An intron was inserted into Dam by ligating oligos with a modified synthetic intron (‘IVS’) from (*70*) into the BamHI restriction site within Dam, between the 3rd and 4th helix of the DNA-binding domain of the Dam methylase (*71*). pCDH-mCherry-i4DAM-Nup153 plasmid (DAM-Nup153) was cloned using Gibson assembly (NEB E5510) of PCR-amplified Nup153 into the Sal1 restriction site at the C-terminal of DAM. DAM-Nup153, DAM-only control and lentivirus packaging plasmids were produced in dam^-^/dcm^-^ e.coli (NEB C2925H) to avoid methylation of plasmids that were carried over during the lentiviral harvesting.

After 5 days post-transduction, cells were harvested, the DNA extracted using a QiaAmp DNA Micro kit (Qiagen 56304), and processed for DamID as described previously (*72*). DamID fragments were prepared for Illumina sequencing using the NEBNext Ultra II DNA library preparation kit (NEB E7645L). Sequencing was performed as paired end 50 bp reads by the CRUK Genomics Core Sequencing facility on the Illumina NovaSeqX platform. Raw sequencing reads from Dam-only and Dam-Nup153 samples were processed using damidseq_pipeline (v1.5.3) (*73*). Paired-end reads were aligned to the GRCm38/mm10 reference genome using Bowtie2 (v1.3.1), and mapped reads were assigned to GATC fragments (*74*). Reads from Dam-only libraries with lower read-depth were concatenated into a single Dam-only control for each of the 4 Dam-Nup153 replicates. Signal intensities were computed in 300 bp bins using RPM (reads per million) normalization and all resulting binding profiles for one Dam-fusion construct were quantile normalized to each other. The resulting logarithmic profiles in bedgraph format were averaged for all GATC-bins.

Peak calling on the .bam files of each pairwise comparison was performed using MACS3 (v3.0.3) with the Dam-Nup153 sample used as the treatment and the corresponding Dam-only sample used as control (*75*). Peaks were called in broad mode, with a fixed fragment size of 300 bp and no model estimation (--nomodel). The effective genome size was computed from the mm10 reference genome, and significance thresholds were set at p < 0.05 with an mfold range of 5 to 50. Reproducible peaks from the pairwise comparisons were identified across the biological replicates and were only considered if present at a stringent FDR<10^-20^ in at least three of the four replicates using bedtools (v2.31.0) merge and intersect (*76*).

#### Visualizing DAM-ID reads in the Integrated Genomics Viewer (IGV)

To create the bigwig files for visualization in IGV (v2.11.7) (*77*), genomcov with -bg option and scaling in bedtools(v2.31.0) was used to create a bedgraph file (*76*). Following this, the bedgraph file was converted to a bigwig file using the UCSC tool bedGraphToBigWig (*78*).

#### Correlation between replicates

To plot the correlation between each DAM-Nup153 replicate, mutiBigWigSummary function from DeepTools(v3.5.2) was used to calculate average scores for each bigwig files in a binsize of 10000bp across entire genome. These scores were visualized using DeepTools(v3.5.2) function plotCorrelation.

#### Downstream analyses

Reproducible peaks were annotated to different genomic elements such as promoters, enhancers, and intergenic regions, and assigned to the nearest genes using the ChIPSeeker(v1.30.3) package in R (v4.1.2). Pie charts for the distribution of peaks in different genomic regions were generated using the plotAnnoPie function in ChIPSeeker. Gene ontology analysis was performed using the enrichGO package in clusterProfiler (v4.2.2) package within R (*79*). The Benjamin Hochberg multiple testing correction was applied and used with an adjusted p-value cutoff of 0.01. Motif analysis of the reproducible peaks was performed using Homer (*80*) using command findMotifsGenome.pl with motif length of 8,10 &12.

To compare the overall change in the expression of several genesets in Fig. 5G, 5J, log_2_ fold changes of the respective gene sets in the respective conditions were compared to log_2_ fold changes of the remaining genes. Effect size was reported using Cohen’s d, which was calculated using the cohens_d function of the effectSize (v0.8..8) package in R. Fisher’s exact test was performed to evaluate overrepresentation between gene-sets in Fig.5E, 5F, 5H in R (v4.5.1) using function “fisher.test”. Venn diagrams were generated using the online data visualization tool https://www.meta-chart.com/venn.

### RNA isolation and qPCR

PCNs were either processed in the basal state or after drug treatment. Total RNA was extracted with Trizol according to the manufacturer’s instructions (Invitrogen). cDNA preparation was performed with the iScript gDNA Clear cDNA Synthesis Kit (BioRad) according to the manufacturer’s instructions. Quantitative reverse transcription PCR (qPCR) was performed using the BioRad CFX Connect system. The primers used for qPCR are listed in the Table S8. The ΔΔCt method was used to quantify the expression of *Fos, Egr1, Npas4*, and *Nup153* obtained via qPCR. Expression of *GAPDH* was used as an endogenous control. The normality of the dataset was assessed using the Shapiro–Wilk test. An F-test was performed to compare variance in the dataset. The statistical significance of the expression levels (qPCR data) obtained for the activity-dependent genes was assessed using a two-tailed one-sample t-test.

### RNA-Sequencing

#### Sample preparation

Samples confirmed to have sufficient Nup153 knockdown were further processed for RNA sequencing. To remove any contaminating DNA, samples were treated with TURBO DNAase (Life Technologies), and RNA integrity was checked by Bioanalyzer (Agilent). Only samples with RIN number greater than 8 were used for further processing. A total of 3-4 biological replicates were used for each condition analyzed in the study, with one biological replicate being primary neurons derived from pooled cortical tissue from embryos of one litter.

#### Library preparation and sequencing

mRNA was isolated from an average of 150-300 ng total RNA by poly-dT enrichment using the NEBNext Poly(A) mRNA Magnetic Isolation Module (NEB) according to the manufacturer’s instructions. Samples were then directly subjected to the workflow for strand-specific RNA-Seq library preparation (Ultra II Directional RNA Library Prep, NEB). Ligation was performed using the NEBNext Adapter of the NEBNext Multiplex Oligos for Illumina Kit. After ligation, the adapters were depleted by XP bead purification (Beckman Coulter). Unique dual indexing was used during the subsequent PCR enrichment (12 cycles) with amplification primers carrying the same sequence for i7 and i5 indices (primer 1: AAT GAT ACG GCG ACC ACC GAG ATC TAC AC NNNNNNNN ACA TCT TTC CCT ACA CGA CGC TCT TCC GAT CT, Primer 2: CAA GCA GAA GAC GGC ATA CGA GAT NNNNNNNN GTG ACT GGA GTT CAG ACG TGT GCT CTT CCG ATC T). After purification, libraries were quantified on a Fragment Analyzer (Agilent). Libraries were sequenced on an Illumina NovaSeq 6000 in 100 bp paired-end mode to a depth of 30-55 million fragments per library.

### ATAC-Sequencing

#### Sample preparation and tagmentation

ATAC-seq was performed as previously described (*81*). Briefly, cells were treated with DNase (Worthington) for 30 min at 37°C to digest any free-floating DNA and DNA from dead cells. Subsequently, PCNs were directly lysed in 1 ml cold ATAC-Seq resuspension buffer (RSB: 10 mM Tris-HCl pH 7.4, 10mM NaCl, and 3 mM MgCl2 in water, containing 0.1% NP40, 0.1% Tween-20, and 0.01% digitonin). After cell lysis, approximately 50,000 nuclei were centrifuged and washed with 1ml of RSB without detergent. The nuclei were then centrifuged again, the supernatant was removed, and the nuclei were resuspended in 50 µl transposition mix (25 µl 2X Tagmentation DNA buffer (20 mM Tris-HCl pH 7.6, 10 mM MgCl2, 20% Dimethylformamide), 2.5 µl transposase (100 nM final), 16.5 µl PBS, 0.5 µl 1% digitonin, 0.5 µl 10% Tween-20, and 5 µl water). Transposition reactions were incubated for 30 minutes at 37°C with shaking at 1000 rpm. Samples were then purified using the MinElute PCR clean up kit (Qiagen). Two biological replicates per condition were processed.

#### Library preparation and sequencing

A total of 10 µl of purified tagmented DNA was indexed and pre-amplified for the initial 5 PCR cycles with 1x KAPA HiFi HotStart Readymix and 100 nM unique dual-index P5 and P7 primers compatible with Illumina Nextera DNA barcoding, using the following PCR conditions: 72°C for 5 min, 98°C for 30 s, thermocycling for 5 cycles at 98°C for 10 s, 63°C for 30 s and 72°C for 1 min. Subsequently, qPCR was performed on the LightCycler 480 (Roche) with 1 µl of the pre-amplified material to determine the remaining PCR cycle numbers to avoid saturation and potential bias in library amplification. Purification and double-sided size selection of amplified libraries was performed using AMPure XP beads (Beckmann Coulter), and libraries were checked for their quality and quantity on a Fragment Analyzer (Agilent). Libraries were sequenced on the Illumina NovaSeq 6000 in 100 bp paired-end mode to a depth of 70-110 million read pairs.

### RNA sequencing data analysis

#### Alignment and read counts

FastQC was used to perform basic quality control of the sequencing data. Fragments were aligned to the mouse reference (mm10) with support of the Ensembl 98 splice sites using the gsnap aligner (v2020-12-16) (*82*). Fragments per gene and sample were obtained based on the overlap of the uniquely mapped fragments with the same Ensembl gene annotation using featureCounts (v2.0.1) (*83*).

#### Differential expression analysis

Differential gene expression analysis was performed using the DESeq2 (v1.34.0) package within the R programming environment (R v4.1.2) (*84*). For exploratory analysis, sample to sample Euclidean distance, Pearson and Spearman correlation coefficients (r), and PCA were calculated based on the top 500 genes with highest variance to check the correlation between biological replicates and conditions. Genes with a maximum false discovery rate of 1% (padj ≤ 0.01) and log_2_ fold change of < −0.58 and > 0.58 were considered as significantly differentially expressed for the comparisons of shNup153-1 vs. shControl and shNup153-2 vs. shControl. Genes that were significantly dysregulated by both shRNAs in the basal state and also significantly dysregulated in the presence of TTX-AP5 (padj ≤ 0.01) were classified as activity-independent, and that lost significance were classified as activity-dependent (Fig. 2J). Each category was further annotated by shRNA specificity (shNup153-1-specific, shNup153-2-specific) (Fig. S6D, Table S1). Log fold changes and p-values for differential peaks were visualized by creating volcano plots using the EnhancedVolcano (1.12.0) package in R (v4.1.2). Gene Ontology analyses were conducted using the enrichGO package in the clusterProfiler (v4.2.2) package within R (*79*). Specifically, to compare Gene Ontology enriched in Nup153-dysregulated genes, we used the “compareCluster” function to evaluate enriched ontologies (biological processes) within the genes defined in org.Mm.eg.db (the entire mouse genome ∼22000 genes). Upon enrichment, we visualized the top enriched terms to compare the different categories in question.

To generate a heatmap for selected ARGs, normalized count data using the rlog and pheatmap package (version 1.0.12) in R with scaling by row was used to generate a heatmap. Gene set enrichment analysis (GSEA) was performed on a gene set of ARGs defined in a previous work using the GSEA function of the clusterProfiler (v4.2.2) package in R(*27, 85, 86*). Heatmaps for each gene-set (Fig. S6 and S7) were generated using rlog-transformed expression values from DESeq2 the relevant shRNA conditions under the basal and activity-silenced (TTX-AP5) states. Expression values were row-scaled to z-scores (centered and scaled per gene) and plotted with ComplexHeatmap (2.26.0) (*87*), with columns split and ordered by condition and rows ordered by significance category. The per-condition z-scores were also displayed as boxplots using ggplot2 (v4.0.2) in R (v4.5.1) (*79*), faceted by significance category, with the group mean overlaid. In parallel, the underlying DESeq2 rlog values for the tested conditions were plotted as boxplots to compare normalized expression of the gene collection (and of individual genes) across conditions. Differences across conditions were assessed by one-way ANOVA with Benjamini–Hochberg-corrected pairwise t-tests, and effect sizes were quantified using Cohen’s d (rstatix v(0.7.3)).

To compare the overall change in the expression of the previously defined ARGs, log_2_ fold changes of the ARGs in the respective conditions were compared to log_2_ fold changes of the remaining genes. Effect size was reported using Cohen’s d using the cohens_d function of the effectSize (v0.8..8) package in R. Venn diagrams were generated using the online data visualization tool https://www.meta-chart.com/venn.

#### DecoupleR based transcription factor analysis

Transcription factor (TF) activity was inferred from differential expression statistics using decoupleR (v(2.16.0) (*88*). For each comparison, the gene-level Wald statistic from DESeq2 was used as the input signature, and analyses were performed for the shNup153-2 knockdown under basal and activity-silenced (TTX-AP5) conditions. The CollecTRI mouse regulon was used as the prior-knowledge network of signed TF–target interactions (*89*). Analyses were run separately for two predefined gene-sets: activity-independent and activity-dependent activity-regulated genes (ARGs) (Fig. 2J). TF activity was then estimated by applying multiple enrichment statistics (weighted mean, univariate linear model, and VIPER) and aggregated into a single consensus score across methods. To assess robustness to regulon size, the analysis was repeated across minimum-regulon-size thresholds (minsize = 1 and 2), and results were aggregated across runs. Inferred TF activity scores were visualized as a heatmap restricted to TFs reaching statistical significance. A network plot was generated in which each TF was connected to its CollecTRI target genes, retaining only targets present in the ARG set evaluated for that TF. Edges were signed by the CollecTRI mode of regulation, and the log2 fold change of each ARG (under the basal or TTX-AP5-treated conditions) was overlaid to indicate the direction and magnitude of expression change.

### ATAC-Sequencing data analysis

#### Aligning ATAC-seq reads

To align ATAC-seq reads, the nf-core/atacseq pipeline written in the Nextflow domain-specific language was used (*90*). Briefly, the nf-core pipeline for ATAC-seq (v2.0) was used to perform a quality check on fasta files generated after sequencing, followed by alignment of reads to the mm10 genome using the bwa aligner and filtering aligned reads. After the quality check, separate bam files for each replicate, and bam files where filtered reads for replicates are merged. The bam files for merged replicates were used for peak calling and to create bigwig files. The code used to run the nf-core pipeline for ATAC-seq analysis is the following: nextflow run nf-core/atacseq --input Samplesheet.csv --genome mm10 --read_length 200 --outdir Results -profile singularity -c config.txt -r 2.0.

#### Peak Calling

[Genrich](https://github.com/jsh58/Genrich) (v0.6.1) was used to evaluate genomic regions with peaks of significant enrichment. Peak calling was performed in ATAC-Seq mode with a q-value threshold of 0.05, and an area under the curve (AUC) threshold of 2. An example of the code used for peak calling with Genrich is : ./Genrich -t ∼/Input.bam -o Output.narrowPeak -r -j -q 0.05 - e chrM -a 2. Following peak calling, Bedops (v2.4.41) tool commands such as sort-bed and merge were used to create merged peak files for further analysis (*85*).

#### Visualizing ATAC-seq reads in the Integrated Genomics Viewer (IGV)

To create bigwig files for visualization in IGV (v2.11.7) (*77*), genomcov with -bg option and scaling in bedtools(v2.31.0) was used to create a bedgraph file (*76*). Following this, the bedgraph file was converted to a bigwig file using the UCSC tool bedGraphToBigWig (*78*).

#### Differential analysis

Differential analysis was performed for basal and TTX-AP5-treated samples separately. To perform differential analysis, a consensus peak file for control and shNup153-2-treated samples was created. Following this, the count of reads for each peak interval was summarized for the two replicates in control and shNup153-2-treated samples using the Subread/RSubread (v2.8.2) package featureCount (*83*). This was used as the count matrix input in DESeq2(v1.34.0) in R (v4.1.2) for performing differential analysis (*84*). Peak intervals with a maximum of false discovery rate (FDR) of 5% (padj < 0.05) were considered as differentially accessible peaks or DAPs in shNup153-2-treated samples compared to shControl-treated PCNs. Motif analysis of the differential peaks was performed using Homer (*80*).

#### Association between gene expression and chromatin accessibility

To examine the association of differential accessibility with changes in gene expression upon Nup153 depletion, the log_2_ fold change in gene expression and the corresponding log_2_ fold change in accessibility in nearby ATAC peaks were compared on a graph. To visualize the association, we plotted the relationship of log_2_ fold change in gene expression compared to changes in associated-chromatin accessibility. The density of the plotted genes was calculated using 2D kernel density estimation function from the MASS R package.

#### Consensus peak selection

A single consensus ATAC peak set was constructed spanning both the basal and TTX-AP5-treated states. Within each state, the per-condition peak calls (shControl and shNup153-2) were coordinate-sorted with sort-bed and merged with bedops -m to give a basal consensus and a TTX-AP5 consensus; these two state-level consensus sets were then merged together (bedops -m) into a single combined reference peak set (BEDOPS v2.4.41). Read counts over this consensus set were quantified with featureCounts (*83*) (Subread/Rsubread v2.8.2) for the two replicates of each condition, and counts were depth-normalized in DESeq2 by dividing each peak’s raw count by the corresponding sample’s size factor, which was estimated using the median-of-ratios method (counts(dds, normalized = TRUE)).

#### ATAC signals at overlapping and non-overlapping DAPs

Overlapping or non-overlapping DAPs (Fig.4C) were used as reference regions and extended by 5 kb (2.5 kb on each side) around their center. A normalized ATAC signal was assigned to each reference region by overlap with the combined consensus peak set. The reference region which overlapped with more than one consensus peak was aggregated into a single per-region value by averaging count. Regions lacking signal were removed, and per-region values were row-scaled to z-scores. Scaled signals were displayed as a heatmap (ComplexHeatmap v(2.26.0) (*87*), with samples ordered by conditions and states, and rows clustered by k-means (k = 2) to separate regions with increased versus decreased accessibility between conditions. Cluster assignments were then used to summarize signal per cluster: size-factor-normalized counts were log1p-transformed and plotted as boxplots across conditions and states, faceted by cluster, and control versus shNup153-2 differences were tested within each cluster and state using Wilcoxon rank-sum tests (Benjamini–Hochberg-corrected) with Cohen’s d as the effect-size measure.

#### ATAC signal aggregation at DAPs near activity dependent and independent genes

For gene-focused analysis, basal-state DAPs were restricted to a defined RNA significance category (Fig.2J) (activity-dependent or activity-independent) and to peaks annotated near the genes of interest. Region-level signals were derived as above, after which all regions annotated to the same gene symbol were aggregated by averaging to a single value for each gene. Genes lacking signal or with zero variance were removed, per-gene values were row-scaled to z-scores, and the gene-level matrix was visualized and clustered as above. Per-cluster signal was again summarized as log1p-transformed normalized-count boxplots with Wilcoxon tests and Cohen’s d, and ARGs within the selected set were labelled on the heatmap.

Specifically for Figure S8G, all basal-state peaks belonging to a single gene were selected, regardless of whether they were DAPs or did not change significantly in response to shNup153-2 treatment. ATAC and H3K27ac signals were both derived and aggregated across all peaks of that gene as described above: for each modality, the consensus-peak counts overlapping the gene’s peaks were averaged across peaks, then averaged across all regions mapping to that gene symbol to give one ATAC and one H3K27ac value per gene.

#### TOBIAS based occupancy hrelated analysis

Transcription factor (TF) footprinting was performed with TOBIAS (v0.14.0) (*32*). Merged ATAC-seq BAM files for each condition (shControl, shNup153-2, basal, and TTX-AP5) were first corrected for Tn5 insertion bias using the command “ATACorrect”, with bias estimated over the relevant peak set. Two peak sets (“Overlapping DAPs” and “Non-overlapping DAPs”) were processed (Fig. 4C). Bias-corrected signals were converted to continuous footprint scores per condition with the command “FootprintScores”, and differential TF occupancy was then evaluated across conditions with “BINDetect`”` using JASPAR motifs, generating per-motif and per-motif-cluster differential binding scores and p-values for each pairwise contrast. BINDetect output was summarized as ranked tables and dumbbell plots in R (v4.5.1) (Fig.S9, Table S2). For the ranked tables, motifs and motif clusters were ordered by the significance of the basal contrast (shControl vs shNup153, oriented as shControl − shNup153, where negative changes indicate higher binding in the knockdown), and the corresponding TTX-AP5 contrast was carried out . Each entry was assigned a qualitative assessment based on the ratio of the TTX-AP5 to basal state differential binding score: motifs with a negligible basal effect were flagged as flat, sign reversals as “Reversed,” and otherwise classified as “Attenuated” (ratio < 0.66), “Amplified” (ratio > 1.50), or “Maintained.” For the dumbbell plots, the top 25 motifs or clusters by basal significance were displayed with their basal and TTX-AP5 differential binding scores as paired points joined by a connecting segment, visualizing the magnitude and direction of each TF’s occupancy change and how it shifts withTTX-AP5 treatment. Plots were generated with ggplot2 (v4.0.2).

### ChromHMM

To learn the relevant chromatin states underlying basal and active neurons, we trained a multivariate Hidden Markov Model using ChromHMM (*33*). We used previously published ChIP-seq data for transcription factor and histone modifications Fos, Jun, CBP, CREB, Pol2, H3K4Me3, H3K4Me1, H3K27Me3, H3K27Me3(*8*), and H3K27ac (*34*), in basal and active neurons.

The bigWig format files obtained from GSE21161 and GSE60192 were converted to bedGraph format using the UCSC tool bedGraphToBigWig (*78*). The BedGraph files were then converted to BED format using a custom script. As some of the datasets were aligned to the mm9 genome, they were lifted over to the mm10 genome using the UCSC *liftOver* tool (*91*). The ChromHMM option *BinarizeBed* was used to convert the BED files for each mark into a binarized data format, where 1 corresponds to enrichment for the mark in a 200bp bin, and 0 indicates absence. Control reads for the ChIP-seq data were used for reliable binarization whenever available; otherwise, binarization was performed based on a Poisson background model. The ChromHMM command *LearnModel* was used to learn chromatin states in active and basal neurons. We tested models with 10 to 24 states and ultimately selected a 15-state model, as this captured meaningful chromatin states relevant to active and basal neurons. *OverlapEnrichment* was employed to evaluate significant enrichment of different genomic subsets with ChromHMM-learned chromatin states in active or basal neurons.

### ChIP-Sequencing data analysis

#### Aligning ChIP-seq read and visualization

To align ChIP-seq reads, the nf-core/atacseq pipeline was used (v2.0) (*90*). After alignment and filtering of reads, the nf-core pipeline generated separate bigwig files, which were used for visualization of reads in the IGV browser.

#### Peak Calling and analysis

Peak calling was performed within the nf-core pipeline using MACS2 with the default options.The Qvalue cutoff for narrow peaks was 5.00e-02, and the Qvalue cutoff for broad peaks was 0.1. The average fragment length for HDAC1 and H3K27ac was estimated by MACS2 to be in the range of 300-350bp. After peak calling, GenomicRanges was used to obtain genomic regions with peaks reproduced in both replicates (*92*). Reproducible ChIP-seq peaks were annotated with different genomic elements such as promoters, enhancers, and intergenic regions, and were assigned to the nearest genes using the ChIPSeeker. The list of genes found near HDAC1 peaks was overlapped with the previously discovered ARGs and Nup153-regulated genes to obtain a subset of genes associated with HDAC1.

To assess changes in global histone acetylation levels upon Nup153 depletion, DiffBind (v3.12.0) in R was used(*93*). Briefly, DiffBind counted the number of reads in the consensus peak set derived from two replicates of shControl or shNup153-2-treated samples using the dba.count function. The count data for each condition was then normalized by the total library size of that respective sample using the dba.normalize function. Non-specific counts in the peak set were removed by normalizing each sample to its corresponding input control. Finally, we evaluated differential peaks between shControl and shNup153-2-treated samples using DESeq2, which can be specified within the dba.analyze function in the DiffBind package. To evaluate differences between the basal state and the TTX-AP5-treated state, samples were jointly processed by running dba.count, and dba.normalize followed by dba.plotPCA (Fig.6F). To visualize chromatin signal in DAPs near activity dependent and independent genes (Fig. 2J, 6G), the computeMatrix function from DeepTools (v3.5.6) was used in reference-point mode (centered on the region midpoint) to score the indicated bigwig tracks across a ±5000 bp window in 100 bp bins. The resulting matrix was rendered as a profile heatmap using the plotHeatmap function from DeepTools (v3.5.6).

Log fold change and p-values for differential peaks were visualized by creating volcano plots using the EnhancedVolcano (1.12.0) package in R. Differential peaks were annotated to different genomic elements such as promoters, enhancers, and intergenic regions, and assigned to the nearest genes using the ChIPSeeker(v1.30.3) package in R (v4.1.2). Pie charts for the distribution of peaks in different genomic regions were generated using the plotAnnoPie function in ChIPSeeker. Gene ontology for the genes associated with differentially acetylated regions was performed using the enrichGO function of the ClusterProfiler package in R(v4.2.2).

#### Association between gene expression and histone acetylation levels

To check the association of differential acetylation with changes in gene expression upon Nup153 depletion, log_2_ fold changes in gene expression and corresponding log_2_ fold changes in acetylation at nearby consensus H3K27ac peaks were compared on an XY graph. To better visualize the association, we plotted the distribution of log_2_ fold change in gene expression compared to the changes in associated histone acetylation levels. The density of the plotted genes was calculated using 2D kernel density estimation function from the MASS R package.

#### Alluvial plots to combine multi-layer dataset

To trace how genes flow across regulatory layers, alluvial diagrams were generated with ggplot2 (v4.0.2) in R(v4.5.1). Each gene was classified along four categories: RNA-seq gene class, Nup153/HDAC1 binding status, ATAC-seq accessibility, and H3K27ac status. The RNA-seq class was taken from the basal-versus-TTX-AP5 classification (activity-dependent, activity-independent). Binding status was assigned by intersecting each gene with the Nup153- and HDAC1-bound gene sets (peak-annotated nearest-gene symbols), giving four categories: Nup153 + HDAC1, Nup153 only, HDAC1 only, or neither. The ATAC and H3K27ac axes each described the state in which a gene’s associated peak was differentially accessible /acetylated in basal only, TTX-AP5 only, both, or neither according to the basal- and TTX-AP5-treated-state differential-peak annotations. Genes were tabulated by their combination of categories across the four axes, and the resulting frequencies were plotted as strata connected by sigmoid alluvia, and colored by binding status, with each stratum labelled by its gene count. Separate diagrams were produced for each RNA-seq class and, in parallel, for the activity-regulated gene (ARG) subset of each class. The underlying classifications were exported as supplementary tables: a per-gene detail table (each gene with its RNA-seq class, binding, ATAC, and H3K27ac calls), a full flow summary counting genes per unique four-axis combination, pairwise transition tables between adjacent layers (RNA-seq → binding, binding → ATAC, ATAC → H3K27ac), and a path-level table listing the member genes of each combination. All tables were generated for both the complete gene sets and the ARG-only subsets (Table S6).

### Multielectrode array (MEA) recording and analysis

MEA recording was done as described previously (*41, 94*). Briefly, MEA plates were sterilized under laminar flow UV-light for 1 hour. Sterilized MEA plates were coated with 100 ug/ml poly-DL-ornithine (PDLO, Sigma) and incubated overnight at 37 °C in a CO2 incubator. The next day, MEA plates were washed 4 times with sterile double-distilled water (DDW) and left to dry. Primary cortical neurons of 150,000 cells were seeded on the active recording area of each well on the MEA plate and incubated at 37 °C with 5% CO2 and 95% humidity. Three days after cell plating, the medium was replenished with BrainPhys-based neuronal media. At DIV8, AAVs were applied. All reagents were obtained from Life Technologies unless indicated otherwise.

Extracellular recordings were performed using the Axion Maestro Edge acquisition system (Axion Biosystems). CytoView MEA plates composed of 24 wells were used. We collected 10 minutes of spontaneous firing activity at a 12.5 kHz sampling rate. Spike detection was performed with the adaptive threshold crossing algorithm with a threshold of 6x standard deviation using AxIS software (Axion Biosystems). All spike trains were exported from AxIS software and were analyzed using custom-written Python scripts as previously described (*94*). The first-order statistics of network-wide mean activity parameters (MBR, Network burst, spike frequency) and topological maps of network activity were computed as previously described (*94*).

### MAGMA

To perform the MAGMA based gene-set analysis (*42*), we followed the following steps. The first step was the annotation step, in which SNPs from the European 1000 Genomes phase 3 cohort were mapped onto genes in the GRCh37/hg19 human genome build with an annotation window extending 2kb upstream and downstream of each gene. The second step involves calculating gene p-values, which determine the association of each gene with the relevant phenotype. We used the 1000 Genomes dataset from the European ancestry as a reference for linkage disequilibrium. SNPs associated p-values were derived from existing meta-analysis for the relevant phenotype.

#### MAGMA summary statistics

We focused on neuropsychiatric disorders and behavior traits including obsessive compulsive disorder (OCD) (*95*), bipolar disorder (*96*), Attention-deficit/hyperactivity disorder (ADHD) (*97*), neuroticism (*98*) schizophrenia (*99*), depression (*100*), panic (*101*), antisocial behavior(*102*), intelligence (*103*), insomnia (*104*), metabolism (*105*) and Alzheimer (*106*). Most of the summary statistics were derived from the following databases: https://cncr.nl/research/summary_statistics/

#### GSEA analysis for disease related genes

To identify genes associated with different diseases, we used p-values obtained in the second step of MAGMA-based analysis. We sorted the genes according to their -log10 p-values, and selected those that passed the false discovery rate threshold of either 1% or 10%. These disease-associated gene-sets were converted to mouse orthologue genes and tested them for gene set enrichment analysis (GSEA) using the GSEA function of the clusterProfiler (v4.2.2) package in R(*27, 85, 86*).

### Quantification and statistical analysis

Statistical analyses were performed using GraphPad prism 8 (GraphPad software Inc.) or in the R programming environment (R v4.1.2) (R core team, 2022). Details of the statistical tests are given in the description of the results.

## Acknowledgments

We are grateful to Dr. F.H. Gage and Dr. H. Bading for their advice. We are thankful to Dr. M. Capelson for sharing her ChIP protocol and Dr. M. Hetzer for sharing Nup153 antibody. Drs. I. Solovei and U. Simon for discussion. We would also like to thank the DRESDEN-concept Genome Center (DcGC) at TU Dresden and the CRUK Cambridge Institute Genomics Core Facility for processing NGS libraries and carrying out sequencing. We are grateful for Z. Blalock, Dr. S. Zocher, Dr. M. Albert, and the members of the Toda lab for their comments on the manuscript.

## Author contributions

Conceptualization, T.T. and A.S.; Consolidation of the idea, A.S. and T.T.; Methodology, A.S. and T.T.; Investigation and Formal Analysis, A.S. M.L.S., K.S. with support from N.R., J.H., A.K., J.H.P, T.T., in biological experiments and A.P., M.L., K.S., E.S, D.E., in bioinformatic analysis; Investigation and Data Analysis (MEA), H.A. and D.K.; Investigation and Data Analysis (DAM-ID), S.P., A.S. and J.V.D.A.; Resources, C.R., and T.T.; Visualization, A.S. and T.T.; Writing – Original Draft, T.T and A.S.; Writing – Review & Editing, All authors; Supervision, T.T.; Project Administration, T.T.; Funding Acquisition, T.T., J.V.D.A., H.A., & A.P.

## Data and materials availability

Next generation sequencing data have been deposited at GEO and are publicly available as of the date of publication. The GEO accession numbers for the RNA-seq, ATAC-seq, DAM-ID seq, and ChIP-seq data reported in this paper are: GSE275919, GSE275916, GSE275917, and GSE300192. The DOI is listed in the table S10. Microscopy data reported in this paper will be shared by the lead contact upon request. The present study did not generate original codes. Any additional information required to reanalyze the data reported in this paper is available from the lead contact upon request. This study did not generate unique reagents. Further information and requests for resources should be directed to and will be fulfilled by the lead contact, Tomohisa Toda.

## Supplementary Materials

Supplementary information is present in Document S1. Fig. S1-S14 and Tables S1-S8.

## Supplementary Tables

**Table S1.** RNA-seq analysis related datasets. Associated with Fig. 2, Fig. S4-S6

**Table S2.** ATAC-seq analysis related datasets. Associated with Fig.4, Fig. S7-S9

**Table S3.** Nup153 DAM-ID analysis related datasets. Associated with Fig.5,6 and Fig.S10

**Table S4.** HDAC1 CHIP-Seq analysis related datasets. Associated with Fig.6 and Fig.S11

**Table S5**. H3K27ac CHIP-Seq analysis related datasets. Associated with Fig.6 and Fig.S11

**Table S6**. Alluvial plots in Fig. 6H and S13A,13B related details.

**Table S7**. MAGMA related dataset in Fig. 7G & 7H.

**Table S8**. Reagent and Resource information.

