## Supplementary Figures for "Nup153 regulates neuronal responsiveness through HDAC1-mediated epigenetic modulation"

Abhinav Soni et al.

**This PDF file includes:**

Figs. S1 to S14

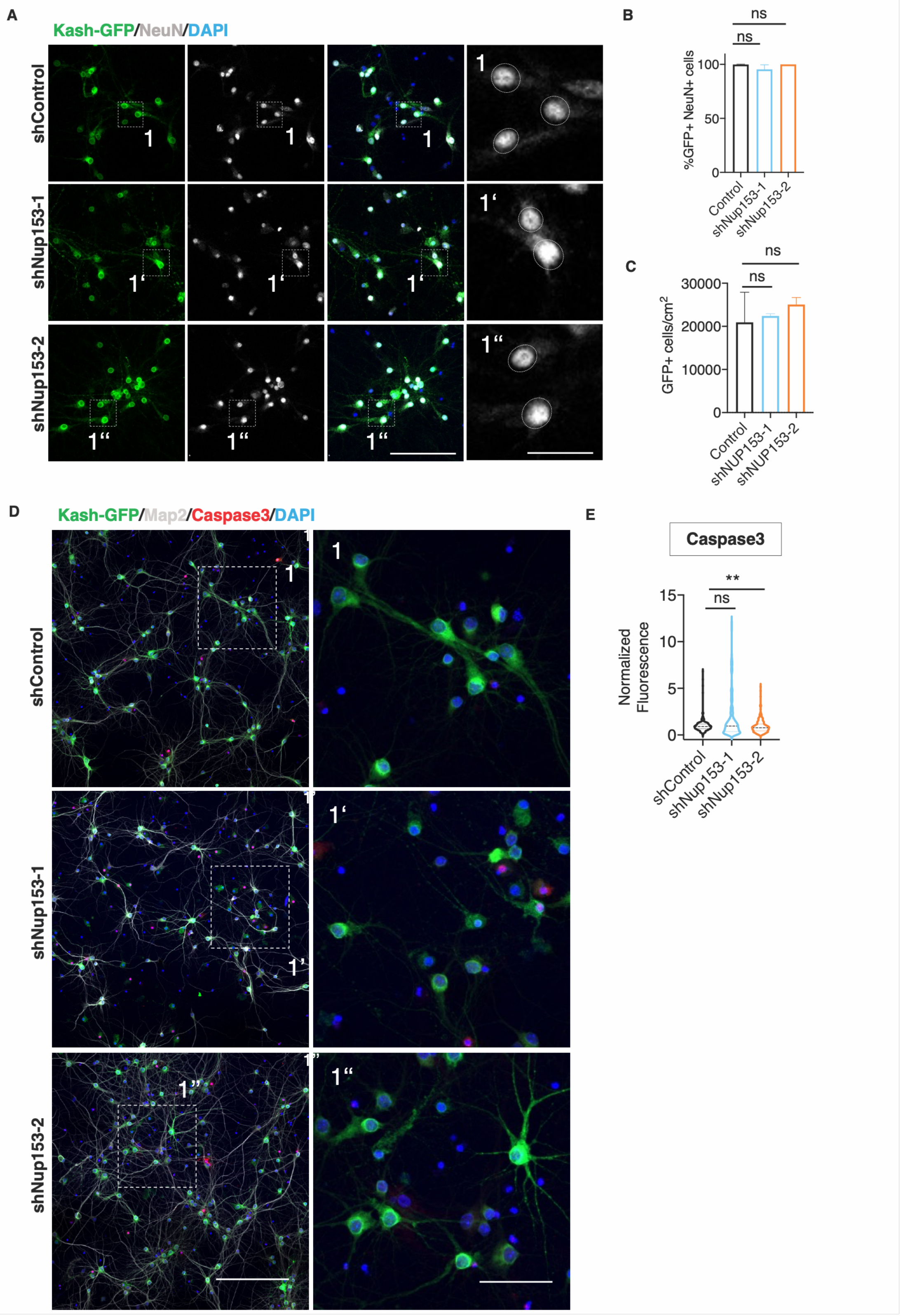

A
Kash-GFP/NeuN/DAPI
shControl
shNup153-1
shNup153-2
B
%GFP+ NeuN+ cells
Control
shNup153-1
shNup153-2
ns
C
GFP+ cells/cm$^2$
Control
shNUP153-1
shNUP153-2
ns
D
Kash-GFP/Map2/Caspase3/DAPI
shControl
shNup153-1
shNup153-2
E
Caspase3
Normalized Fluorescence
shControl
shNup153-1
shNup153-2
ns
**

**Supplementary data associated with the main Fig.1**

**Fig. S1. Effect of Nup153 depletion on neuronal identity and survival**

(A) Confocal images of PCNs after Nup153 knockdown at DIV13, co-stained for KASH-GFP (green) and NeuN (gray). Scale bar = 100 μm.
(B) Quantification of the fraction of $GFP^+$ $NeuN^+$ cells among $NeuN^+$ in (A) ($p > 0.05$, Mann-Whitney test for $GFP^+/NeuN^+$ cell density; Welch's t-test for $GFP^+$ cell density). Three independent experiments were conducted.
(C) Quantification of $GFP^+$ cell density in (A) ($p > 0.05$, Welch's t-test for $GFP^+$ cell density). Three independent experiments were conducted.
(D) Confocal images for active-Caspase3 (red), KASH-GFP (green), MAP2 (gray), and DAPI (blue). Insets 1, 1', and 1" are magnified in the right panels. Scale bars = 350 μm (left) and 25 μm (right).
(E) Quantification of apoptotic cells after Nup153 knockdown. The normalized fluorescence intensity of active-Caspase3 in KASH-$GFP^+/DAPI^+$ PCNs was measured (**$p < 0.01$, Mann-Whitney test). At least three independent experiments were conducted.

The bar graphs in Fig. S1B& S1C are presented as mean ± standard deviation (s.d). Not significant, n.s. The data in Fig. S1E are presented as violin plot showing median line.

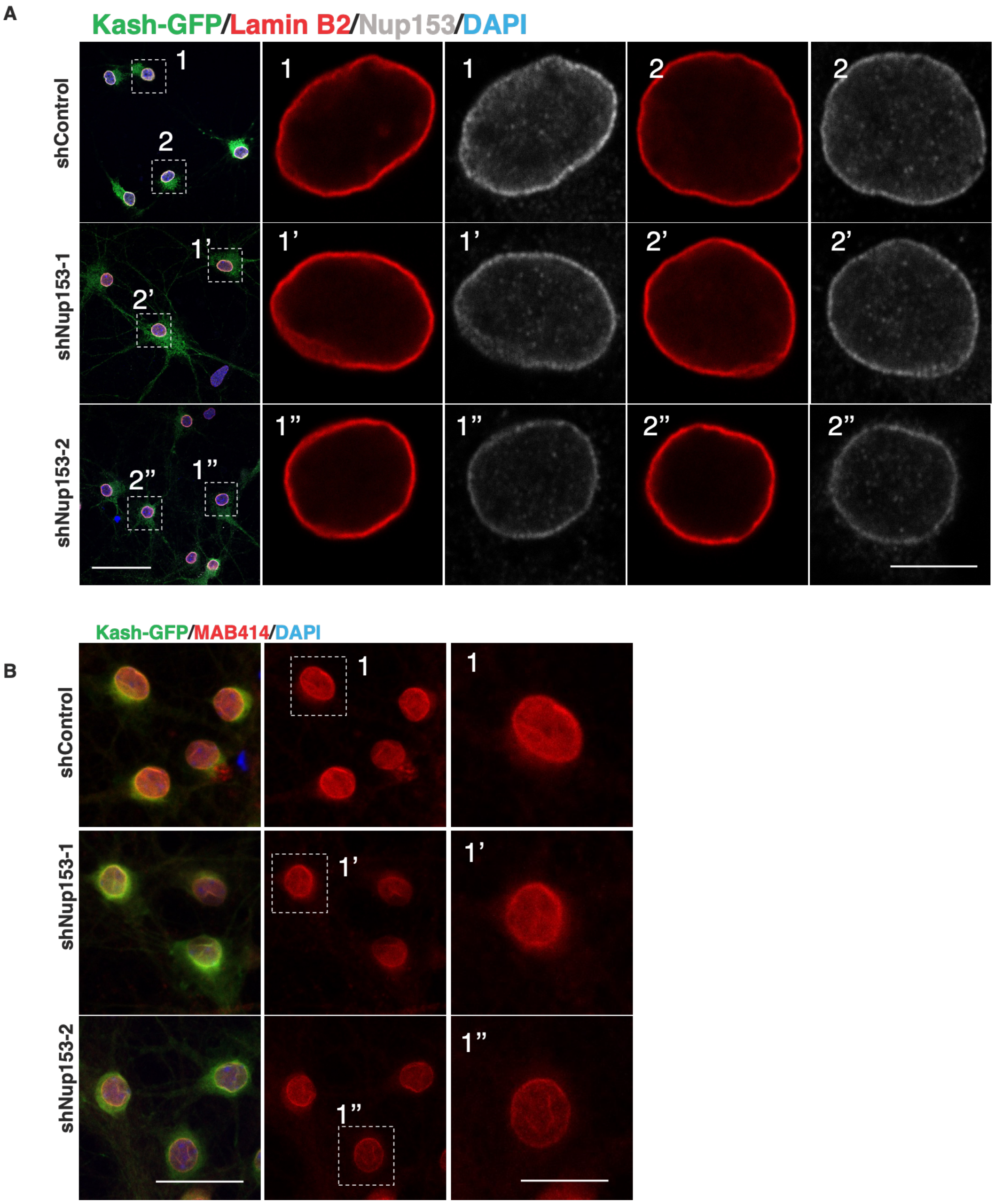


**Supplementary data associated with the main Fig. 1**

**Fig. S2. Effect of Nup153 depletion on nuclear structure and integrity**

(A) Assessment of general nuclear architecture after Nup153 knockdown. Confocal images of Lamin B2 (red), KASH-GFP (green), Nup153 (gray), and DAPI (blue) in PCNs were taken. The insets are magnified in the right panels. Scale bars = 50 μm (left) and 6.25 μm (right).

(B) Assessment of nuclear pores after Nup153 knockdown. Maximum-projected confocal images of Mab414 (red), KASH-GFP (green), and DAPI (blue) in PCNs. Insets within single z-plane images were magnified and are displayed in the right panels. Scale bars = 50 μm (left) and 25 μm (right).

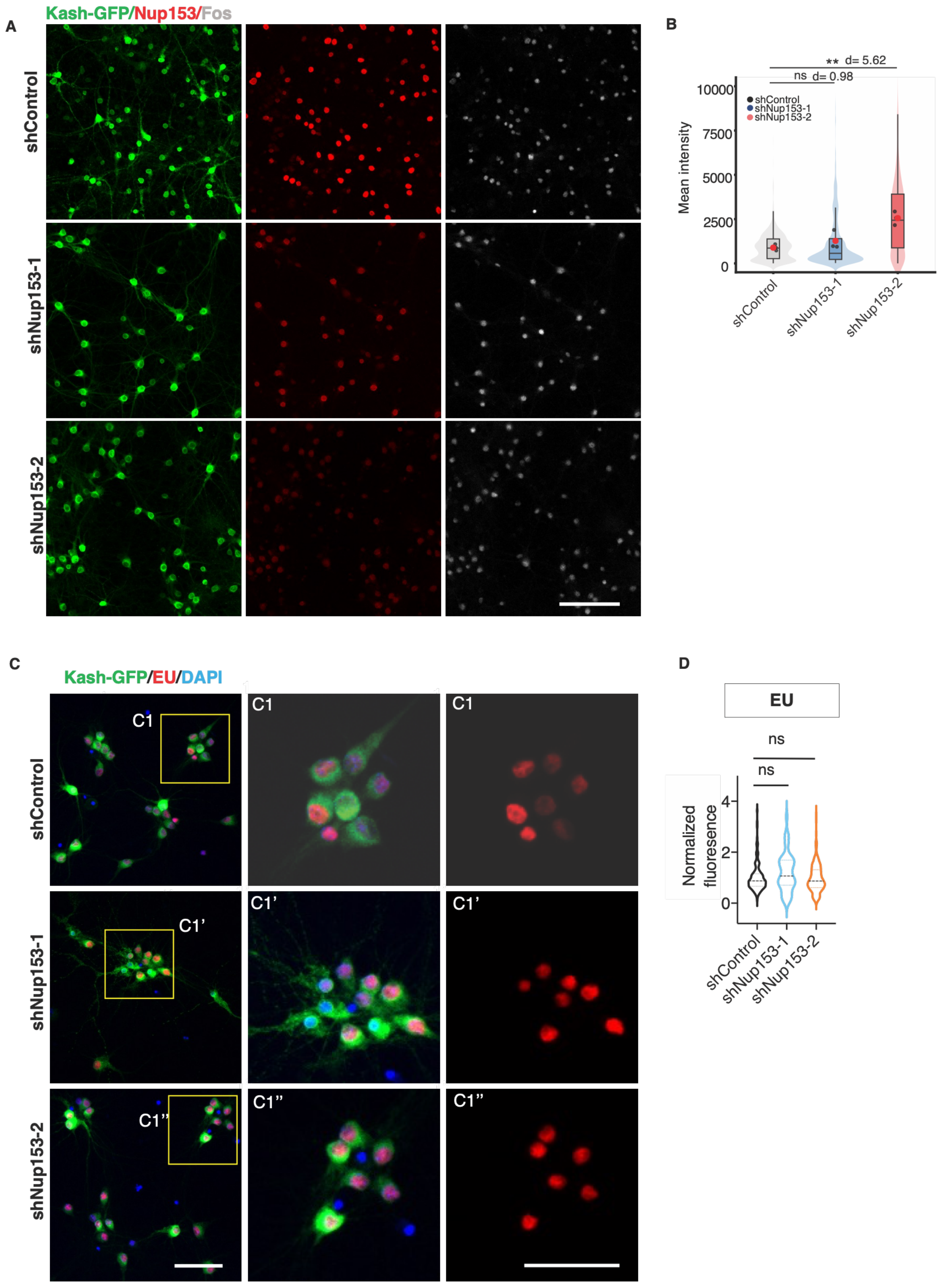


**Supplementary data associated with the main Fig. 1**

**Fig. S3. The essential role of Nup153 in balancing the levels of ARGs in basal neurons**

(A) Changes in Fos expression after Nup153 depletion. Confocal images of Fos (gray), co-KASH-GFP (green), and Nup153 (red) were taken. Scale bar = 100 µm.
(B) Quantification of Fos (grey) levels of KASH-GFP$^+$ (green) neurons in (A). The integrated fluorescence intensity of Fos in KASH-GFP$^+$ PCNs was quantified (**p < 0.001,ns = not significant; shControl, 889 ± 169; shNup153-1, 1273 ± 530; shNup153-2, 2556 ± 385; Welch's two sample t-test; n = 3 biological replicate, Cohen's d=5.62, shControl vs shNup13-2, cohen's d=0.98, shControl vs shNup153-1).
(C) Assessment of nascent transcriptional levels after Nup153 knockdown. Nascent transcribing RNA were labeled with 5-ethynyluridine (EU), co-stained for KASH-GFP (green) and DAPI (blue). Insets 1, 1', or 1" are magnified on the right. Scale bars = 50 µm.
(D) Grap depicts Quantification for data in (C). No changes in EU levels were observed (p > 0.05, Mann-Whitney test). Data from three independent experiments were used.

Not significant, n.s. The data in Fig. S3D is presented as violin plot showing median line.

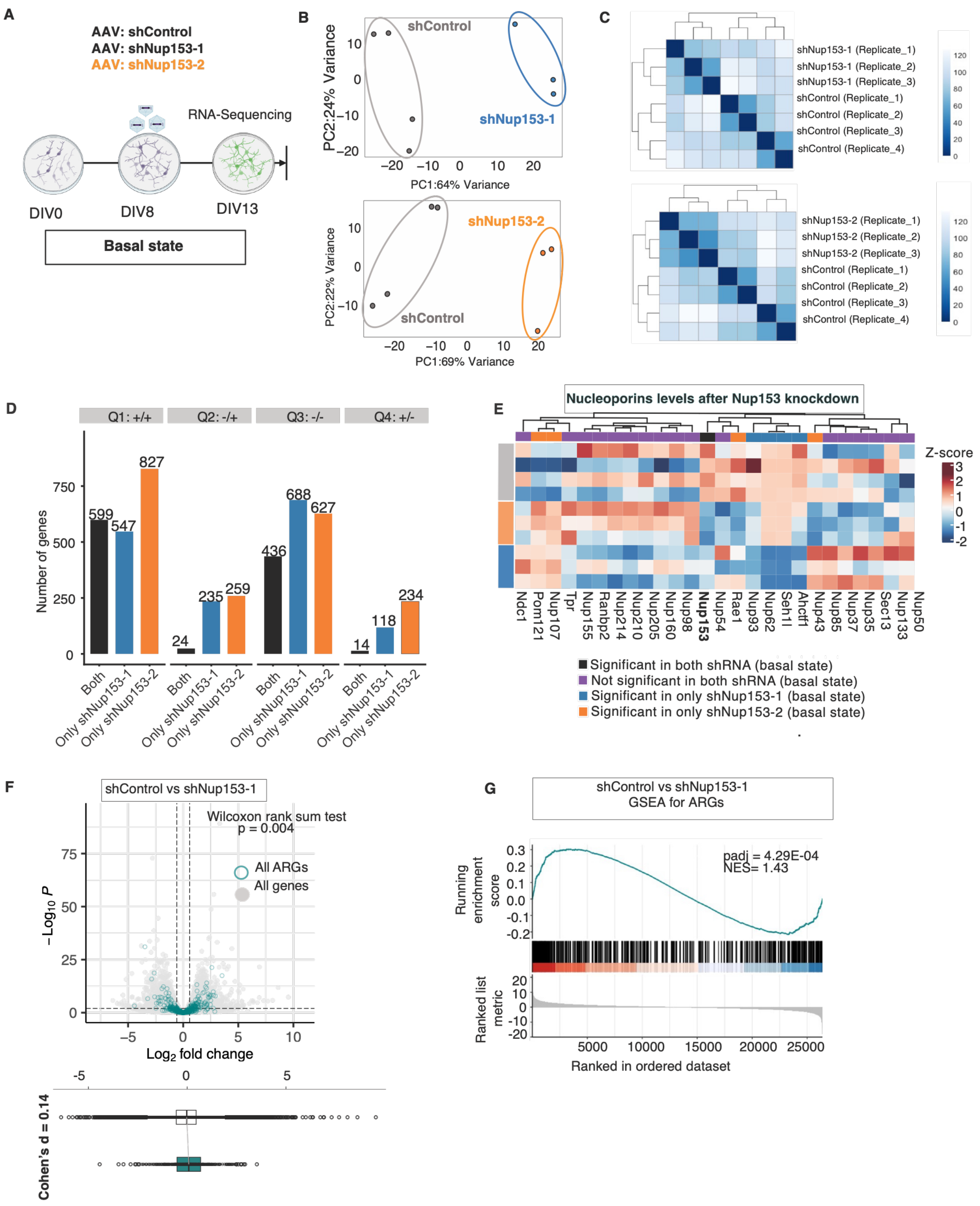


**Supplementary data associated with the main Fig. 2 & Table S1**

**Fig. S4. Global de-repression of ARGs in Nup153-knockdown neurons at the basal state**

(A) A schematic illustration of the experimental plan.
(B) Principal Component Analysis (PCA) for the top 500 genes for shControl vs. shNup153-1 and shControl vs. shNup153-2.
(C) Correlation between each replicate based on the Euclidean distance.
(D) Quantification of the number of DEGs in different quadrants related to main Fig. 2C. The number of genes is counted according to their significance category, i.e., whether they are significantly dysregulated by "both" shRNAs ,or significantly dysregulated using only shNup153-1 (blue) or shNup153-2 (orange).
(E) A heatmap displays regularized log (rlog) values of nucleoporin expression in basal state samples treated with shControl, shNup153-1, or shNup153-2. The scale indicates z-score for changes in gene expression. Note that *Nup153* is the only commonly dysregulated nucleoporin gene.
(F) A volcano plot highlighting the differential expression of ARGs (*8*) (green) upon Nup153 knockdown with shNup153-1. A boxplot shows the fold change in ARGs in comparison to all other genes upon Nup153 knockdown ($p = 0.004$, Wilcoxon Rank sum test, Cohen's $d = 0.14$, 95% CI [0.05, 0.24]).
(G) Gene Set Enrichment Analysis (GSEA) for the previously defined ARGs (*8*) after Nup153 knockdown with shNup153-1.

Not significant, n.s.

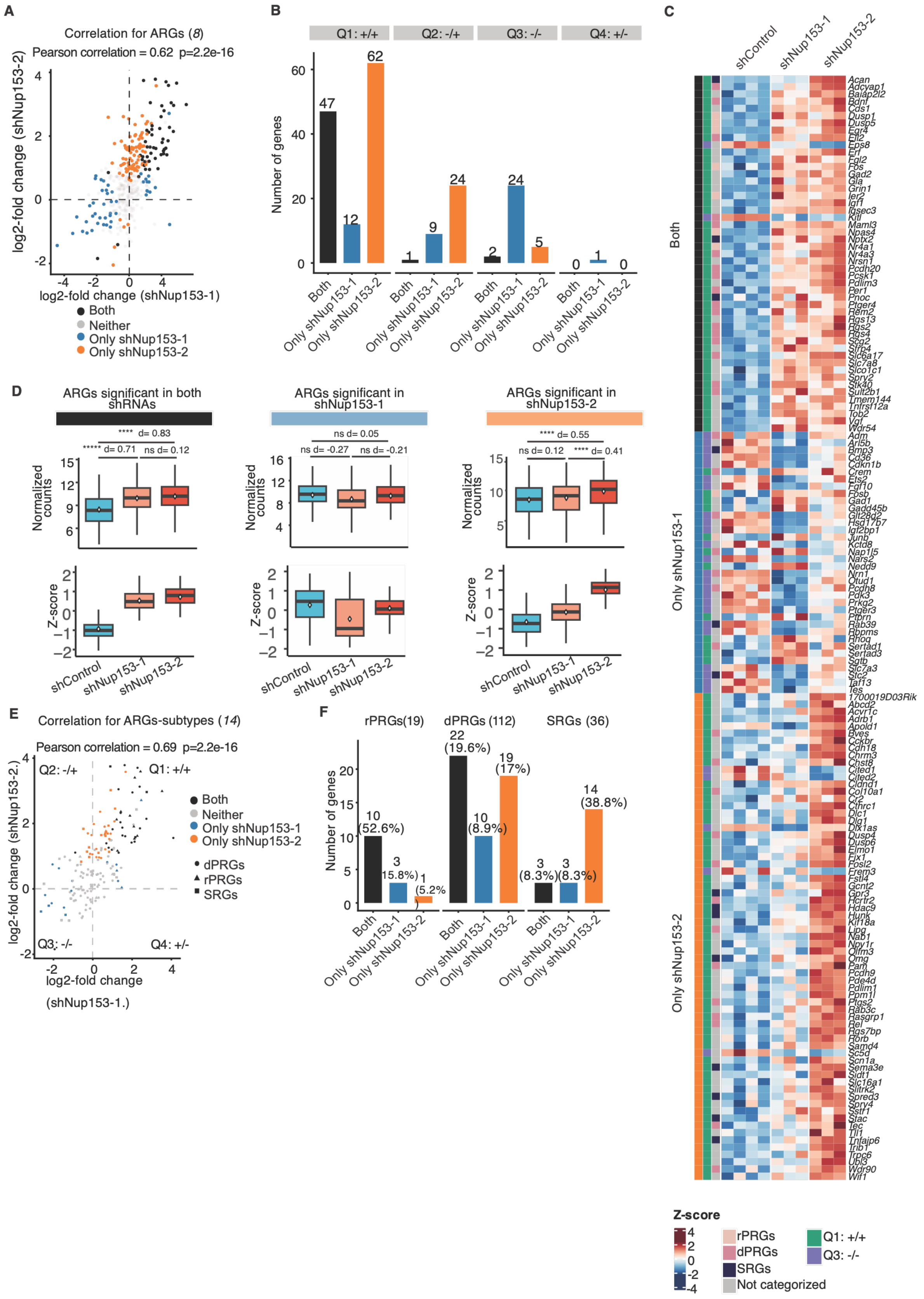


**Supplementary data associated with the main Fig. 2 & Table S1**

**Fig. S5. Global de-repression of ARGs in Nup153-knockdown neurons at the basal state**

A. Correlation of ARGs (*8*) between the two shRNAs. ARGs induced by the two shRNAs were highly similar (r = 0.62, p = 2.2e-16, Pearson's correlation). Common DEG-ARGs between the two shRNAs are shown as black dots.
B. Quantification of the number of DEG-ARGs in different quadrants in C. The number of ARGs are counted according to their significance category.
C. A heatmap displays regularized log (rlog) values of ARGs in Q1+/+ and Q3-/- in (C). The scale indicates the z-score of the respective ARGs between different shRNA-treated conditions. The scale indicates rlog fold changes in gene expression. The heatmap is annotated with three different metacolumns, the first metacolumn indicating quadrants (Q1+/+ (green) or Q3-/- (purple)), the second metacolumn indicating significance category ("Both" (black), "Only shNup153-1" (blue), "Only shNup153-2" (orange)), and the third metacolumn indicating the type of ARGs in accordance with Tyssowski et al. 2018 ( i.e., rapid primary responsive genes (rPRGs) (light pink), delayed primary responsive genes (dPRGs) (dark pink), secondary responsive genes (SRGs) (dark blue), and ARGs not categorized into a subtype of ARGs ("Not categorized") (grey).
D. Box plots showing the mean values of ARGs in (E) categorized by "significance category", for samples treated with shControl/shNup153-1/shNup153-2. Two forms of values are presented: (top) rlog values for each gene per condition and significance category; (bottom) mean z-score values derived from the heatmap in (E) for ARGs (ns=not significant, ****p < 0.0001, pairwise t-test, d = cohen's effect size for the compared conditions).
E. Correlation of the fold changes of ARGs between the two shRNAs within ARGs (*14*) subcategories (r = 0.69, p = 2.2e-16, Pearson's correlation). The black dots indicate ARGs that are significantly dysregulated by both shRNAs. Blue or orange dots indicate ARGs that are significantly dysregulated by one of shRNAs. ARGs in quadrant 1 and 3 (Q1 and Q3) are dysregulate in the same direction by two shRNAs.
F. Nup153 regulated subcategories of ARGs (*14*): rapid primary responsive genes (rPRGs), delayed primary responsive genes (dPRGs) and secondary responsive genes (SRG). All categories indicate significant change using both shNup153-1 and shNup153-2, but rPRGs are affected more preferentially.

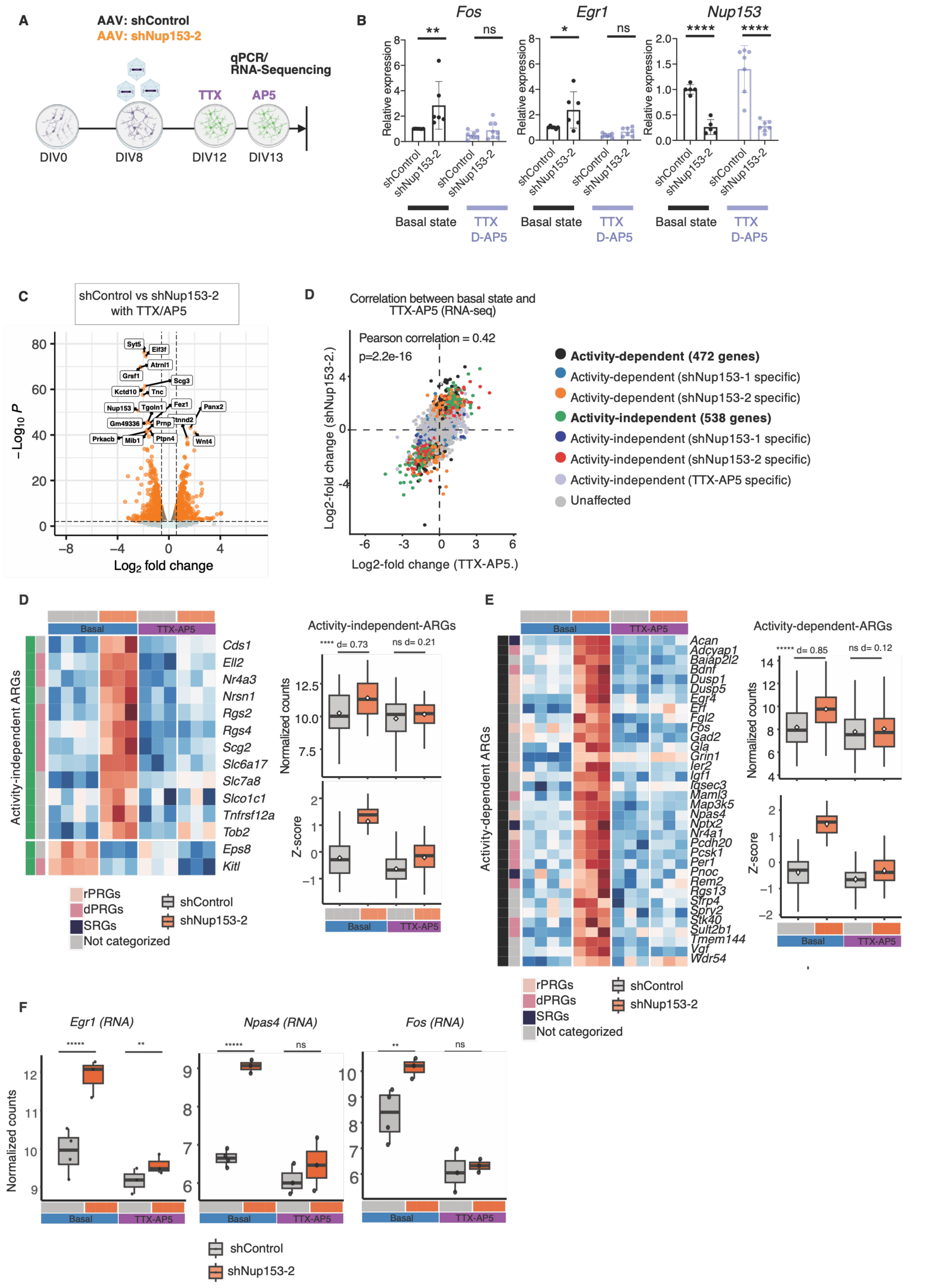
A
AAV: shControl
AAV: shNup153-2
qPCR/
RNA-Sequencing
TTX
AP5
DIV0
DIV8
DIV12
DIV13
B
Fos
Egr1
Nup153
Relative expression
shControl
shNup153-2
Basal state
TTX
D-AP5
C
shControl vs shNup153-2
with TTX/AP5
−Log10 P
Log2 fold change
Syt5
Elf3f
Atrnl1
Grsf1
Scg3
Kctd10
Tnc
Fez1
Nup153
Tgoln1
Panx2
Gm49336
Prnp
Ctnnd2
Prkacb
Mib1
Ptpn4
Wnt4
D
Correlation between basal state and
TTX-AP5 (RNA-seq)
Pearson correlation = 0.42
p=2.2e-16
Log2-fold change (shNup153-2.)
Log2-fold change (TTX-AP5.)
Activity-dependent (472 genes)
Activity-dependent (shNup153-1 specific)
Activity-dependent (shNup153-2 specific)
Activity-independent (538 genes)
Activity-independent (shNup153-1 specific)
Activity-independent (shNup153-2 specific)
Activity-independent (TTX-AP5 specific)
Unaffected
D
Basal
TTX-AP5
Activity-independent ARGs
Cds1
Ell2
Nr4a3
Nrsn1
Rgs2
Rgs4
Scg2
Slc6a17
Slc7a8
Slco1c1
Tnfrsf12a
Tob2
Eps8
Kitl
Activity-independent-ARGs
**** d= 0.73
ns d= 0.21
Normalized counts
Z-score
rPRGs
dPRGs
SRGs
Not categorized
shControl
shNup153-2
E
Activity-dependent ARGs
Acan
Adcyap1
Baiap2l2
Bdnf
Dusp1
Dusp5
Egr4
Ell
Fgl2
Fos
Gad2
Gla
Grin1
Ier2
Igf1
Iqsec3
Maml3
Map3k5
Npas4
Nptx2
Nr4a1
Pcdh20
Pcsk1
Per1
Pnoc
Rem2
Rgs13
Sfrp4
Spry2
Stk40
Sult2b1
Tmem144
Vgf
Wdr54
Activity-dependent-ARGs
***** d= 0.85
ns d= 0.12
Normalized counts
Z-score
Basal
TTX-AP5
rPRGs
dPRGs
SRGs
Not categorized
shControl
shNup153-2
F
Egr1 (RNA)
Npas4 (RNA)
Fos (RNA)
Normalized counts
Basal
TTX-AP5
shControl
shNup153-2

**Supplementary data associated with the Fig. 2G-2M & Table S1**
**Fig. S6. Basal activity-independent vs basal activity dependent Nup153-directed gene regulation.**

(A) A schematic illustration of the experimental plans for TTX-AP5 treatment in PCN.
(B) Quantification of *Fos, Egr1,* and *Nup153* transcript levels in the basal state or after treatment with TTX-AP5. TTX/AP-5 treatment reversed Nup153 depletion–induced upregulation of *Fos* and *Egr1* (*p < 0.05, ** p < 0.01, ****p < 0.0001, *t*-test or Mann-Whitney test, 4 to 7 independent experiments were conducted) (qPCR mean expression *Fos in basal state*: shControl 0.52±0.31 vs shNup153-2 0.90±0.67; qPCR mean expression *Egr1 in basal state*: 0.52±0.31 vs shNup153-2 0.90±0.67).
(C) Correlation between DEGs in basal neurons and those in TTX-AP5 treated neurons (r = 0.42, p = 2.2e-16, Pearson's correlation). We have categorized genes further into activity-dependent or -independent, or shRNA specific activity-dependent or -independent. See methods for more details
(D) (Left) A heatmap displaying the regularized log (rlog) values of "basal activity-independent" DEG-ARGs in Q1+/+ in samples that were treated with shControl or shNup153-2 in the basal or TTX-AP5 treated conditions. The scale indicates the z-score of the respective DEG-ARGs between the different shRNA-treated conditions. The scale indicates rlog fold changes in gene expression. The heatmap is annotated with two different metacolumns. The first metacolumn indicates the activity dependence. The second meta column indicates the type of ARGs in accordance with Tyssowski et al. 2018 (*14*); i.e., rapid primary responsive genes (rPRGs, light pink); delayed primary responsive genes (dPRGs, dark pink); secondary responsive genes (SRGs, dark blue); and ARGs not categorized into a subtype ("Not categorized", grey). (Right) A heatmap associated boxplot quantifying rlog values (top) or z-score (bottom) corresponding to the basal activity-independent-ARGs in Nup153 depleted neurons in basal and TTX-AP5 treated neurons.
(E) Heatmap and boxplot for "basal activity-dependent" DEG-ARGs.
(F) Boxplot quantifying rlog values (top) corresponding to genes *Egr1, Npas4 and Fos* in Nup153 depleted neurons in basal and TTX-AP5 treated neurons (****padj < 0.0001,**padj < 0.01, DESeq2 based adjusted p-value for the compared conditions).

The bar graphs in Fig. S6 are presented as mean ± standard deviation (s.d).

Not significant, n.s.

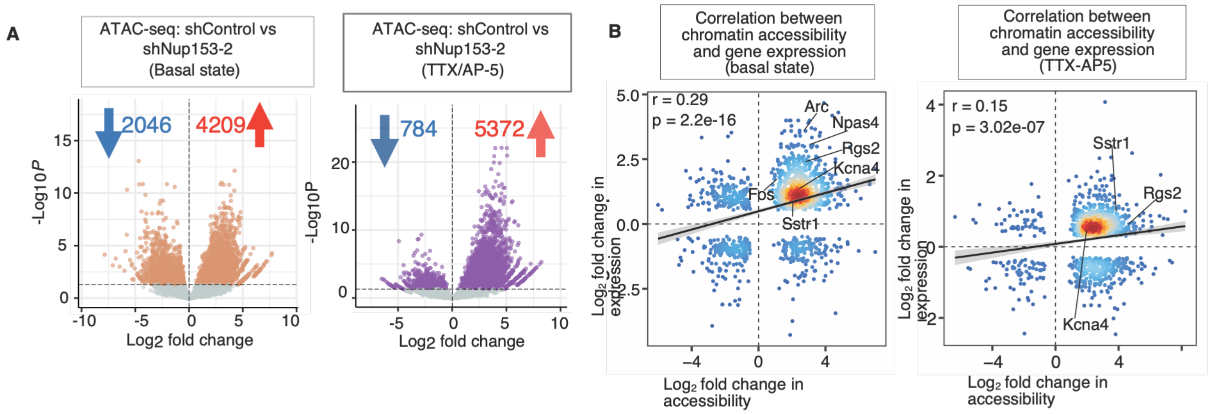


**Supplementary data associated with the main Fig. 4 & Table S2**

**Fig. S7. Nup153-directed, basal-activity-dependent and independent regulation of chromatin organization**

(A) A volcano plot displays changes in chromatin accessibility between shControl- and shNup153-2-treated neurons in the basal state (left) and TTX-AP5 (right). DAPs (marked brown and purple) are defined by a 5% FDR cutoff. Refer also to Table S2.

(B) Association between changes in chromatin accessibility with fold changes in gene expression of the nearest annotated genes in basal state (right) and TTX/AP5-treated Nup153-depleted neurons(left). Genes with increased chromatin accessibility in nearby genomic regions were strongly associated with upregulation in gene expression (r = 0.29, p = 2.2e-16, pearson correlation, basal state; r = 0.15, p = 3.021e-07, pearson correlation, TTX-AP5).

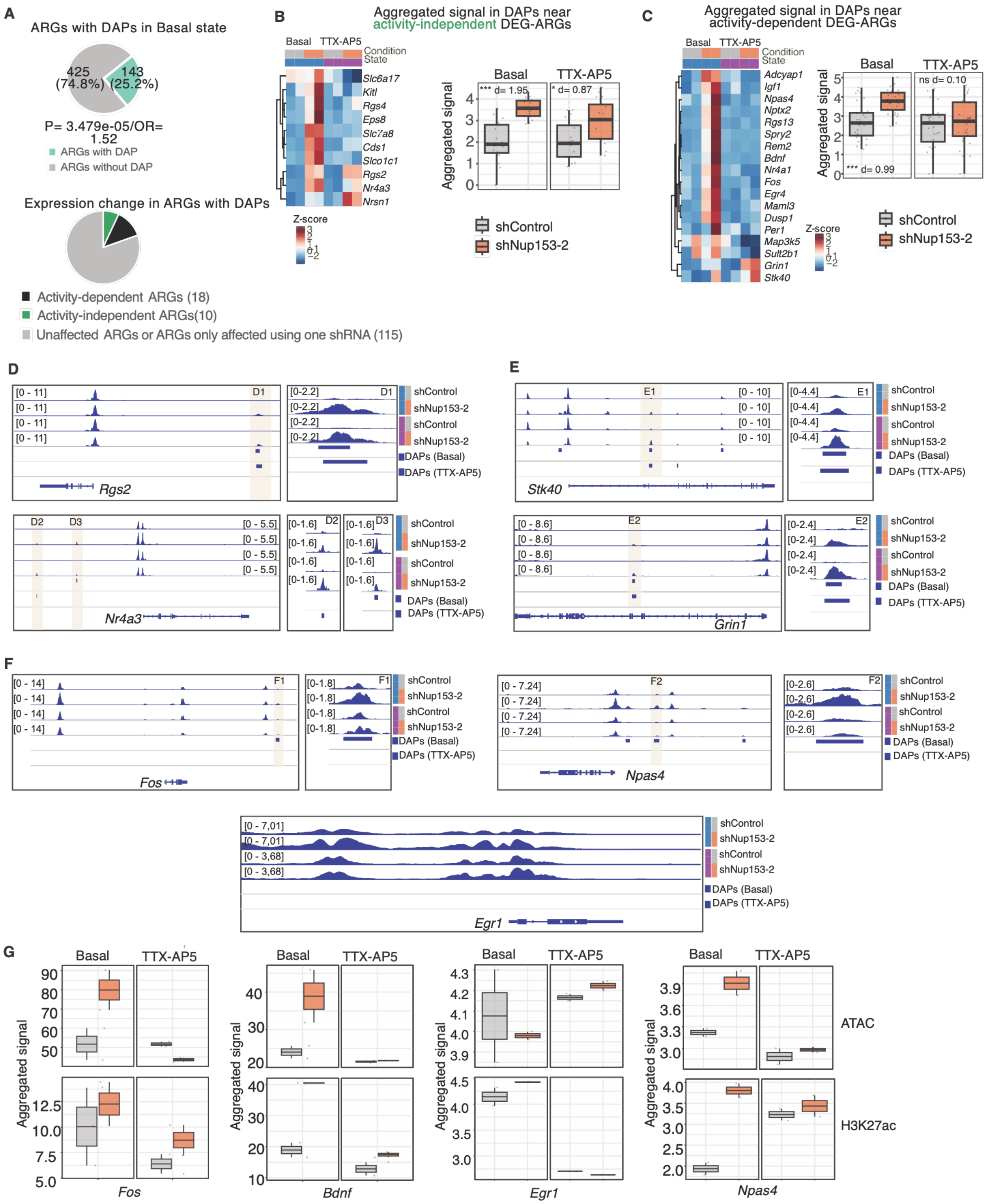


**Supplementary data associated with the main Fig. 4 & Table S2**

**Fig. S8. Nup153-directed, basal-activity-dependent and independent regulation of chromatin organization**

A. (Top) A pie chart summarizing total ARGs (Kim et al.,)(*8*)associated with DAPs. (Bottom) A pie chart visualizing the fraction of DAP-associated ARGs (from Top) with the change of their expression in a basal activity-dependent (green) or activity-independent (black) manner.

B. (Left) A heatmap showing the normalized count values within DAPs annotated near activity-independent DEG-ARGs. Count values are compared between the basal or TTX-AP5 treated conditions. (Right) Normalized aggregated count values for the heatmap are visualized as boxplot (***p < 0.0001, *p < 0.05, Wilcox-test; Cohen's d shControl vs shNup153-2 = 1.95 (Basal state) and 0.87 (TTX-AP5)).

C. (Left) A heatmap showing the normalized count values within DAPs annotated near activity-dependent DEG-ARGs. Count values are compared between the basal or TTX-AP5 treated conditions. (Right) Normalized aggregated count values are visualized as boxplots. (***p < 0.0001,ns = not significant, Wilcox-test; Cohen's d shControl vs shNup153-2 = 0.99 (Basal) and 0.10 (TTX-AP5)).

D. Visualization of ATAC-sequencing peaks around the activity-independent DEG-ARGs *Rgs2* (top) and *Nr4a3* (bottom) treated with shControl- (grey) and shNup153-2- (orange) in basal state (blue) and TTX-AP5 (purple) conditions via the IGV genome browser. DAPs in basal state and TTX-AP5 states are highlighted as dark-blue bars. Further region D1 (Rgs2) and D2/D3 (Nr4a3) are further magnified and visualized on the right of respective panels.

E. Visualization of ATAC-sequencing peaks around the activity-dependent DEG-ARGs *Stk40* (top) and *Grin1* (bottom) via the IGV genome browser with specific regions (E1 and E2) magnified on the right.

F. ATAC-sequencing peaks around the activity-dependent DEG-ARGs *Fos* (left-top), *Npas4* (right-top) and *Egr1* (bottom) via the IGV genome browser with specific regions (F1 and F2) magnified on the right.

G. (Top) Boxplots quantifying the normalized count values for accessibility within the consenus peaks annotated near respective ARGs. (Bottom) Boxplots quantifying the normalized count values for H3K27ac within the consenus peaks annotated near respective ARGs.

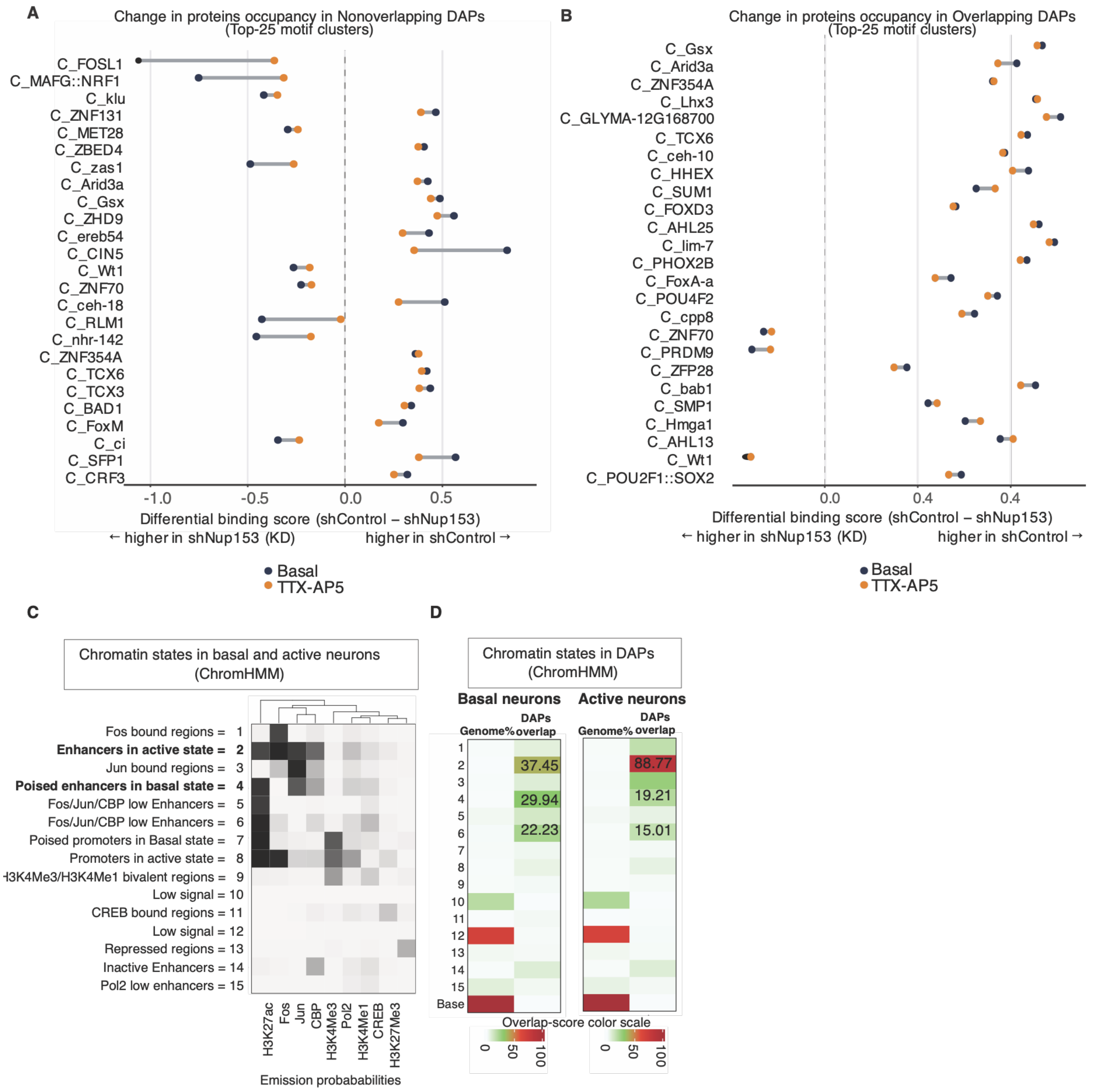


### Supplementary data associated with the main Fig. 4 & Table S2

### Fig. S9. Nup153-directed, basal-activity-dependent and independent regulation of chromatin organization

A. Visualization of the top 25 motif clusters with differential occupancy upon Nup153 depletion in Non-overlapping DAPs.

B. Visualization of the top 25 motif clusters with differential occupancy upon Nup153 depletion in overlapping DAPs.

C. The panel displays a heatmap of the emission parameters, with each column representing a chromatin state specific to either basal or active neurons. Each row corresponds to a transcription factor or a histone modification that was used to train ChromHMM with 15 chroamtin states (*33*). The scale reflects the probability of the molecular factors being present or absent in the corresponding state.

D. The heatmap shows the overlap enrichment of DAPs with ChromHMM states in either basal or active neurons. The scale reflects the proportion of DAPs that are annotated as the corresponding ChromHMM states in each cellular state.

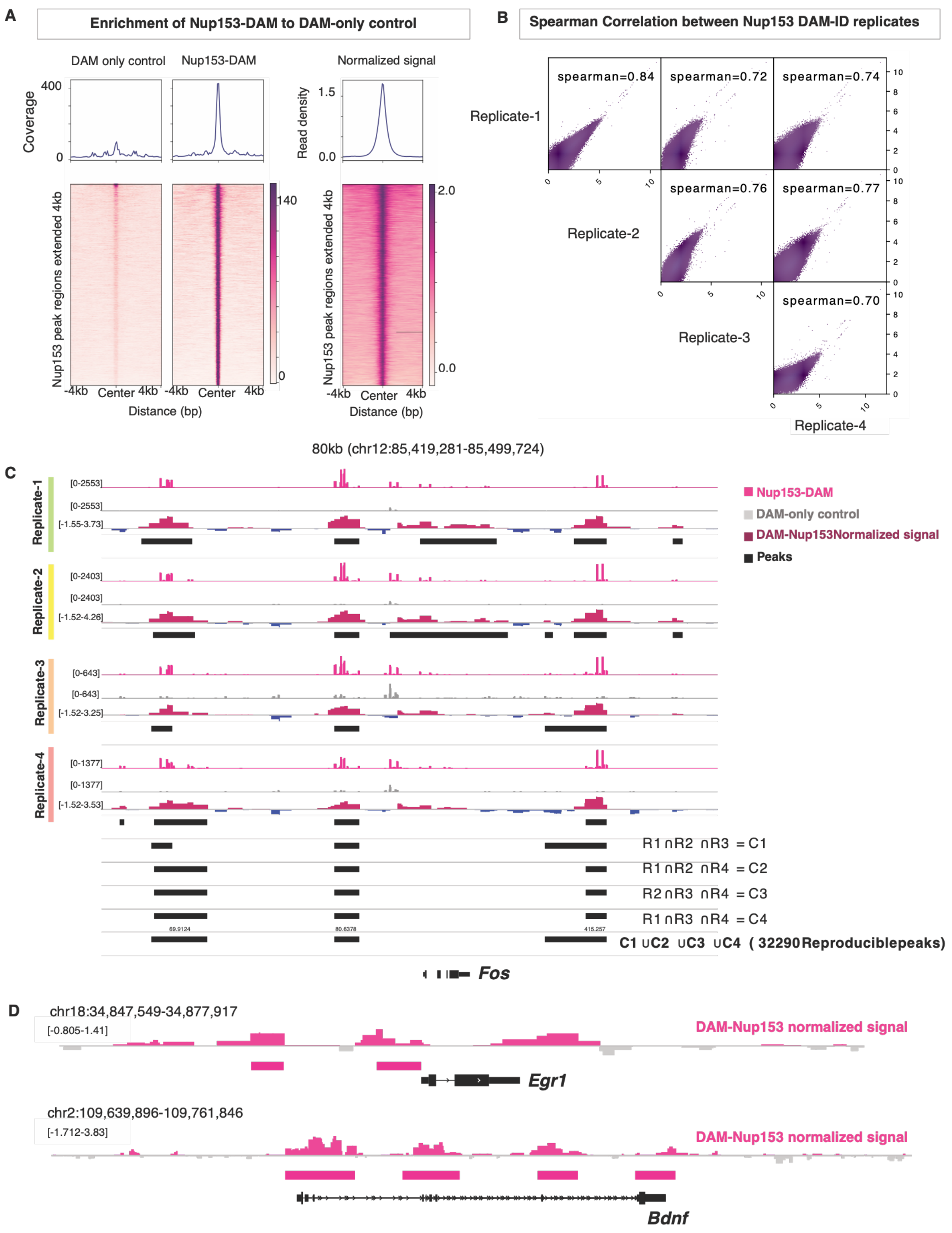


**Supplementary data associated with the main Fig. 5 & Table S3**
**Fig. S10. Nup153-associated chromatin regions in basal neurons**

(A) Enrichment of reads within the reproducible Nup153 peaks. The heatmap indicated the specificity of the DAM-Nup153 signal compared to the DAM-only control in the called peaks regions.

(B) High concordance of DAM-Nup153 binding across the genome. Spearman correlation between each of the four DAM-Nup153 replicates is visualized.

(C) Visualization of DAM-Nup153 (pink), DAM-only control signal (grey) alongside DAM-Nup153 normalized signal (magenta for positive signal and blue for negative signal) for each of the four biological replicates via IGV genome browser. C1-C4 depicts intersecting peaks in different combinations of three replicates. Final reproducible peaks were union of peaks in C1-C4.

(D) Visualization of normalized DAM-Nup153 tracks (pink) around the *Egr1 and Bdnf* genes in the basal state via the IGV genome browser. The reproducible DAM-Nup153 peaks are highlighted with pink boxes underneath the IGV tracks.

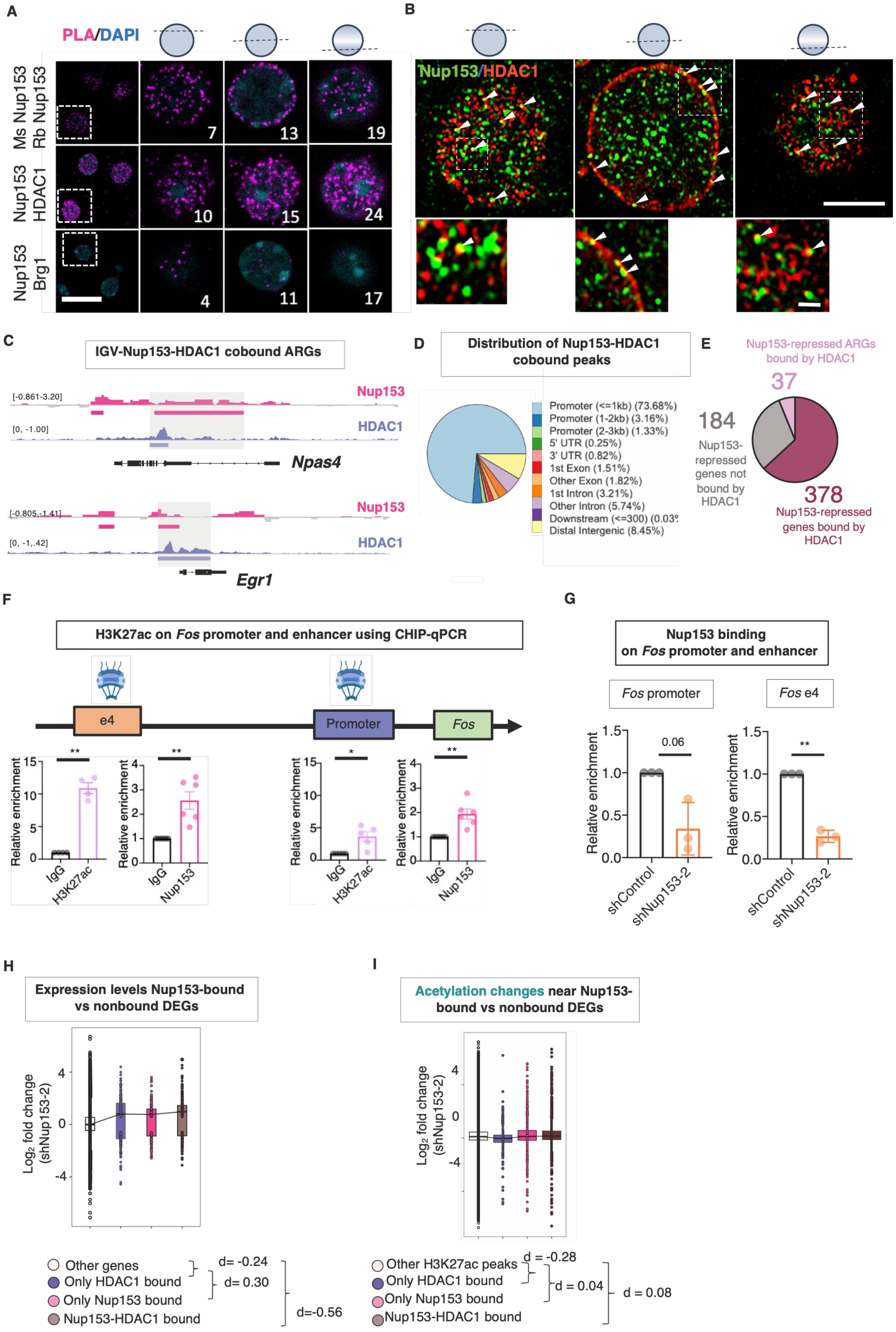
A
PLA/DAPI
Ms Nup153 Rb Nup153
Nup153 HDAC1
Nup153 Brg1
7
13
19
10
15
24
4
11
17
B
Nup153/HDAC1
C
IGV-Nup153-HDAC1 cobound ARGs
Nup153
HDAC1
Npas4
Egr1
D
Distribution of Nup153-HDAC1 cobound peaks
Promoter (<=1kb) (73.68%)
Promoter (1-2kb) (3.16%)
Promoter (2-3kb) (1.33%)
5' UTR (0.25%)
3' UTR (0.82%)
1st Exon (1.51%)
Other Exon (1.82%)
1st Intron (3.21%)
Other Intron (5.74%)
Distal Intergenic (8.45%)
E
Nup153-repressed ARGs bound by HDAC1
37
184
Nup153-repressed genes not bound by HDAC1
378
Nup153-repressed genes bound by HDAC1
F
H3K27ac on Fos promoter and enhancer using CHIP-qPCR
e4
Promoter
Fos
Relative enrichment
IgG
H3K27ac
Nup153
G
Nup153 binding on Fos promoter and enhancer
Fos promoter
Fos e4
0.06
shControl
shNup153-2
H
Expression levels Nup153-bound vs nonbound DEGs
Log2 fold change (shNup153-2)
Other genes
Only HDAC1 bound
Only Nup153 bound
Nup153-HDAC1 bound
d= -0.24
d= 0.30
d=-0.56
I
Acetylation changes near Nup153-bound vs nonbound DEGs
Other H3K27ac peaks
Only HDAC1 bound
Only Nup153 bound
Nup153-HDAC1 bound
d = -0.28
d = 0.04
d = 0.08

**Supplementary data associated with the main Fig. 6 & Table S4, S5**

**Fig. S11. Molecular mechanisms underlying the repression of ARGs by Nup153**

(A) *In situ* interactions between Nup153 and different epigenetic repressors revealed by PLA. Cells in the insets are magnified in the right panels. A series of Z-stack confocal image sections from the same cell are shown. The numbers in the panel indicate the position of the Z-stack images. Schema shows Z-location with respect to neuronal nuclei for Z-stack image below. Scale bars = 25 µm (left) and 6 µm(right).

(B) Visualizing co-localization of HDAC1 (green) with Nup153 (red). Confocal (airyscan) images of HDAC1 (green) and Nup153 (red) in PCNs were taken. The insets are magnified in the panels below. Arrows are highlight co-localization of Nup153 and HDAC1. Schema shows Z-location within the nuclei for image below. Scale bars = 5 µm (top) and 1 µm (bottom).

(C) Visualization of ChIP-seq peaks for HDAC1 (purple) and Nup153 (pink) around the *Npas4* and *Egr1* genes in PCNs using the IGV genome browser. Grey boxes highlight the overlapping peaks.

(D) Genomic distribution of Nup153-HDAC1 co-bound peaks in PCNs.

(E) A pie chart illustrates the number of Nup153-repressed genes associated with HDAC1 in PCNs. Refer also to Table S6.

(F) Quantification of Nup153 binding and H3K27ac levels with ChIP-qPCR on the promoter and e4 enhancer of the *Fos* gene in basal PCNs (**$p < 0.01$, one-sample *t*-test, N = 3 independent experiments).

(G) Quantification of Nup153 binding with ChIP-qPCR revealed a reduction of Nup153 binding on the promoter and e4 enhancer of the *Fos* gene in PCNs treated with shNup153-2 compared to shControl (**$p < 0.01$, one-sample *t*-test, N = 3 independent experiments).

(H) Boxplot showing $\log_2$ fold change in Nup153-bound DEGs, HDAC1-bound DEGs, and Nup153-HDAC1-bound DEGs compared to all other DEGs (Other genes vs Nup153-HDAC1 bound DEGs, $p < 2.2e-16$ Wilcoxon rank sum test; Cohen's d = 0.56, 95% CI [0.46, 0.67] ; Other genes vs only Nup153 bound DEGs, p = 0.0007, Wilcoxon Rank sum test; Cohen's d = 0.30, 95% CI [0.17, 0.42] ; Other genes vs HDAC1 bound DEGs, p = 0.0006, Wilcoxon Rank sum test; Cohen's d = 0.24, 95% CI [0.09, 0.38]).

(I) A boxplot showing $\log_2$ fold changes in histone acetylation near Nup153-DEGs bound by only Nup153, only HDAC1, both Nup153 and HDAC1, or neither of the proteins (Other H3K27ac peaks vs Nup153-HDAC1 bound peaks, $p < 4.31e-07$, Wilcoxon Rank sum test; Cohen's d = 0.08, 95% CI [0.04, 0.12]).

The data (Fig. S9F & S9G) are presented as mean ± standard deviation (s.d). Data for Fig. S9F are presented as a boxlplot with a connecting line to the median of each data group.

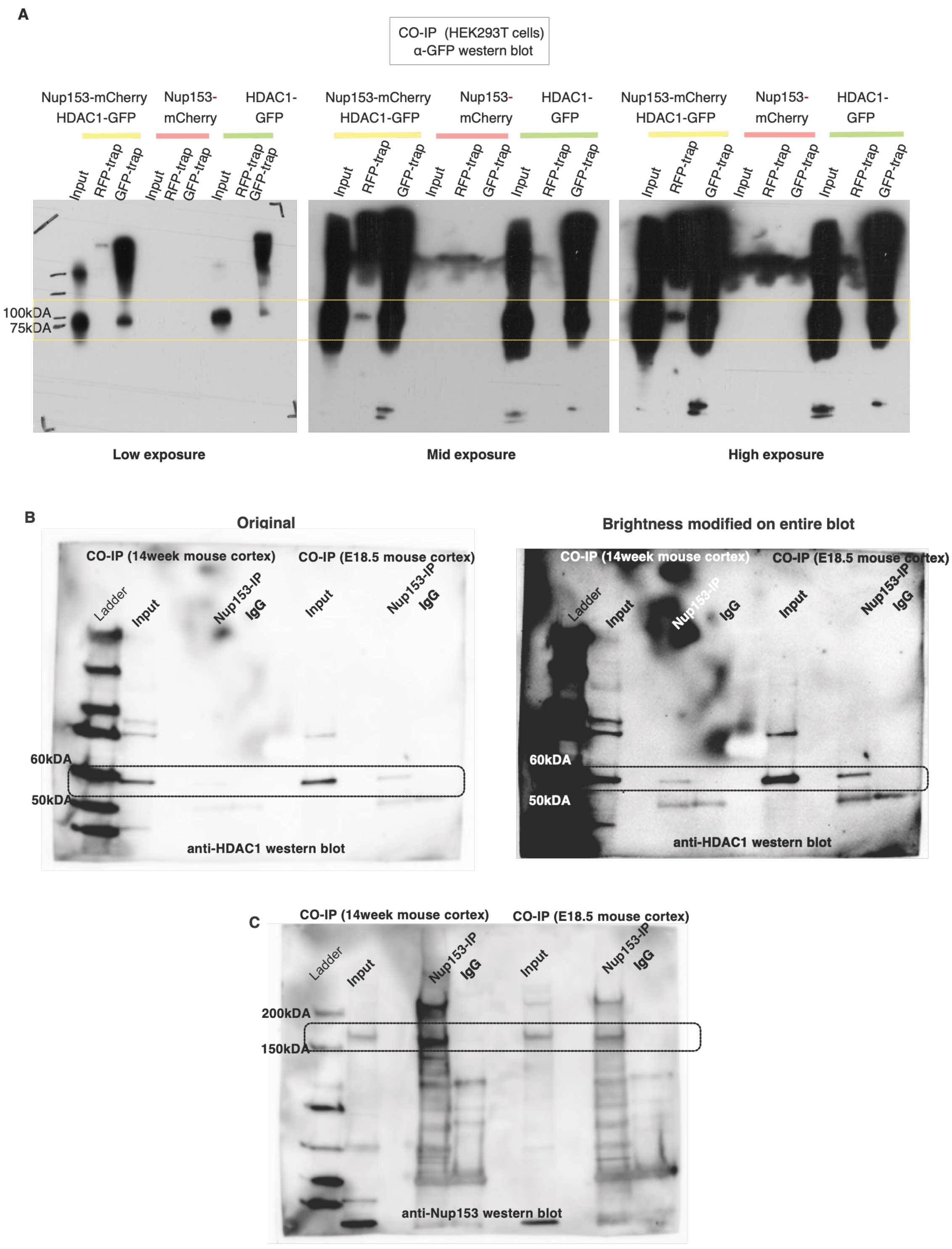


**Supplementary data associated with the main Fig. 6**

**Fig S12. Coimmunoprecipitation for Nup153 and HDAC1.**

**(A)** Immunoblots with CO-IP samples depict positive interaction between Nup153-mCherry and HDAC1-GFP exogenously expressed in HEK293T cells. Same blots are presented with three different exposures. HDAC-GFP is co-immunoprecipitated along with

Nup153-mCherry using RFP-trap. However, we do not see immunoprecipitation of HDAC1-GFP in absence of Nup153-mCherry via RFP-trap.

(B) Co-immunoprecipitation revealed that HDAC1 interacts with Nup153 in the embryonic mouse cortex (E18.5) and adult mouse (14 week). Dotted box highlight HDAC1 co-immunoprecipitated with Nup153 in comparison to IgG control.

**(C)** Blot in (B) was modified via changing brightness and contrast setting on the entire blot to improve visibility of HDAC1 immunoprecipitation in adult mouse cortex.

**(D)** Western blot for Nup153 on the same blot as in (B) and (C) to confirm presence of Nup153, confirming HDAC1 immunoprecipitation specifically with Nup153

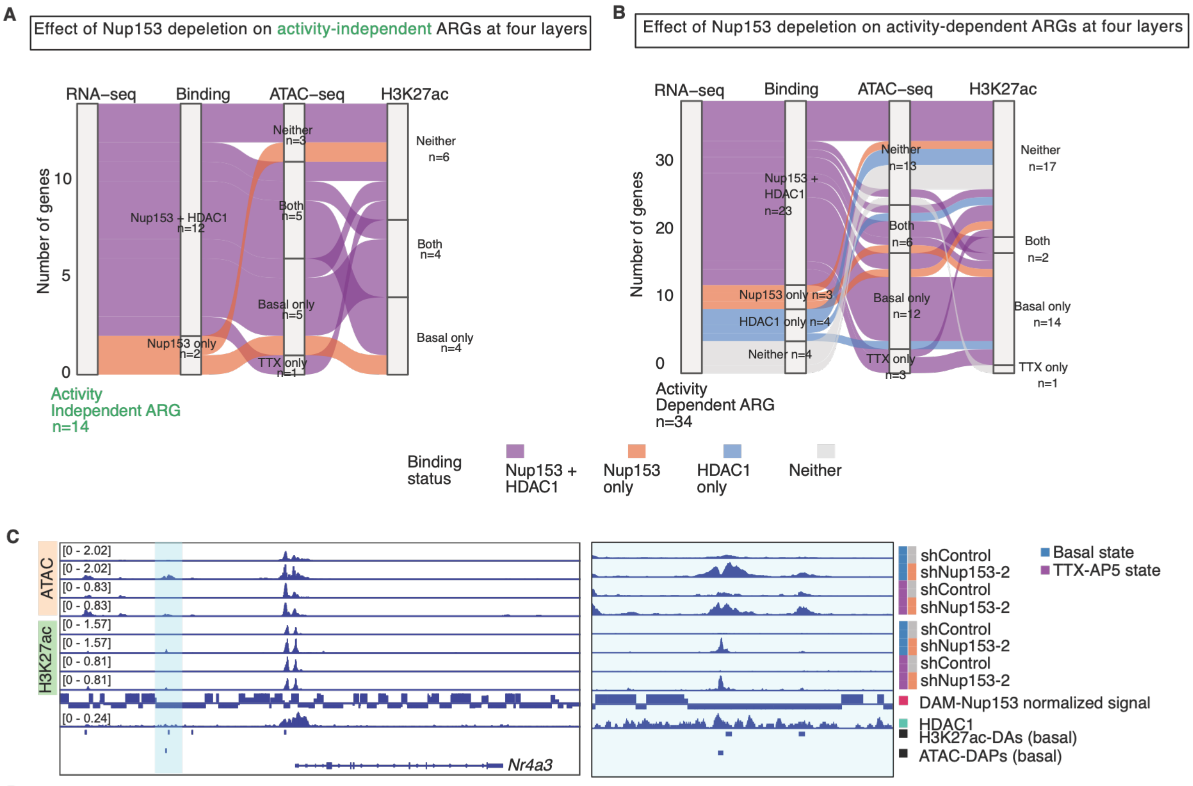


**Supplementary data associated with the main Fig. 6 & Table S6**

**Fig. S13. Nup153-HDAC1-mediated histone acetylation changes near ARGs**

A. A alluvial plot showing the connection of activity-dependent ARGs with different epigenomic layers and it's relation to Nup153 depletion in the basal TTX-AP5 treated conditions.

B. A alluvial plot showing the connection of activity-independent ARGs with different epigenomic layers and it's relation to Nup153 depletion in the basal and TTX-AP5 treated conditions.

C. (Left) A IGV plot visualizing the *Nr4a3* gene and associated changes at ATAC and H3K27ac levels in the basal and TTX-AP5 treated states. (Right) Magnified view of blue region highlighted in (Left).

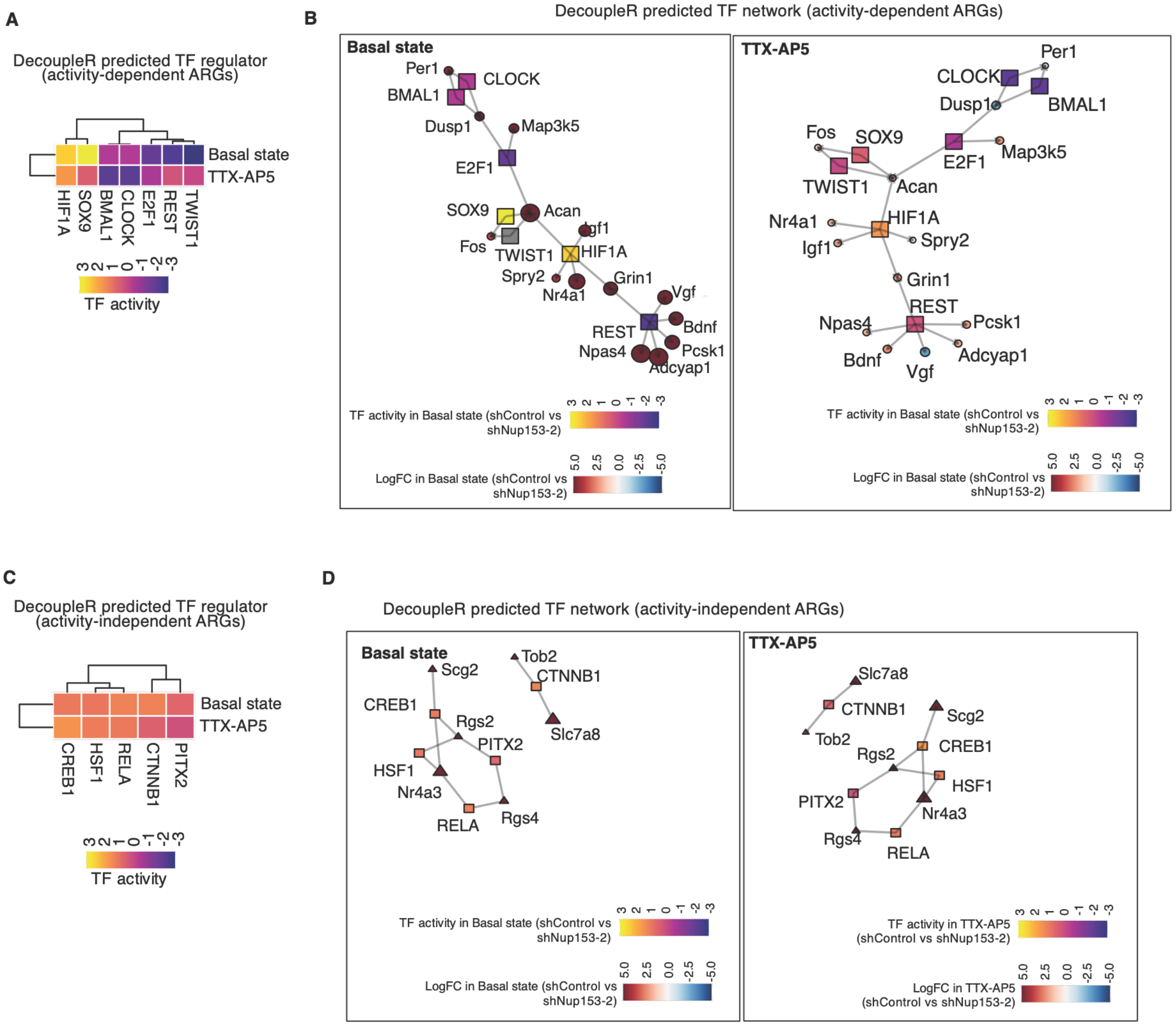


**Fig. S14. DecoupleR (*88*) based analysis of transcription factor activity**

A. A heatmap visualizing DecoupleR-derived transcription factor (TF) activity for the top regulators of basal-activity dependent ARGs. The scale shows the range of transcription factor activity values.
B. Network plot visualizing the top TF regulators (squares) connected to their target basal activity-dependent ARGs (circle) in the basal or TTX-AP5 treated conditions. The top scale shows the range of TF activity, and the bottom scale shows the range of $\log_2$ fold change in samples with Nup153 depletion in the basal state (left bottom scale) and the TTX-AP5 treated condition (right-bottom scale).
C. A heatmap visualizing DecoupleR derived transcription factor (TF) activity for top regulators of basal activity-independent ARGs. The scale shows the range of transcription factor activity values.
D. Network plot visualizing top TF regulators (square) connected with their target basal activity-independent ARGs (circle) in the basal or TTX-AP5 treated conditions. The top scale shows the range of TF activity, and the bottom scale shows the range of $\log_2$ fold

change in samples with Nup153 depletion in the basal (left bottom scale) and TTX-AP5 treated condition (right-bottom scale).
